# Reshaping of the *Arabidopsis thaliana* proteome landscape and co-regulation of proteins in development and immunity

**DOI:** 10.1101/2020.03.09.978627

**Authors:** Mona Bassal, Petra Majovsky, Domenika Thieme, Tobias Herr, Mohammad Abukhalaf, Mohamed Ayash, MHD Rami Al Shweiki, Carsten Proksch, Ali Hmedat, Jörg Ziegler, Steffan Neumann, Wolfgang Hoehenwarter

**Author notes:** Address correspondence to Wolfgang Hoehenwarter, Proteome Biology of Plant Interactions Research Group, Leibniz Institute of Plant Biochemistry.

## Abstract

Proteome remodeling is a fundamental adaptive response and proteins in complex and functionally related proteins are often co-expressed. Using a deep sampling strategy we define *Arabidopsis thaliana* tissue core proteomes at around 10,000 proteins per tissue and absolutely quantify (copy numbers per cell) nearly 16,000 proteins throughout the plant lifecycle. A proteome wide survey of global post translational modification revealed amino acid exchanges pointing to potential conservation of translational infidelity in eukaryotes. Correlation analysis of protein abundance uncovered potentially new tissue and age specific roles of entire signaling modules regulating transcription in photosynthesis, seed development and senescence and abscission. Among others, the data suggest a potential function of RD26 and other NAC transcription factors in seed development related to desiccation tolerance as well as a possible function of Cysteine-rich Receptor-like Kinases (CRKs) as ROS sensors in senescence. All of the components of ribosome biogenesis factor (RBF) complexes were co-expressed tissue and age specifically indicating functional promiscuity in the assembly of these little described protein complexes in Arabidopsis. Treatment of seedlings with flg22 for 16 hours allowed us to characterize proteome architecture in basal immunity in detail. The results were complemented with parallel reaction monitoring (PRM) targeted proteomics, phytohormone, amino acid and transcript measurements. We obtained strong evidence of suppression of jasmonate (JA) and JA-Ile levels by deconjugation and hydroxylation via IAA-ALA RESISTANT3 (IAR3) and JASMONATE-INDUCED OXYGENASE 2 (JOX2) under the control of JASMONATE INSENSITIVE 1 (MYC2). This previously unknown regulatory switch is another part of the puzzle of the as yet understudied role of JA in pattern triggered immunity. The extensive coverage of the *Arabidopsis* proteome in various biological scenarios presents a rich resource to plant biologists that we make available to the community.

## Introduction

Proteome biology is receiving increasing interest in the last years but still proves difficult on a genome wide scale in plants. The proteome is the most fundamental active determinant of an organism’s phenotype and its landscape is large, complex and dynamic, entailing changes in protein abundance, interaction, post translational modification (PTM) and sub-cellular localization.

Steady state protein abundance at a certain time point is to a considerable part determined by the abundance of its transcript and the latter’s translation rate. Synthesis is however only half of the equation governing protein abundance, indeed *Arabidopsis* has more than 600 F-box proteins as components of diverse E3 ubiquitin ligase complexes that direct protein degradation. Newer evidence has shown that post transcriptional and translational mechanisms (Ponnala et al., 2014; Merchante et al., 2017) and phenomena such as cell-to cell mobile mRNAs (Thieme et al., 2015) and proteins (Han et al., 2014; Guan et al., 2017) play equally important roles in determining the proteome’s temporal and spatial plasticity foremost in steady state shifts. Therefore, despite the continued practice of quantifying the abundance of proteins’ cognate transcripts to estimate and quantify functional protein coding gene expression, direct, large scale measurement of protein abundance and PTM should be the explicit end point of functional genomics.

The problem with this is that proteomics has long been beset by a lack of sensitivity, especially in plants. Indeed, only a handful of true deep proteomics studies in this kingdom can be found. Walley and co-workers constructed a protein co-expression network based on measurement of 17,862 proteins and 6,227 phosphorylated proteins in maize (Walley et al., 2016). The tissue and development specific wheat proteome has been mapped with measurement of 15,779 proteins (Duncan et al., 2017) as well as the tomato fruit where 7,738 proteins were measured in one or more developmental stages (Szymanski et al., 2017). Song and co-workers reported optimized FASP sample preparation in conjunction with 2D-LC-MS/MS allowed measurement of 11,690 proteins from a single *Arabidopsis* leaf sample (Song et al., 2018) and Baerenfaller and co-workers reported measurement of nearly 15,000 *Arabidopsis* proteins more than ten years ago (Baerenfaller et al., 2008).

Here we go beyond this classic study and elucidate *Arabidopsis thaliana* tissue specific proteome remodeling in development and immunity in unprecedented resolution and detail. We describe a procedure that allows deep sampling of up to 9,000 proteins per plant sample and does not require the expertise necessary for 2D-LC-MS. We quantified nearly 16,000 proteins in absolute terms (copy number per cell) and conducted a global analysis of protein PTM.

A long standing caveat in proteomics is proteomics “dark matter” (Skinner and Kelleher, 2015) meaning an abundance of high quality MS2 spectra that do not result in PSM. Many of these MS2 spectra are derived from peptides bearing PTM, however classic search algorithms necessitate PTM predefinition, limiting potential PTM identification to a handful. Recent years have seen the advent of “open search” algorithms (Chick et al., 2015; Kong et al., 2017; Bagwan et al., 2018; Chi et al., 2018) that allow unrestricted precursor mass shifts in PSM, and so have the potential to identify the vast array of biologically occurring modifications on a proteome wide scale. Here we performed such a survey using the MSFragger (Kong et al., 2017) software suite in *Arabidopsis*.

Large scale protein co-expression analysis is a potentially powerful strategy to determine functional relationships between proteins because it circumvents the limitations inherent to transcriptomics measurements described above. We applied clustering algorithms to our data set that provides extensive coverage of the *Arabidopsis* proteome to uncover tissue and developmentally specific protein expression patterns. The approach was effective in pinpointing co-regulation of all components of protein complexes and developmentally timed signaling modules. More importantly, it then allowed inference of previously unknown functions of proteins, protein families and entire signaling modules based on the same expression patterns. This included processes such as tissue and age specific ribosome biogenesis, photosynthesis, ABA signaling and NAC transcription factors in seed development and establishment of dormancy and senescence and abscission.

The various sequential steps of ribosome biogenesis including the involved RPs and RBFs are well described in yeast and human (Henras et al., 2008; Henras et al., 2015). Mutations in numerous RPs and RBFs are known to cause severe developmental defects, so called ribosomopathies in humans. This also holds true in *Arabidopsis* where a number of mutations in RBFs affect gametophyte development and embryogenesis (Byrne, 2009; Weis et al., 2015). Furthermore *Arabidopsis* ribosomes are extensively heterogenic, each individual RP being encoded by two to seven paralogs (Weis et al., 2015). This heterogeneity of ribosome species is dependent on developmental stage, tissue and environmental stimuli, suggesting that the specific ribosome constituency may play a regulatory role in these processes.

Cotyledons of young seedlings are characterized by light induced chloroplast biogenesis. Photomorphogenesis includes tetrapyrrole and chlorophyll biosynthesis to establish photosystems along with carotenoid synthesis as an accessory pigment and ROS scavenger. Once photoautotrophic, protein biosynthesis is ramped up, hallmarked by the increased abundance of ribosomal proteins and proteins involved in translation and folding. Concomitantly, nuclear encoded proteins synthesized in the cytosol are imported. Finally mature chloroplasts with their established structures proliferate by division to accommodate cell expansion and division in the growing leaves.

Senescence is a coordinated process with several stages and developmental check points (Bleecker and Patterson, 1997; Rogers and Munne-Bosch, 2016). Early senescence syndrome involves reprogramming of gene expression (Breeze et al., 2011) and redox and ROS signaling. This is followed by ordered dismantling of the photosynthetic apparatus leading to ROS production and involving ROS control and nutrient remobilization to other plant parts in the case of leaves and to the developing ovary in the case of floral petals. The final result is dell death lastly followed by abscission in some organs such as floral petals.

Several post-transcriptional/translational mechanisms controlling protein abundance have just recently been uncovered in plant immunity (Meteignier et al., 2017; Xu et al., 2017; Tabassum et al., 2019). Therefore we investigated reshaping of the proteome in the steady state shift from growth to pattern triggered immunity (PTI). We measured changes in protein abundance of more than 2,000 proteins in all avenues of PTI as well as photosynthesis and primary metabolism, giving a comprehensive picture of altered proteome architecture in basal immunity. The focus was set on hormone signaling and the deep proteomics measurements were complemented by targeted parallel reaction monitoring (PRM) proteomics, qPCR and phytohormone and amino acid measurements. Salicylic acid (SA) and Jasmonate (JA) are the phytohormones quintessentially associated with plant defense, the former constituting the backbone of resistance to biotrophic pathogens, the latter to necrotrophic pathogens as well as wounding (Pieterse et al., 2012). There mutual antagonism to prioritize one defense strategy over the other as needed is a long standing paradigm in the field of plant immunity. However research in the last decade has made it clear, that JA also plays a role in resistance to biotrophs and pattern triggered immunity (PTI) which is however not yet fully understood (Nickstadt et al., 2004; Hillmer et al., 2017; Mine et al., 2017). The integrative –omics strategy applied here brought to light a previously unknown regulatory switch wherein a MYC2 dependent negative feedback loop controls JA-Ile and JA levels via deconjugation and hydroxylation by IAR3 and JOX2 respectively. It reconciles the well-known dampening of JA levels and suppression of JA signaling downstream of JA synthesis by SA, adding another feature to the picture of JA activity in PTI.

Finally we make both the raw and meta data produced in this study available to the general public (submission to Proteome Exchange and TAIR pending). The sampling depth of the proteome in development and immunity should prove to be a valuable resource to plant biologists.

## Results

### Deep proteomics method

We conducted discovery proteomics measurements of several *Arabidopsis thaliana* Col-0 tissues throughout the lifetime of the plant. These were roots, leaves, cauline leaves, stem, flowers and siliques/seeds as well as whole plant seedlings as early as seven days up to 93 days of age when the plant was in late senescence (Supplemental figure 1 and Supplemental methods and data tables 1 and 2). In addition we measured proteomes of PTI elicited plants treated with the peptide flg22. Primarily we were interested in reshaping of proteome architecture in these different biological scenarios to capture protein co-regulation and predict tissue and developmentally specific protein function.

To this end it was essential to develop a method that allows deep sampling of the plant proteome in a reasonable time frame. It was clear that this would have to begin with comprehensive extraction of tissue proteins and further entail multi-step fractionation of the complex extract. We settled on and optimized 4% SDS protein extraction and Gel-LC MS combining SDS-PAGE protein separation and reverse phase (RP) LC peptide separation on-line with ESI MS (see Supplementary methods and data and Supplemental figure 2 for full method and optimization details).

Plant tissue is more recalcitrant to proteomics analysis than other samples. The plant cell wall requires harsh disruption techniques. Plant tissue contains an abundance of secondary metabolites, oils and waxes and more significantly, in the case of green tissue, pigments as part of the light harvesting complexes, that all interfere with LC-MS. Green tissue also contains the most abundant protein on earth, Ribulose bisphosphate carboxylase (RuBisCo) leading to suppression of less prominent ion signals in the mass analyzer and severely hampering detection of less abundant proteins.

SDS-PAGE alleviated all of these issues (Supplemental figure 3 A): i) It allowed the depletion of high amounts of SDS used for protein extraction. ii) Pigments and small molecules conglomerated in a green low molecular weight band below the protein front effectively partitioning them from the proteins. iii) RuBisCo large and small subunits (RBCL and RBCS) migrated to two prominent bands facilitating their separation from the rest of the proteome.

Five gel slices were in gel digested with trypsin and individually injected into the LC-MS neatly fractionating the proteome according to the molecular weight of its constituents (Supplemental figure 3 B). The most abundant plant proteins including RBCL and RBCS were separated in individual fractions in both leaves and roots, diminishing the suppressive effects of over abundant proteins on peptide and protein identification in single shot LC-MS measurement of the entire proteome (Supplemental figure 3 C and D). This allowed the identification of between 5977 and 9524 protein groups per sample, each identified with at least 1 unique peptide at protein and peptide FDR thresholds of 1%. These protein groups will henceforth be referred to as proteins (Supplemental table 1 and Supplemental figure 3 E). Measurements of the individual proteomes showed intra sample variability that was considerably lower that the observed inter sample variability (Supplemental figure 3 F and G).

### The *Arabidopsis* proteome

In the entire study we identified and quantified 15926 *Arabidopsis thaliana* proteins in total, 15845 encoded in the nuclear genome (Supplemental table 2). For further details on the parameters of the entire data set see Supplemental methods and data Table 3. Nuclear protein identifications were evenly distributed over all five chromosomes. We complemented the proteomics data with transcriptomics data from an extensive study of *Arabidopsis* development using microarrays that measured gene expression on 22157 nuclear loci including 1058 non-protein coding genes (Schmid et al., 2005). Our proteomics data presents mass spectrometric evidence for cognate protein expression of 60% (and locally more) of the *Arabidopsis* nuclear genome and close to 70% or more of the transcriptome, including 1886 proteins for which no transcripts were identified or which were not on the microarray (Figure 1 A and B).

**Figure 1.**
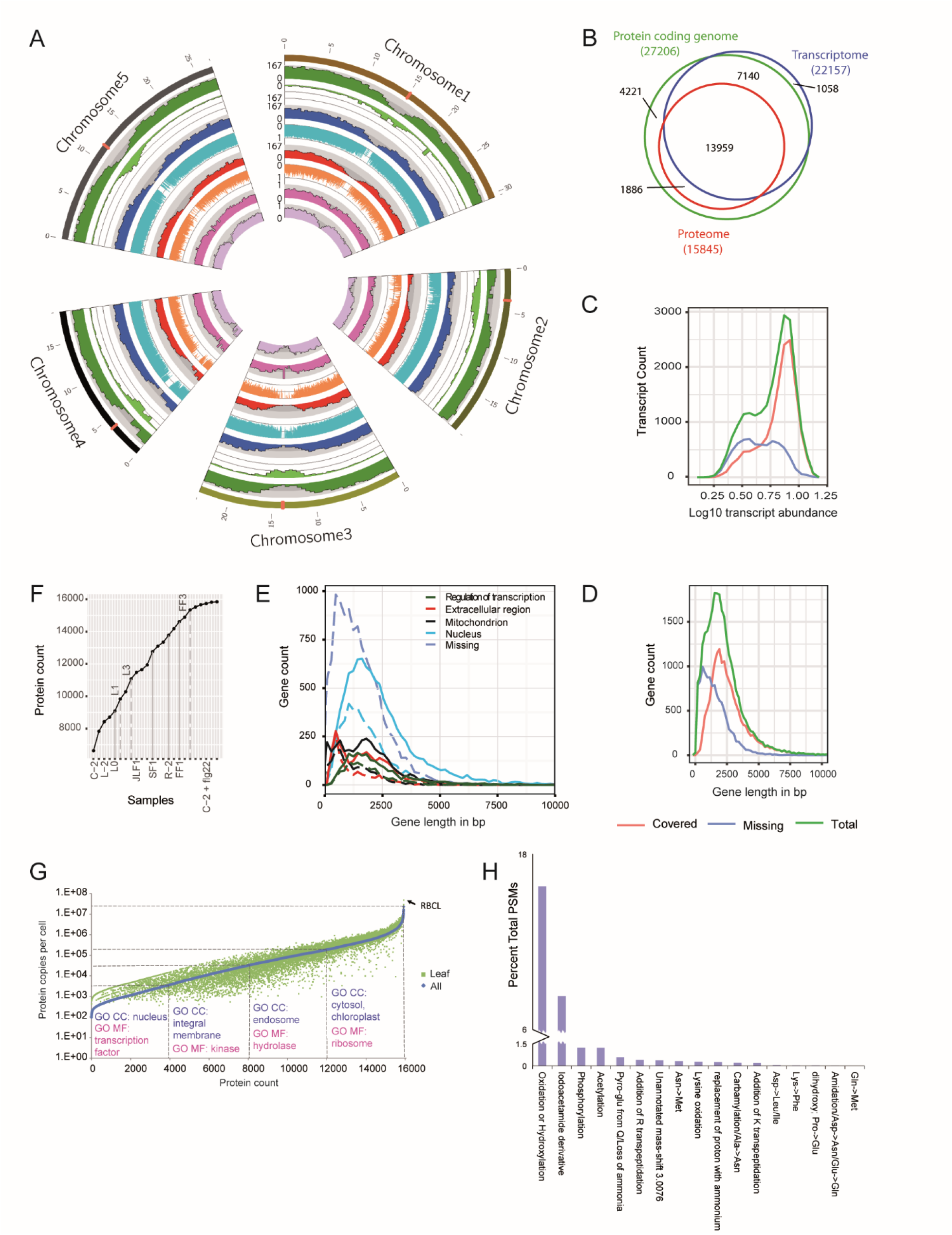
Deep coverage of the Arabidopsis thaliana proteome. A. Mapping protein expression to the 5 Arabidopsis nuclear chromosomes (scale in Megabases (Mb), centromers are indicated). Tracks from outside to inside: Number of protein coding genes per 500 Kb; scale 0 to 167 (maximum) [dark green]. Number of non-protein coding genes per 500 Kb; scale 0 to 167 [light green]. Number of transcripts per 500 Kb; scale 0 to 167 [dark blue]; Log10 transcript abundance normalized to maximum transcript abundance; scale 0 to 1 [light blue]. Number of proteins per 500 Kb; scale 0 to 167 [dark red]. Log10 protein abundance normalized to maximum protein abundance; scale 0 to 1 [light red]. Fraction of protein coding genes for which cognate proteins were detected; scale 0 to 1 [magenta]. Fraction of transcripts for which cognate proteins were detected; scale 0 to 1 [light purple]. B. Scaled VENN diagram showing the coverage of protein coding genes by cognate transcripts and proteins. C. Relationship between transcript abundance and detection of its cognate protein. Green: all protein coding transcripts, red: protein coding transcripts with a detected cognate protein, blue: protein coding transcripts without a detected cognate protein. Frequency polygon drawn with 20 bins D. Relationship between protein coding gene size in bp and detection of its cognate protein. Green: all protein coding genes, red: protein coding genes with a detected cognate protein, blue: protein coding genes without a detected cognate protein. Frequency polygon drawn with bin size 190 bp. E. Size in bp of genes assigned to the gene ontology terms regulation of transcription, DNA templated (GO:0006355), extracellular region (GO:0005576), mitochondrion (GO:0005739) and nucleus (GO:0005634). Solid lines indicate all members of the GO term, dashed lines indicate members of the GO term not identified in the proteomics measurements, i.e. the missing proteome. Gene size of all proteins not identified in the proteomics measurements, the entire missing proteome is also indicated. Frequency polygon drawn with 350 bins and bin size 190 bp. F. Cumulative increase in identified proteins, i.e. proteome coverage, as different tissues and flg22 treatment are sampled. G Copy numbers per cell of all identified proteins and leaf proteins as determined by proteomics ruler method (Wisniewski et al., 2014). Quartiles are indicated by dashed lines. Significantly enriched GO annotations by DAVID are shown for the individual quartiles. H Global survey of PTM in the Arabidopsis thaliana proteome. The relative abundance PTMs comprising 0.1% or more of the total PSMs in at least 1 tissue type or biological scenario (flg22 treatment) are shown.

We then asked what precluded more extensive coverage of the *Arabidopsis* proteome, i.e. the missing proteome. We first looked at a potential impact of transcript abundance on the detection of cognate proteins and found that missing (not detected) proteins are evenly distributed over the range of transcript abundance with the exception of the most abundant transcripts for nearly all of which a cognate protein was detected (Figure 1 C). This indicates that transcript abundance is not the primary factor impeding protein detection. Next we investigated if gene length in base pairs as a proxy for protein length may have an impact and found that indeed the distribution of missing proteins was shifted markedly towards smaller genes from the distribution of detected proteins (Figure 1 D). This suggests that protein size is a factor limiting protein detection. This effect could be observed for several gene ontology categories that were overrepresented in the missing proteome (Supplemental figure 4). Then we examined how tissue specific protein expression contributed to the cumulative expression of the entire proteome (Figure 1 F). While every tissue contributed to cumulative protein expression curve with a steep slope, it can also be seen that the curve converges on saturation as sample numbers increase. This was also the case for flg22 treated tissues sampled last, where one could expect expression of a host of immunity specific proteins not detected in normally developing tissue. This suggests that further sampling of *Arabidopsis* would not greatly increase coverage of the entire proteome.

We quantified all 15,927 measured proteins in terms of protein copy number per cell in the entire sample set and in leaf tissue (LF) by way of the proteomics ruler approach (Wisniewski et al., 2014). It equates the total histone MS signal (total number of histone PSMs in our case) to the cellular DNA mass allowing the conversion of PSMs to mass units and calculation of the total cellular protein mass. Individual to total protein PSM ratios can then be used to calculate individual protein copy numbers per cell.

The total cellular protein mass was calculated as 256.5 pg, in agreement with amounts from cell lines and tissues (Wisniewski et al., 2014). Protein copy numbers spanned a range of more than 5 orders of magnitude from just below 100 to 2.29e+07 copies per cell (Figure 1 G, Supplemental table 3). The most abundant protein was RBCL with 4.73e+07 cellular copies in leaves. The total copy number of all detected small subunits was 5.4e+07 so the ratio of large to small subunits was 0.876, close to a 1:1 ratio. Gene ontology analysis of the proteins in the four quartiles showed that nuclear proteins and regulators of transcription were very specifically among the least abundant proteins (1750 of 3970 1^st^ quartile proteins were annotated as nucleus). Membrane spanning proteins and kinases were generally also lower abundant, 1108 of 3970 2^nd^ quartile proteins being annotated as integral membrane.

Protein post translational modification (PTM) is an essential modulator of protein function. Therefore we performed a proteome wide survey of global PTM in Arabidopsis thaliana with our deep proteomics data set using the “open-search” algorithm MS-Fragger (Kong et al., 2017). This led to the identification of more than 3.5 million PSMs from more than 11.6 million total acquired MS2 spectra. The most abundant PTMs comprising more than 0.1% of total PSMs in at least one tissue type or biological scenario are shown in figure 1 H (for full data see Supplemental table 4). Next to predominantly experimentally induced PTM (protein oxidation and carbamidomethylation of cysteine residues to reduce disulfide bonds), serine or threonine phosphorylation and N-terminal acetylation were abundant naturally occurring modifications affecting approximately 1.25% of total protein abundance. Transpeptidation reaction, a non-translational mechanisms for the formation of peptide bonds, derived addition of amino acids was also detected. Furthermore, a number of amino acid substitutions in protein primary structure were common.

### Architecture of Tissue and Developmental Proteomes

We utilized our extensive MS data collection as a resource to investigate the individual tissue proteomes during the *Arabidopsis* life cycle and co-regulation of protein abundance between them (Supplemental table 5). First we grouped all of the data tissue specifically also merging rosette and cauline leaves and qualitatively compared the tissue proteomes which each comprised around 10,000 proteins (Supplemental table 6). Around 6,500 proteins (by far the largest set), were ubiquitous to all tissues whereas 500 to 600 proteins were unique to each tissue with exception of leaves which showed nearly 1,000 unique proteins, perhaps as a consequence of the larger number of aggregated leaf samples (Figure 2 A). Also around 1,000 proteins were absent in roots but present in all other tissues, reflecting the former tissues below ground nature.

**Figure 2.**
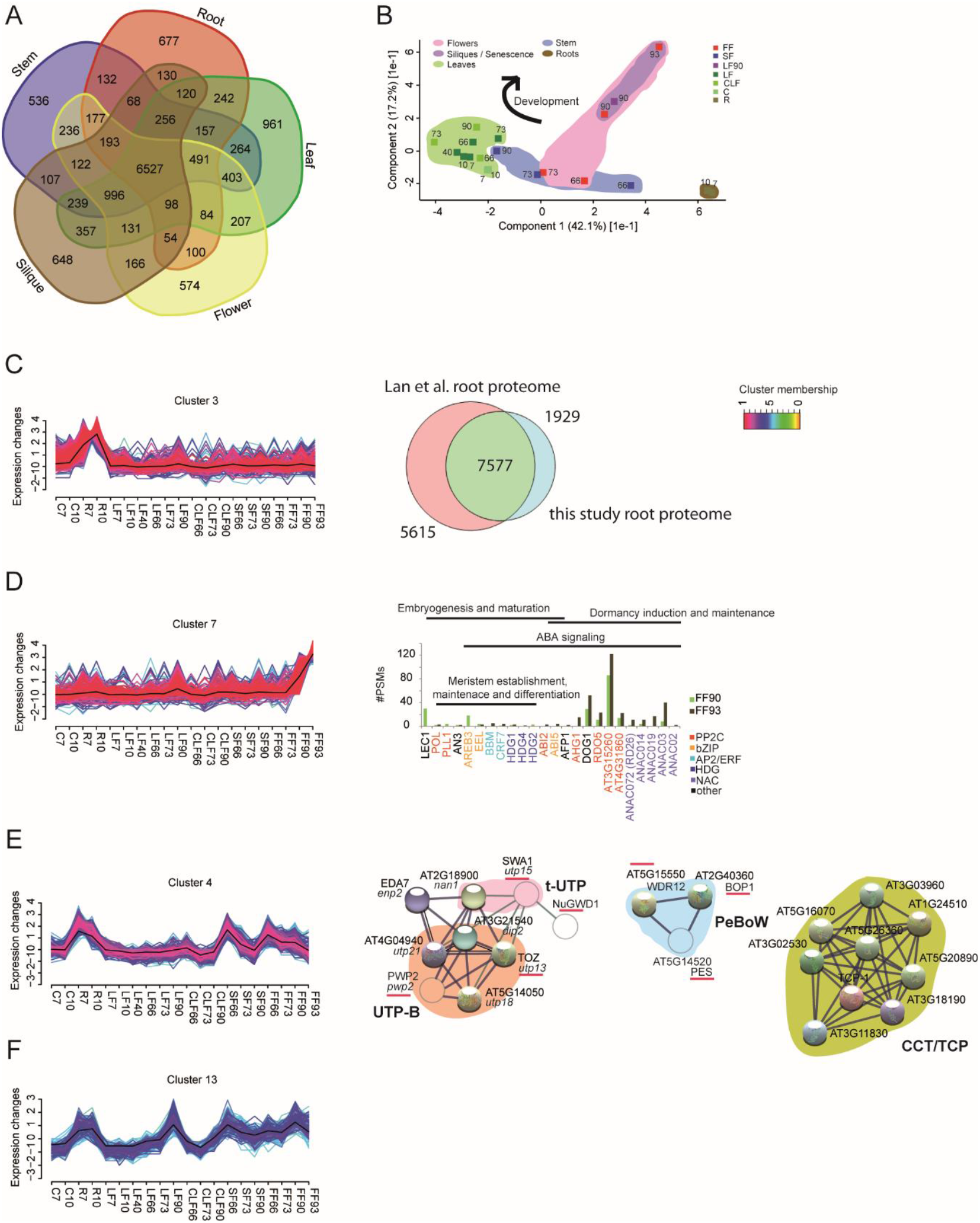
Tissue and development specific dynamic architecture of the Arabidopsis proteome. A. Qualitative comparison of Arabidopsis tissue proteomes. All measurements of a tissue regardless of plant age were aggregated, leaves and cauline leaves as well. B. Principal component analysis (PCA) of the deep proteomics measurements of sampled tissues. Numbers indicate sample, i.e. plant age in days. Tissues are color coded. Purple indicates late flowering stages, i.e. siliques and leaves in senescence. C. FCM cluster 3 showing proteins exclusively abundant in roots. Right panel: Qualitative comparison of all root proteins identified in this study and all root proteins identified in Lan et al. FCM core clusters were extracted wherein all clustered proteins have membership values α exceeding a threshold of 0.5 meaning cluster members all have similarity to the cluster centroid greater than 0.5 (Futschik and Carlisle, 2005). D. FCM cluster 7 showing proteins increasing in abundance in seed development and exclusively present in seeds or siliques. Right panel: Protein abundance of kinases and transcription factors (PQI given is raw #PSMs) in the respective samples. E. FCM cluster 4. Right panel: Physical protein interaction networks produced with STRING database based on homology to yeast and human studies. Solid spheres in the UTP-B, t-UTP and PeBoW networks are proteins designated as DWD40 by DAVID gene ontology analysis and part of the respective STRING input dataset (WD40/YVTN repeat-like-containing domain list Supplemental Table 9). Unfilled spheres are not part of the WD40 input set but are also members of cluster 4. Thin black edges indicate bona fide physical interactions identified in humans (Wan et al., 2015). Green edges indicate bona fide physical interactions identified in Arabidopsis (Ishida et al., 2016). Red underlines indicate deletion of the gene has a developmental phenotype. Yeast homologs of Arabidopsis gene names are given in italics. F. FCM cluster 13.

The large volume of the data made PCA an attractive method to reduce its dimensionality and explore the relationships of the sampled proteomes. The first two principal components accounted for nearly 50% of the total variance indicating that the linear projection in this two dimensional subspace reflects the predominant data structure (Figure 2 B). The individual tissue proteomes were clearly separated with the exception of flowers and the inflorescence stem at the same point in development (66 and 73 days). More interestingly, a developmental component was also visible in all sampled tissues, reflecting proteome remodeling during ageing (increasing sample age bottom right to top left; arrow). Hierarchical cluster analysis (HCL) corroborated the PCA results (Supplemental figure 5).

To investigate the dynamics of proteome structure and protein co-expression in tissue development in detail and thereby extrapolate protein function in previously undisclosed contexts, a noise robust soft partitioning technique that does not assign a feature exclusively to a single group, called fuzzy c-means clustering was applied (citation). The procedure produced 7 out of 16 clusters (clusters numbered 3, 4, 7, 8, 10, 11, 13) with biologically meaningful changes in protein abundance with a permutation based FDR of less than 1%.

### Root Proteome

The largest set of 577 proteins assigned to a cluster were root specific and not abundant or present in any other tissues at any of the sampled developmental stages (Cluster 3, Figure 2 C, Supplemental table 7). Gene ontology analysis of these proteins indicated that many of them were involved in processes related to the extracellular region, metal-binding and oxidation and reduction. We described the expression of these proteins in the context of phosphate metabolism previously (Hoehenwarter et al., 2016). We compared our results to an exhaustive study of the root proteome by Lan et al (Lan et al., 2012) and found that around 70% of the total root proteins we detected were also present there. This intersection of 7577 proteins between our two respective studies allows us to define the core set of proteins most abundant in the root, the root core proteome.

### Seed Proteome

Cluster 7 (Figure 2 D) contains proteins whose abundance increased specifically in seed. As expected, many of these were the more abundant seed storage proteins such as oleosins, albumins, cruciferins and enzymes of sugar and fatty acid metabolism. However, many of the core transcription factor, phosphatase and chromatin modifier modules that regulate seed development were also detected and quantified (Supplementary table 8 and Figure 2 D). The PP2C POL directs *WUS/WOX5* gene expression and is essential for meristem establishment and stem cell maintenance in the early embryo together with PLL1 (Song et al., 2008). Similarly, the AIL transcription factor BBM and HDG1 act antagonistically to balance stem cell proliferation and differentiation (Horstman et al., 2015) and AN3 establishes cotyledon identity upstream of PLETHORA1 (Kanei et al., 2012). The ABA signaling proteins which are known to play a major role in seed development were underrepresented in cluster 7 mainly because they either did not accumulate exclusively in seed or not in both of the measured stages (developing green and ripe brown siliques) but there abundance could be easily reconstituted from the data. LEC1, the master inducer of seed development was highly abundant in the earlier stage together with AREB3 and EEL, two ABA responsive bZIP transcription factors that govern early seed maturation (Agarwal et al., 2011). The ratio of the bZIP transcription factor ABI5 to AFP1, a repressor of ABA signaling, both detected exclusively in mature to post mature seeds was greater than 1 indicative of high ABA levels and induction and possible maintenance of seed dormancy in the brown siliques sample. Concomitantly the DOG1 protein and the RDO5 PP2C that are essential for seed dormancy (dormancy is completely abolished in the *dog1* knockout mutant also in the presence of ABA (Nee et al., 2017)) were abundant in the post mature seed proteome. A number of other PP2C proteins also accumulated to high levels specifically in siliques, particularly in brown siliques. The same was true for several members of a clade of NAC transcription factors also hitherto not known to play a role in seed development indicating potential functions for them in seed development and establishment of dormancy.

### Ribosomal Proteins in Development

The abundance of the 315 proteins in cluster 4 strongly increased or was exclusively measured in young roots, young stem and early flowers/floral buds. There abundance decreased in the latter two tissues as development progressed (Figure 2 E). More than half of these proteins were localized in the nucleus and a substantial number of them pertained to nucleolar processes and ribosome biogenesis with WD40 proteins, containing the WD40 repeat molecular interaction domain, being highly significantly enriched (Supplemental table 9).

One of these was LEUNIG (LUG), a transcriptional co-repressor and master regulator of flower development that directly modulates antagonistic A and C class gene expression in the inner and outer whorls (Grigorova et al., 2011). TAIR10 annotated six WD40 proteins as DWD components of CUL4 RING E3 ubiquitin ligases (Supplemental table 9) but closer inspection revealed that AT3G56990 (EDA7) and AT4G04940 actually lacked the canonical DWD motif (Supplemental material and data), while the others belonged to the A clade of Arabidopsis DWD proteins (Lee et al., 2008). Among them was DWA2, a negative regulator of ABA signaling that targets ABI5 to degradation (Lee et al., 2010).

We investigated potential interactions among the 18 WD40 proteins in cluster 4 using the STRING database set to produce essentially only true positive binary physical interactions. Five of the six components of the UTP-B complex, a sub-complex of the SSU-processome/90S pre-ribososme, were found to interact based on experimental evidence from yeast and human (Gavin et al., 2002; Gavin et al., 2006; Krogan et al., 2006; Wan et al., 2015), showing tissue and development specific expression of this entire protein complex (Figure 2 E, Supplemental figure 6 and Supplemental table 9 for raw abundance values). The complex interacted with EDA7, the homolog of enp2 (Soltanieh et al., 2014), a putative ribosome biogenesis factor (RBF) which is not known in *Arabidopsis*. NuGWD1 a sugar inducible WD40 protein was reported to interact with SLOW WALKER 1 (SWA1) which is a component of the t-UTP sub-complex of the SSU processome and which was also shown to interact with several UTP-B members by co-IP MS in *Arabidopsis* (Ishida et al., 2016). SWA1 is not a WD40 protein but the pattern of its abundance was very similar to the protein expression pattern of the cluster 4 proteins (Supplemental Table 9 for raw abundance values). Another WD40 protein containing complex, all of whose components were measured and showed the same expression pattern, elevated in roots, young shoots and early flowers/floral buds was the PeBoW complex which is essential for pre-rRNA processing (Cho et al., 2013; Ahn et al., 2016). Additionally, extensive physical interaction between a large number of ribosomal (RPs) and other ribosome associated proteins with this expression pattern became evident when interactions between all 315 cluster 4 proteins was assayed (Supplemental figure 6). However, the cluster 4 expression pattern of the RBF protein complexes was significantly more conserved than expression patterns of ribosomal proteins (Supplemental Figure 7).

Molecular chaperones were also highly significantly enriched among the cluster 4 proteins. Nine of these 21 proteins putatively interact physically based on x-ray crystallography and tandem affinity purification studies in yeast and human (Dekker et al., 2011; Hauri et al., 2016). All of these TCP-1/cpn60 proteins are homologs of the yeast and human cytosolic chaperonin CCT proteins suggesting they constitute the hardly described CCT complex in *Arabidopsis* and that it’s abundance is tissue and developmentally specific (Figure 2 E and Supplemental table 9 for raw abundance values).

### Vesicle Trafficking and Transport

A set of 153 proteins were abundant in roots, increased during ageing and peaking in senescence in leaves and increased in the young inflorescence stem and flowers (Cluster 13, Figure 2 F Supplemental table 10). These proteins were largely related to vesicle trafficking, Golgi apparatus and membrane transport. They contained numerous exocyst complex members, SNAREs and cytoskeletal proteins such as actin. Vesicle trafficking and cytoskeletal remodeling and organization are central processes in tip growing cells which are well studied in root hairs (Rounds and Bezanilla, 2013), but are also prevalent in fast growing tissues such as the emerging inflorescence stem and flowers (Chen et al., 2009). On the other hand, autophagy, a process which also involves the formation of lytic compartments and vesicle trafficking for the degradation of cytoplasmic material is known to play an important role in senescence in leaves and other tissues (Wojciechowska et al., 2018).

### The Proteome in Establishment and Maintenance of Photosynthesis

Clusters 11 and 8 (Supplemental tables 11 and 12 respectively) were almost exclusively comprised of proteins annotated as chloroplast localized (297 of 326 nuclear plus 18 plastid encoded and 223 of 264 nuclear plus 20 plastid encoded, respectively) and essential to photosynthesis by DAVID GO. Proteins were absent in roots and mature brown siliques and predominant in green tissues, primarily leaves and cauline leaves. The abundance of cluster 11 proteins peaked in cotyledons of 7 and 10 day old seedlings and young cauline leaves and declined in these tissues in the course of ageing, whereas cluster 8 proteins were consistently abundant throughout the plant lifecycle, in rosette leaves with a slight maximum at 40 days and declining in senescence (Figure 3 A and B).

**Figure 3.**
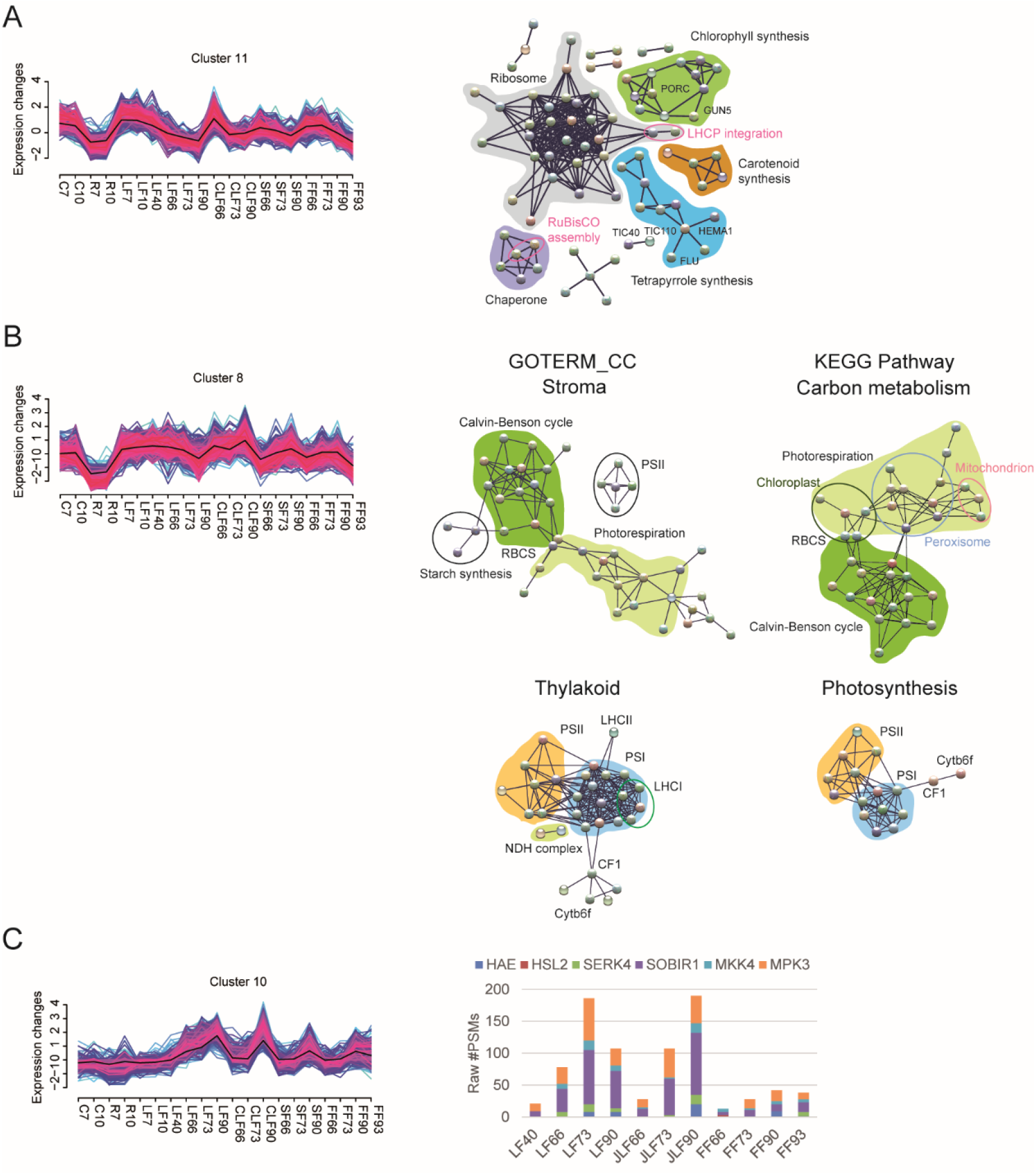
Proteome remodeling in establishment and maintenance of photosynthesis as well as in degradation of photosynthesis apparatus and senescence in leaves. A. FCM cluster 11. Right panel: Physical / functional protein interaction networks generated with the STRING database using all proteins assigned to the GOTERM category Chloroplast (Supplemental Table 11) as input set. Various biochemical pathways in pigment and photosynthesis related protein synthesis and chloroplast biogenesis are color coded and indicated. B. FCM cluster 8. Right panel: Physical / functional protein interaction networks generated with the STRING database using all proteins assigned to the GOTERM categories Stroma, Thylakoid, Carbon metabolism (C Metabolism) and Photosynthesis as input sets (Supplemental Table 12). Light independent reactions are highlighted in green (Calvin-Benson cycle dark, photorespiration light), light dependent reactions in orange (photosystem II PSII) and blue (photosystem I PSI). C. FCM Cluster 10. Right panel Cumulative abundance (PQI given is raw #PSMs) of core abscission signaling module proteins in leaves, cauline leaves, flowers and siliques.

The proteins in cluster 11 primarily function in chloroplast biogenesis and the establishment of the functional photosynthetic machinery. A number of central players in these molecular events were identified including HEMA1 and GUN5 (as well as GUN4), (Liu et al., 2017) that also represent hubs in STRING generated networks of cluster member proteins interacting in tetrapyrrole and chlorophyll synthesis, respectively (Figure 3 A, complete annotated network Supplemental figure 8). Other cluster proteins were interaction partners in carotenoid synthesis. Around 30 cluster proteins were classified as ribosomal proteins or related to translation by DAVID GO which was reflected by a highly connected interaction network composed of chloroplast specific ribosomal proteins, ribosome associated proteins or RBFs, the gene knockouts of many of which have embryo lethal phenotypes. CPN60 and CPN10 class molecular chaperones formed another cluster of interactors. Protein import from the cytosol is essential to chloroplast biogenesis and constituents of the Toc/Tic complex including TOC33, which is known to be most strongly expressed in young seedlings were cluster members (Waters and Langdale, 2009). The two proteins forming the inner division ring mediating chloroplast division, FtsZ1 and FtsZ2, were also cluster members (Waters and Langdale, 2009). DAVID GO mapped the cluster proteins to several plastid structures thereby giving a possible inkling of their localization within the chloroplast.

Cluster 8 mostly contained proteins prevalent in mature chloroplast directly related to photosynthesis. Cluster members constituted extensive parts of both the light dependent and independent reactions. Regarding the former, most components of the oxygen evolving photosystem II (PSII) and photosystem I (PSI), providing the final strong reducing potential, were identified, many of them being close interactors (Figure 3 B). Components of the electron transferring Cytb_6_f complex and the thylakoid ATP synthase CF_1_ were also present. Regarding carbon fixation, nearly all components of the Calvin-Benson cycle and many of the proteins involved in photorespiration were mapped as functional interaction partners (Figure 3 B). Furthermore, functional interaction mapping could resolve organelle specificity of the individual reactions in photorespiration. Interaction networks produced from GO category stroma and carbon metabolism and from GO category thylakoid and photosynthesis protein sets were specific and showed extensive overlap attesting to the quality of protein extraction from the chloroplast compartments (full annotated networks Supplemental figure 9 to 12).

### The Proteome in Senescence

Cluster 10 contained 241 proteins whose abundance increased substantially during the course of plant life and peaked in the latest developmental stage, i.e. during leaf and flower senescence and fruit ripening, predominantly in rosette leaves but also in cauline leaves, stem and flowers (Supplemental table 13). They were not abundant in young tissues (Figure 3 C).

Senescence is a controlled developmental process that entails disassembly of the photosynthetic apparatus in leaves for the purpose of nutrient remobilization and resource allocation to fruit ripening in flower petals (Bleecker and Patterson, 1997). Conversely, numerous proteins involved in chlorophyll and carotenoid degradation were found including Pheophorbide a oxidase that is the key enzyme in the formation of primary fluorescent chlorophyll catabolites (pFCCs). It has been shown that nonfluorescent dioxobilin-type chlorophyll catabolites (NDCCs) represent the major end- products of chlorophyll catabolism as opposed to NCCs and that the Cytochrome P450 monooxygenase CYP89A9 is responsible for their accumulation (Christ et al., 2013). This protein was also a cluster member as well as 13 other CYPs. Twelve of the 14 total CYPs belonged to the CYP71 clan suggesting a broader role for these proteins going beyond CYP89A9.

During senescence large amounts of ROS are produced which must be controlled so as not to lead to tissue damage and premature cell death (Rogers and Munne-Bosch, 2016). Many cluster proteins were involved in oxidative-reductive processes and ROS scavenging including the centrally important cytosolic ascorbate peroxidase APX6. More interestingly, a significant group of cluster proteins were kinases. Six of these (CRK7, CRK8, CRK10; CRK14, CRK21, CRK41) were Cysteine-rich Receptor-like Kinases (CRKs) that have two extracellular DUF26 domains that each contain 4 conserved cysteine residues. Next to the programmed loss of redox control leading to ROS accumulation and ultimately cell death at the later stages of senescence, ROS may also play an important role in signaling, mediating genetic reprogramming during senescence (Breeze et al., 2011), the mechanisms of which however are largely unexplored. The CRK cysteine thiol groups will likely be sensitive to the redox state, potentially implicating these proteins as ROS sensors and ROS signaling initiators in senescence.

Abscission of floral organs after fertilization is another developmental process that occurs late in the *Arabidopsis* life cycle. Most of the proteins of the canonical abscission signaling module (Meng et al., 2016) including the receptor like protein kinases HAESA (HAE), HAESA-like 2 (HSL2), and EVERSHED (EVR/SOBIR1), the co-receptor SOMATIC EMBRYOGENESIS RECEPTOR-LIKE KINASE 4 (SERK4) and MKK4 and MPK3, the mitogen associated protein kinase cascade downstream of the HAE receptor complex were cluster members. The proteins accumulated in flowers and later siliques as development progressed (Figure 3 C right panel). Interestingly, their abundance also increased in leaves and cauline leaves although leaf abscission is not developmentally timed suggesting an unknown function of this abscission signaling module.

### Proteome Remodeling in Steady State PTI

In addition to tissue proteomes and their altercation in the course of development, we were interested in assaying more rapid changes in proteome architecture such as determined by steady state shifts from ordinary growth to immunity. Seven and ten day old seedlings grown in liquid culture were treated with a final concentration of 1 µM flg22 in the medium for 16 hours. Flg22 is the 22 amino acid N-terminal epitope of flagellin and elicits pattern triggered immunity (PTI) downstream of the receptor like kinase (LRR-RLK) FLAGELLIN-SENSITIVE 2 (FLS2). Deep proteomics measurement of the untreated and treated samples identified 8344 proteins in all (Supplemental table 14). HCL showed that the abundance of 1774 proteins increased whereas the abundance of 915 decreased in both samples as a result of flg22 exposure (Supplemental table 15).

These proteins were categorized by mapping them to a self-constructed model of PTI using the MapMan software (Supplemental data for details, Supplemental table 16, Supplemental figure 13). The result was a comprehensive picture of proteome remodeling in PTI showing extensive changes in protein abundance in most of the major perception, signaling and response pathways. Dynamics of the flg22 receptor complex, including decreased abundance of FLS2 as a result of internalization and degradation (Robatzek et al., 2006) and a host of other RLKs were quantified. Vesicle trafficking and transport proteins including exocyst and SNARE complex members increased in abundance along with proteins involved in the early respiratory burst, primarily RESPIRATORY BURST OXIDASE HOMOLOGUE D (RBOHD). All of the components of both MAPK signaling cascades central to plant immunity, MKK4/5-MPK3/6/11 and MKK2-MPK4 (Bigeard et al., 2015) were measured and also showed a slight increase in their abundance. The same goes for the calcium-dependent protein kinases (CDPKs) integral to PTI signaling, CPKs 4, 5 and 6 (Boudsocq et al., 2010), as well as a host of other CDPKs, calmodulin (CAM) and CAM binding proteins and calcineurin (Supplemental table 17). Interestingly several proteins showing changes in their abundance upon PAMP stimulus were mapped to other avenues of plant immunity such as effector triggered immunity (ETI) and programmed cell death (PCD) and systemic acquired resistance (SAR).

We investigated proteome plasticity in more detail beginning with the category hormone signaling and branching out from it to produce a proteomics model of phytohormone activity in PTI (Figure 4 Supplemental table 18). The deep proteomics results were complemented by measurements of the plant hormones themselves, amino acids and secondary metabolites after 16 hours of flg22 treatment. Furthermore, we undertook a retention time scheduled, parallel reaction monitoring (PRM) based targeted proteomics study that allowed accurate quantification and inference of statistical significance of fold changes of 52 model proteins in three sample pools of 10 day old *Arabidopsis* seedlings again following 16 hours of flg22 treatment (Figure 5, Supplemental Table 19). The PRM based estimates of protein abundance fold changes were in full agreement with the deep proteomics quantification highlighting the latter’s accuracy and power of the deep proteomics strategy in general.

**Figure 4.**
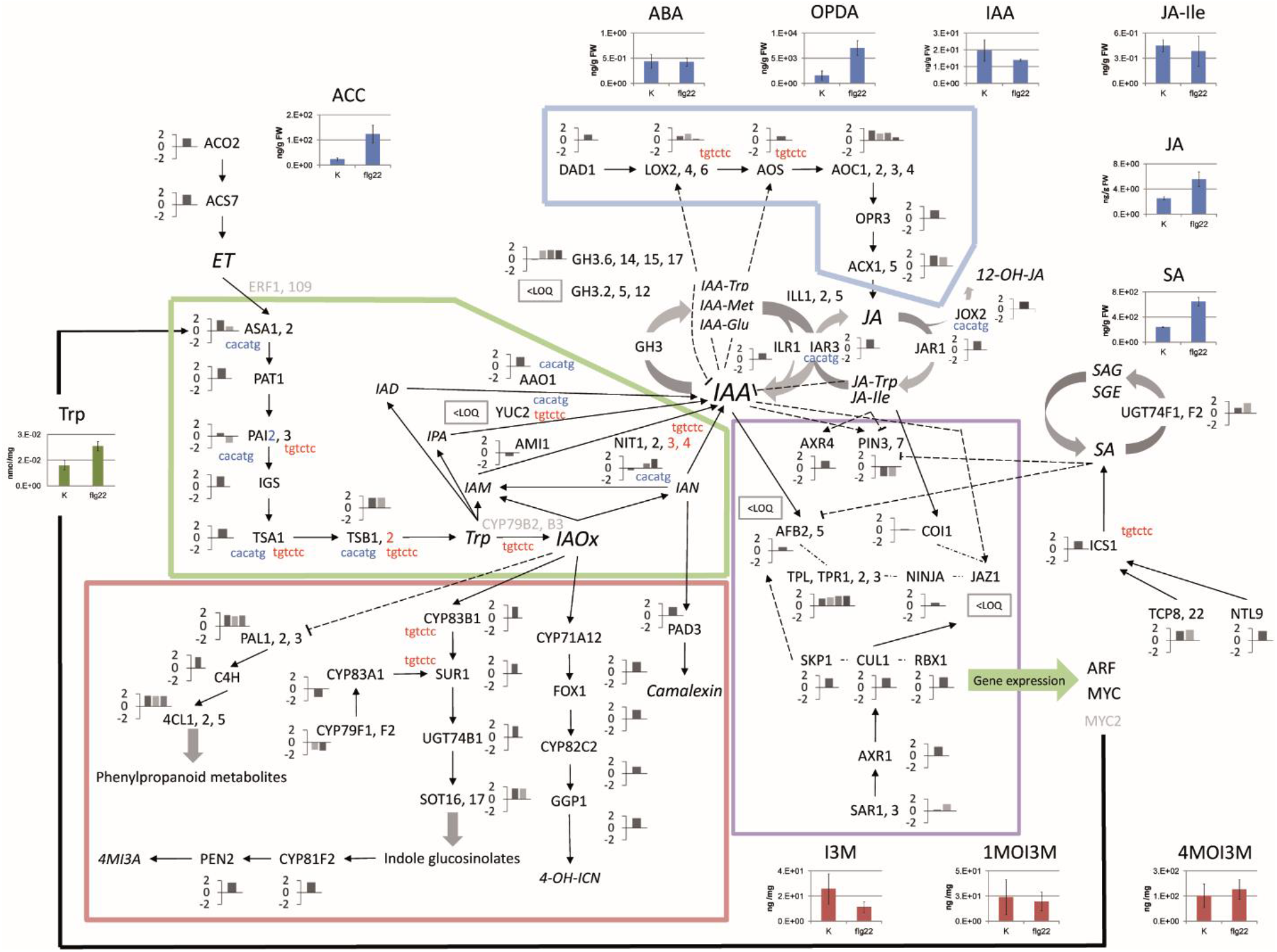
Proteomics model of phytohormone activity in PTI. The figure shows the proteins quantified in the jasmonate (blue), tryptophan and IAA (green), secondary defense compounds/indolic glucosinolate (red) biosynthesis pathways as well as IAA and JA signaling pathways (purple). Note all or nearly all components of these biochemical pathways were measured and quantified in the deep proteomics study. Proteins pertaining to ET and SA synthesis as well as reversible hormone conjugation (GH3s) and modification are also shown. Protein with names in black were measured; bar charts next to protein names indicate relative changes in abundance after flg22 treatment; values are the sum of z-score transformed raw #PSMs in flg22 treated samples. Proteins in grey were not detected. Red nucleotide sequence indicates an ARF1 binding site in the cognate genes promotor region, blue a MYC2 binding site. Solid arrows indicate direct functional interaction or immediately neighboring steps in biochemical pathways. Dashed arrows indicate more distal interactions. Phytohormone, tryptophan and indole glucosinolate abundance 16 hours after flg22 treatment are also shown.

**Figure 5.**
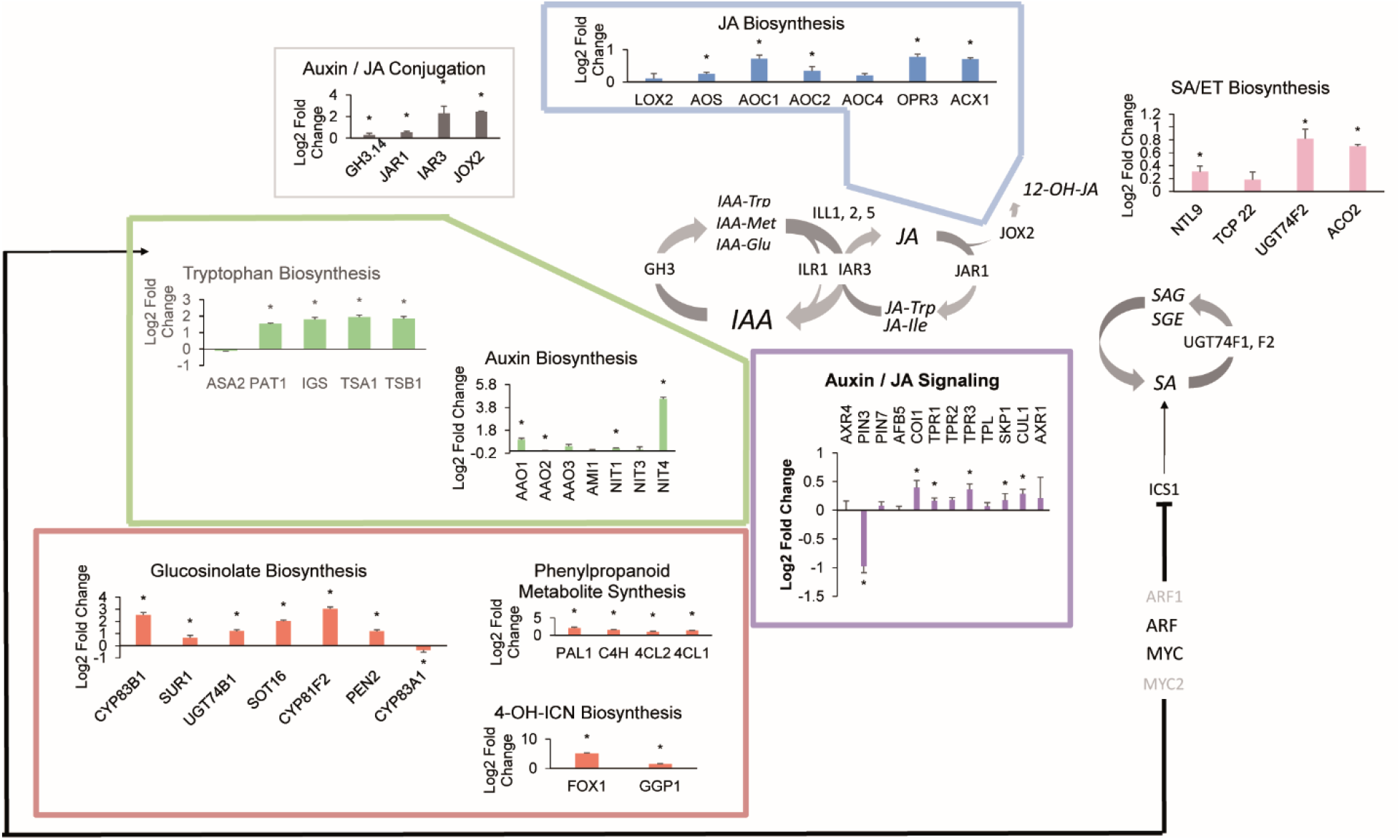
Targeted PRM based quantification of model proteins. Framework is in analogy to figure 4. Bars represent log2 fold changes of protein abundance after 16 hours of flg22 exposure (1µM concentration in medium) estimated by area under the curve label free protein quantification index (PQI) of the 6 most intense product ions from MS2 spectra of targeted proteotypic peptides. Bars represent median PQI of all quantified proteotypic peptides for a given protein in 9 measurements (3 biological replicates each measured 3 times). Standard error is indicated. Star indicates significance α=0.05 if fold change of at least one of the quantified peptides was significant.

The abundance of 43 proteins playing roles in photosynthesis decreased slightly after 16 hours of exposure to flg22 (Supplemental table 20). Fifteen of these were assigned to the photosynthesis light reaction and 6 of these in turn to photosystem II by the MapMan software (MapMan bins PS.lightreaction and PS.lightreaction.photosystem II. LHC-II respectively, Supplemental table 20). Both bin assignments were statistically significant (Benjamini Hochberg corrected p-values <0.01) indicating these categories were enriched among the 43 proteins. Other proteins were assigned to the carbon reactions, more specifically the Calvin-Benson cycle (Supplemental table 20). The relatively small decrease in abundance of a set of these proteins was corroborated by small yet significant fold changes (maximum decrease −0.6 log_2_ fold change) in the PRM study (Supplemental figure 14 and Supplemental table 19). These results suggest that photosynthesis is inhibited upon PAMP perception and conversely in PTI.

The abundance of 93 proteins playing roles in plant primary metabolism, especially carbohydrate metabolism, increased upon exposure to flg22. 22 were categorized as pertaining to glycolysis (MapMan bin glycolysis), 19 to the TCA cycle (MapMan bins TCA / org. transformation.TCA and TCA / org. transformation.other organic acid transformations) and 6 to major CHO metabolism (mapMan bin major CHO metabolism.degradation.starch), among others by the MapMan software. Increases in abundance were more pronounced than the downregulation of photosynthesis related proteins and log_2_ fold changes up to 1.6 were quantified and significant in the PRM study (Supplemental figure 14 and Supplemental table 19).

### Jasmonate and Salicylic Acid Cross Talk

Sixteen hours of flg22 exposure led to an increase in the abundance of both salicylic acid (SA) synthesis pathway proteins, primarily ISOCHORISMATE SYNTHESIS 1 (ICS1) but also upstream transcription factors TEOSINTE BRANCHED1/CYCLOIDEA/PCF 8 and 22 (TCP8, TCP22) (Wang et al., 2015) and NTM1-Like 9 (NTL9) (Zheng et al., 2015) and free SA levels (nearly 3-fold).

Jasmonate (JA) and jasmonate-isoleucine (JA-Ile), its bioactive conjugate, levels were both low, although the former increased around 2-fold, the latter showed no significant change and absolute levels of both were in the range of a few ng/g FW. Concurrently, (+)-12-oxo-phytodienoic acid (OPDA) levels increased from 1.6 to 7 µg/g FW so were high and elevated more than 4-fold following flg22 treatment. OPDA is a primary JA precursor synthesized from allene oxides by ALLENE OXIDE CYCLASE 1 to 4 (AOC1 to 4). ACC, the ethylene precursor and a proxy for the phytohormone’s abundance increased more than 5-fold.

Controversially, the deep proteomics measurements showed elevated protein amounts of all components of the JA biosynthesis pathway, corroborated as significant in the PRM study. Moreover the core JA receptor complex / signaling proteins CORONATINE INSENSITIVE 1 (COI1), TOPLESS RELATED PROTEINS 1 to 3 (TPR1 to 3) and S PHASE KINASE-ASSOCIATED PROTEIN 1 (SKP1) and CULLIN 1 (CUL1) proteins, members of the E3 ubiquitin ligase SCF complex, were all significantly increased in abundance, albeit slightly. The abundance of a large number of GRETCHEN HAGEN (GH) enzymes, the amidohydrolase IAA-ALA RESISTANT3 (IAR3) and JASMONATE-INDUCED OXYGENASE 2 (JOX2) also increased following exposure to the PAMP, the latter two significantly around 5-fold. These enzymes are all involved in conjugating or deconjugating phytohormones, specifically JA and auxin (IAA) to amino acids or small molecules or hydroxylating them (JOX2) thereby modulating their sub-cellular location and/or rendering them active or inactive.

The proteomics results imply, that flg22 induced PTI prioritizes SA over JA signaling. On the other hand, the induction of both JA synthesis and signaling pathways on the protein level and the highly elevated abundance of both IAR3 and JOX2, which deconjugate JA-Ile to JA and hydroxylate the latter to 12-OH-JA, together with the low JA and JA-Ile levels themselves, suggests that phytohormone conjugation may play a pivotal role in this context and a role for JA in PTI.

### Auxin/IAA homeostasis

The detection of numerous GH proteins known to conjugate auxin/IAA (GH3.2, GH3.5/WES1 and GH3.17/VAS2) and others prompted us to investigate the role of this phytohormone in PTI. The levels of GH3.14, GH3.15 and GH3.17 increased upon flg22 exposure, the first significantly 1.2 fold. GH3.15 (AT5G13370) function in conjugating the IAA precursor IBA has just recently been elucidated (Sherp et al., 2018). GH3.14 (AT5G13360), the neighboring gene, does not shown any significant sequence homology indicating a potentially different uncharacterized function in PTI.

Auxin/IAA is synthesized by tryptophan dependent and independent pathways. Both deep and targeted proteomics results showed significant substantial increase in the abundance of all proteins in the tryptophan biosynthesis pathway and tryptophan levels were also elevated almost 2-fold upon 16 hours of PAMP treatment. Tryptophan channels into a host of defense related secondary metabolite synthesis pathways particularly indole glucosinolates (IGs), the protein abundances of which all increased highly (2-fold or more) and significantly. IG levels themselves, indol-3-ylmethylglucosinolate (I3M), 1-methoxy-indol-3-ylmethylglucosinolate (1MOI3M) and 4-methoxy-indol-3-ylmethylglucosinolate (4MOI3M), did not change significantly. Presumably they were hydrolyzed by the mirosinase PENETRATION 2 (PEN2), whose abundance, along with CYP81F2, which showed an 8.2 fold change in abundance, increased 2.3 fold in response to flg22, to play a role in callose deposition (Clay et al., 2009).

Several proteins potentially involved in auxin/IAA synthesis pathways were identified. The abundance of ALDEHYDE OXIDASE 1 (AAO1) increased nearly 2-fold and that of the NITRILASES 1 and 4, 1.2 and 23 fold, respectively. Auxin/IAA levels however decreased slightly (0.75 fold) but significantly (p-value 0.051, two sample t-test equal variance, n=5, α=0.1). AAO1 has recently been shown to play a role in converting the I3M downstream hydrolysis product indole-3-carbaldehyde (ICHO) to indole-3-carboxylic acid (ICOOH) in the abiotic stress response (Muller et al., 2019). *Pseudomonas syringae pv tomato DC3000* (*Pst* DC3000) infection induced a strong transcriptional response of the nitrilases NIT2, NIT3 and NIT4 which was corroborated by increased protein abundance in the case of NIT2. It was postulated that NIT2 is involved in IAA signaling in R gene mediated resistance and defense against biotrophic pathogens (Choi du et al., 2016). The more than 20-fold increase in the abundance of NIT4 that we measured suggests an unknown function of this protein in basal resistance to biotrophic pathogens.

While auxin/IAA levels did not change substantially even 16 hours after PTI induction, the abundance of polar auxin transporters was markedly affected. Particularly, the abundance of PIN-FORMED 3 (PIN3), a polar efflux carrier with well documented functions in lateral root growth and tropism decreased significantly 2-fold. A similar decrease was apparent for PIN-FORMED 7 (PIN7) in the deep proteomics results which is also known to affect lateral root growth. Both PIN 3 and 7 have important roles in establishing local auxin concentration maxima (Jang et al., 2018). The abundance of AUXIN RESISTANT 4 (AXR4) an ER protein that is responsible specifically for the distribution and localization of the influx carrier AUXIN RESISTANT 1 (AUX1) was also elevated slightly. PIN3, PIN7 and AUX1 are the primary components of the polar auxin transport system so our results indicate active, cell-to-cell auxin transport is perturbed in steady sate PTI.

### Chronology and Model of Phytohormones in PTI

It has been documented that JA, IAA and SA act in chronological order in the establishment of SAR and that JA plays an early role (Truman et al., 2010a). Therefore, we measured hormone levels 1, 3 and 16 hours and OPR3 (as a marker for JA synthesis), JOX2 and COI1 and TPR1 (as markers for JA signaling) transcript levels 3 and 16 hours after flg22 exposure (Supplemental tables 21 and 22 and Figure 6). SA accumulated after 1h and levels remained elevated until 16h after exposure. JA and JA-Ile levels remained basal throughout the entire time course. OPDA levels however increased substantially already 1 hour after PAMP treatment and remained elevated over time and OPR3 transcript abundance increased significantly 5.7 fold 3 hours after PAMP exposure and decreased to basal levels at the 16 hour time point. Together this suggests that the JA biosynthesis pathway is induced early in PTI. JOX2 transcripts were elevated substantially and significantly 3 hours and remained high up to 16 hours after exposure to flg22 again reconciling initiation of JA synthesis with low JA levels by way of hydroxylation already at the early stages of biotrophic pathogen defense. The abundance of COI1 and TPR1 transcripts did not change markedly over time suggesting little or no JA signaling in the absence of JA-Ile itself.

**Figure 6.**
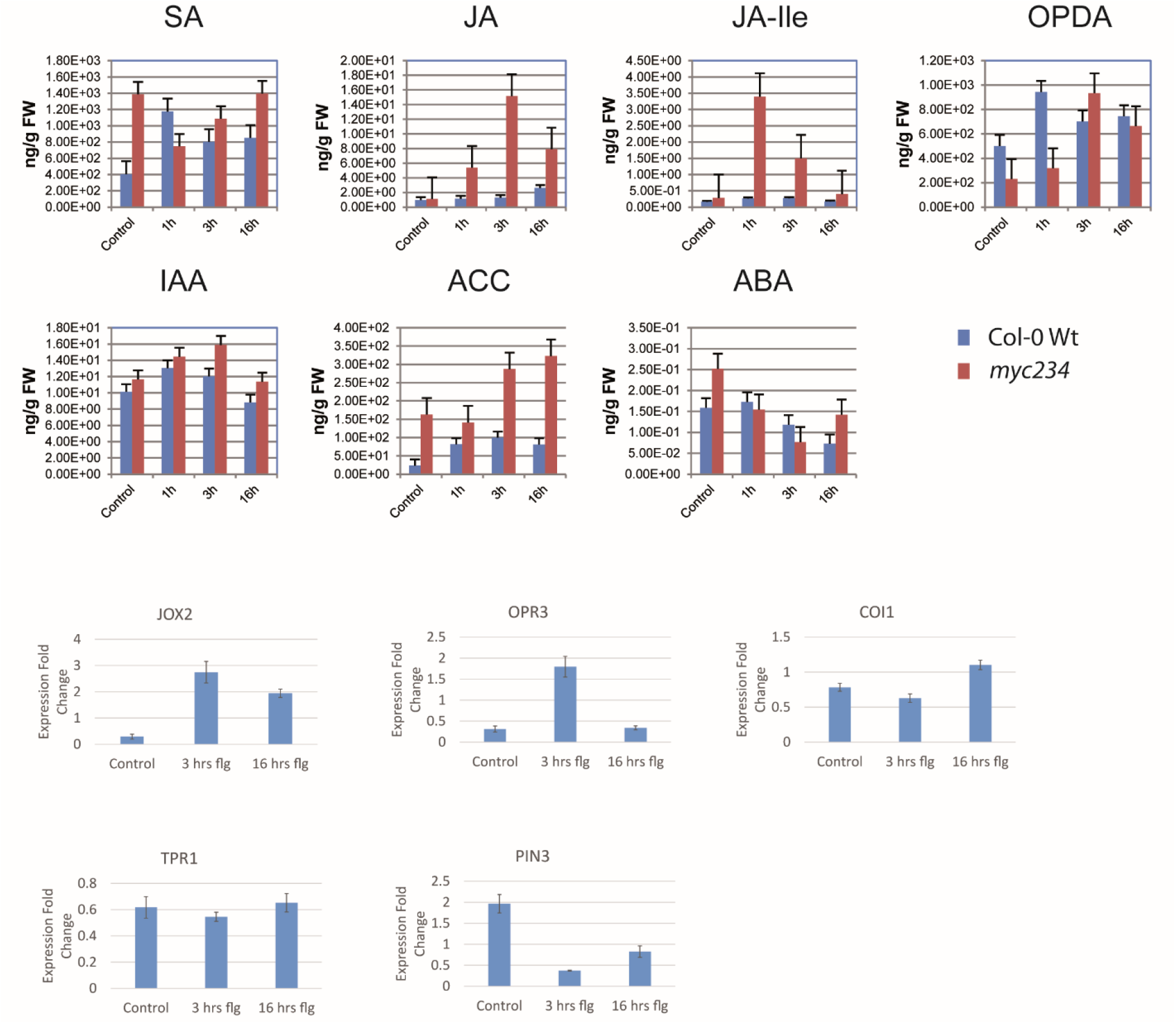
Absolute quantification of phytohormones without and 1, 3 and 16 hours after flg22 exposure (1µM concentration in medium) in Col-0 Wt and myc234 mutant backgrounds using GC-MS. Bars represent mean of three biological replicates; standard error is given. qPCR based transcript abundance profiling of selected genes without and 3 and 6 hours after flg22 exposure (1µM concentration in medium). Bars represent mean of 6 measurements (3 biological replicates each measured twice), standard error is given.

Finally, as we uncovered several potentially new aspects of JA synthesis and homeostasis in PTI, we measured the abundance of the phytohormones at the same time points following flg22 treatment in the *myc234* background. JASMONATE INSENSITIVE 1 (MYC2) is one of the most important transcription factors in JA signaling downstream of COI1 with a host of diverse regulatory functions and the triple knockout exhibits essentially no functional redundancy. SA hyperaccumulated in the triple KO under standard growth conditions as described previously (Nickstadt et al., 2004). Its abundance decreased markedly 0.54- and 0.78-fold 1 and 3 hours, but increased 16 hours after flg22 perception, suggesting that JA signaling plays a role in SA accumulation in PTI and that it is an early one.

Strikingly, JA and JA-Ile levels both greatly increased (maximum JA-Ile 12.1-fold 1 hour, JA 13.5-fold 3 hours after flg22 exposure) throughout the time course. OPDA also increased in the mutant, its profile mirroring that of JA in the wild type, reaching a maximum increase of 4-fold three hours after flg22 perception. This shows induction of JA biosynthesis pathway and synthesis of JA intermediates independently of MYC2. The highly elevated JA levels and JOX2 transcript abundance 3 hours post PAMP treatment prompted us to investigate the IAR3 and JOX2 promotor regions for MYC2 binding sites and indeed two and three were identified as the top ranking hits repsectively. This supports a hypothesis wherein JOX2 expression is under the control of MYC2 and wherein JA is synthesized but continuously depleted by JOX2 by way of hydroxylation in biotrophic pathogen defense. Our combined measurements represent strong evidence of a MYC2 dependent negative feedback loop controlling JA-Ile and JA levels in PTI via deconjugation and hydroxylation by IAR3 and JOX2 (Figure 7).

**Figure 7.**
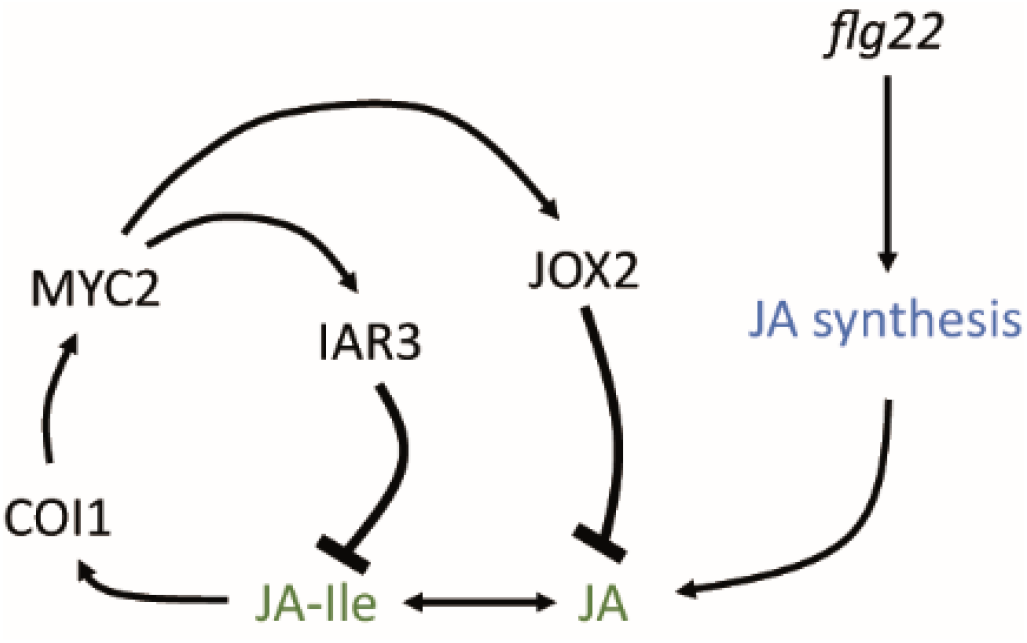
A MYC2 dependent negative feed-back loop controls JA-Ile and JA levels in equilibrium with JA synthesis by way of deconjugation and hydroxylation via IAR3 and JOX2 in immunity to biotrophic pathogens.

The other phytohormones showed similar levels over time in both the wild type and the mutant. The amount of free IAA did not change dramatically, increasing slightly at earlier time points and decreasing as stated above at the 16 hour time point presumably because steady state PTI is achieved. ACC/ET abundance increased already 1 hour after exposure of the seedlings to the PAMP and remained elevated. ABA decreased 3 hours after flg22 treatment and remained low throughout the time course, presumably because of its known antagonism with SA (Cao et al., 2011). Additionally we measured changes in the levels of the 20 proteinogenic amino acids and some others and found alanine, glycine, tryptophane and taurine to increase during the course of flg22 exposure (Supplemental figure 15). The abundance of leucine, isoleucine, lysine, proline and ornithine decreased over the time of exposure.

## Discussion

### Deep Proteomics Study of the *Arabidopsis thaliana* Proteome

In this report we describe the proteome biology of the model plant *Arabidopsis thaliana* comprehensively. We elucidated the proteome landscapes of the plant and tissues throughout its lifecycle as well as in the immune response to PAMP in unprecedented resolution and detail. Importantly we placed an emphasis on remodeling of proteome architecture between these diverse biological scenarios to extrapolate genome wide protein co-regulation and function. This approach has just recently been documented in humans (Kustatscher et al., 2019) and draws its power from deep coverage of the proteome as we have done here, quantifying close to 16,000 *Arabidopsis* proteins. In-depth sampling allowed quantitative measurement of all of the components of entire biochemical pathways such as JA synthesis in PTI, protein complex components such as RBF complexes and protein co-expression of genes in local neighborhoods such as cysteine rich kinases (CRKs), facilitating confident inference of protein abundance co-regulation and functional and temporal connections for instance in seed development and dormancy, the establishment, maintenance and deconstruction of the photosynthetic machinery and in senescence and abscission. With this we alleviated an inherent weakness of most previous plant proteomics studies that were underpowered, quantifying only a few thousand proteins if at all. To do this we developed and optimized a strategy that in our hands allowed quantification of more than 9,000 proteins from a sample in 3.5 days, a duration that is acceptable for medium scale proteomics studies.

Despite our unprecedented sampling depth, we did not delve much deeper into the entire *Arabidopsis* proteome than Baerenfaller and co-workers (or as a matter of fact the other deep proteomics studies in plants). In their classic study published more than ten years ago individual sampling depth was modest however they utilized more than 1,300 low resolution LC-MS measurements to quantify nearly 15,000 proteins. Both studies show a similar impact of transcript abundance on detection of cognate proteins (Figure 2C Baerenfaller et al, Figure 1C here) however we show protein size has a more pronounced effect presumably because small proteins will generate less tryptic peptides amenable to MS detection. Also transcription factors and nuclear proteins whose expression may be highly transient and biological context specific were conspicuously overrepresented in the missing proteome (Supplemental figure 4). Capturing these proteins could require exhaustive sampling of different organs at all developmental stages (imagine par example the diversity of the 12 stages of organ development in the floral bud (Bowman et al., 1989; Smyth et al., 1990)) and environmental interactions and even then may escape detection. Indeed, the more than 8,000 proteins identified in the context of flg22 treatment did not substantially increase the amount of total proteins previously identified in the plant tissues (Figure 1F) Also proteomics ruler method allowed us to quantify proteins with less than 100 copies per cell, which is in general agreement with LOQs of modern large scale proteomics studies in humans (Bekker-Jensen et al., 2017). This suggests that even more comprehensive sampling of the *Arabidopsis thaliana* proteome may be confounding with current technologies.

Next to the prevalence of known artificial and biologically occurring modifications, amino acid substitutions were found to be common. Such exchanges are generally caused by ribosome infidelity, have a number of biological implications and have been reported previously in human cell lines (Chick et al., 2015), human tissue (Bagwan et al., 2018) and in *E.coli* and yeast in detail (Mordret et al., 2019). The authors of the latter study define exchanges as near cognate errors (NeCE) when two of the three bases between error bearing origin and destination codon match and as non-cognate error (NoCE) when there is no such match between possible error bearing origin and destination codons. Interestingly, 5 of the 8 substitutions we found to be abundant in *Arabidopsis* (Asn->Met, Ala->Asp, Pro->Glu, Asp->Asn, Glu->Gln) were also abundant in yeast but not *E.coli*, and three (Ala->Asp, Asp->Asn, Glu->Gln) were NeCEs suggesting conserved patterns of translational error in eukaryotes, a possible method of generating random phenotype variants in genetically identical organisms (Mordret et al., 2019). This phenomenon may also highlight the limitations of searching protein databases derived from reference genomes with proteomics data.

### Root and Seed Proteomes

The root is a highly specialized, plastic tissue that confers structural stability to the above ground portion of the plant and is responsible for the uptake of nutrients and water from the soil. Thus it exudes a number of metabolites, peptides and proteins that allow it to interact with the rhizobiome. The function of the root specific proteins in cluster 3 are well documented. Interestingly the transcription factors and proteins that pertain to hormone signaling in the developing root (Paez Valencia et al., 2016) were not overly prominent among the set of cluster proteins. Comparative evaluation of different deep proteomics studies as we have done here can be useful in defining the core set of proteins ubiquitously expressed in a cell type, tissue or organism.

The proteins in cluster 7 are very specific to the seed proteome. They were exclusively or significantly highly abundant in siliques only and not ubiquitously abundant in other tissues. These include many of the proteins involved in ABA dependent regulation of seed development. The amount of ABI5, a positive regulator of ABA signaling is post translationally controlled by AFP1, presumably promoting its degradation by the proteasome (Lynch et al., 2017). The high ABI5 to AFP1 ratio measured in post mature seeds indicates high ABA levels and ABI5 activity also in the induction and maintenance of dormancy. ABA and DOG1 pathways converge on PP2C phosphatases such as AHG1 to suppress germination (Nee et al., 2017). DOG1 and the PP2Cs RDO5 and AHG1 were some of the most abundant measured proteins suggesting a possibly dominant role in seed dormancy by this pathway over ABA signaling. In addition two other PP2Cs (AT3G15260 and AT4G31860 of the F and I clades respectively) previously not known to be involved in seed development also accumulated to very high levels implicating them in the same processes. The abundance of RD26/ANAC72 (AT4G27410) and its two closest homologs ANAC19 (AT1G52890) and ANAC03 (AT3G15500) as well as two other NAC transcription factors, ANAC02 (AT3G15510) and ANAC14 (AT1G33060) increased specifically and significantly in seeds particularly in post mature seeds (brown siliques). *RD26* and homologs have been shown to be ABA responsive (Fujita et al., 2004) and to be expressed ubiquitously in *Arabidopsis* vegetative tissues in response to drought and salt stress (Tran et al., 2004; Nakashima et al., 2012) and coronatine (Zheng et al., 2012). They are also known to play a role in jasmonate (Bu et al., 2008) and brassinosteroid signaling (Ye et al., 2017) as well as leaf senescence (Takasaki et al., 2015). Given their role in promoting drought tolerance, their seed specific accumulation in the absence of stress may indicate a previously unknown role in the onset and maintenance of desiccation tolerance. No seed specific or reproductive phenotype has been reported for the respective single NAC gene knockout mutants possibly reflecting functional redundancy and to our knowledge no multiple knockout mutants have been produced.

### Ribosomal Proteins in Development

In our study we identified a large number of RPs and RBFs that all show the same tissue and developmentally specific pattern of protein abundance, increased in young (seedling) roots, young stem and early flowers/floral buds. The deep proteome coverage of our approach allowed us to identify entire RBF protein complexes, all of whose members had the same expression pattern. The RBF complexes were part of the SSU-processome or the pre-60S pre-ribosomal particle and mutation of many of their constituent RBFs lead to aberrant gametophyte development. This suggests that particularly these complexes regulate gametophyte development in early flowers in the context of ribosome biogenesis and translation, underscored by the their much more conserved, significant expression pattern as opposed to ribosomal proteins in general. It has been reported that the transcripts of several of these genes are especially abundant in developing tissue such as young roots, stem and flowers (Missbach et al., 2013). Furthermore, the extensive physical interaction between RPs and RBFs with the protein abundance pattern found here may indicate these particular isoforms assemble ribosomes specific to young roots, young stem and early flowers/floral buds with possible function specific to these tissue states. Closer inspection of the protein interactions may also bring to light new RBFs and RP/RBF complex conformations that have not yet been described. Indeed, although the complexes detected here are well described in yeast, much of what is known in *Arabidopsis* is by inference, so this study presents evidence of their tissue specific expression on the protein level, particularly pertaining to the CCT complex.

ABI5 is a positive regulator of ABA signaling that has been shown to repress the flowering transition (Wang et al., 2013) next to its well documented function in the induction and maintenance of seed dormancy (Finkelstein et al., 2008). So far a handful of proteins have been reported to target ABI5 to degradation and thus negatively impact the ABA response, including AFP1 (Lopez-Molina et al., 2003) and DWA2 (Lee et al., 2010). Both had distinct abundance patterns in this study. AFP1 was most abundant in ripe brown siliques (see above). DWA2 abundance was elevated in young roots, young stem and early flowers/floral buds wherein it presumably mediates ABI5 degradation to induce flowering. These results suggest different developmental and tissue specific mechanisms post-translationally control ABI5 levels and ABA activity.

### Vesicle Trafficking

Interestingly many of the membrane trafficking cluster 13 proteins were abundant in both fast growing tissues, primarily roots but also stem and flowers and senescent leaves. Vesicle trafficking is preeminent in the targeted deposition of new cell wall material in the clear zone at the apex of tip growing cells as is the formation of an apical actin structure. Autophagy, which also involves transport of cytoplasmic components in membrane vesicles is important in senescence, both in counteracting premature cell death and degradation of cellular structures such as the photosynthetic apparatus in nutrient remobilization. Our results at least suggest, that some of the molecular players in these distinct processes in different organs may be the same.

### Photosynthesis

The tissue and developmental abundance profile of cluster 11 proteins implicates them in chloroplast biogenesis and indeed, large sets of interacting proteins as well as known individual proteins central to its molecular processes were the major constituents of this cluster. These processes are known to be most prevalent in the young plant. The proteins of the photosynthetic apparatus are abundant in green tissues in nearly all developmental stages beginning in the cotyledons of the young seedling prior to photoautotrophy. This can be seen in the abundance profile of the proteins in cluster 8 which decreases in leaves in the later senescent stages, concomitantly with an increase of proteins that facilitate disassembly of the apparatus and pigment degradation in cluster 10.

### Senescence and Abscission

As such it is no surprise that a large number of antioxidant proteins and proteins involved in redox regulation and ROS scavenging increased in abundance as ageing progressed, peaking in senescence in leaves, cauline leaves and flowers/siliques. Chlorophyll and carotenoid catabolism are central aspects of leaf senescence and CYP89A9 is important in this respect catalyzing the oxidative deformylation of FCCs to dioxibilin-type FDCCs and ultimately the accumulation of NDCCs (Christ et al., 2013). *CYP89A2* has been shown to be co-expressed with *CYP89A9* (Obayashi et al., 2009) and we have observed this here for the cognate proteins. In addition to these two, 10 other CYP71 clan members had the same protein abundance pattern of cluster 10. This makes it likely, that a larger number of CYPs play undescribed roles in chlorophyll catabolism, as was also speculated by Christ and co-workers.

We detected 6 CRKs showing the characteristic abundance pattern associated with senescence. This RLK gene family consists of 44 members all in close proximity on chromosome 4 with a host of different physiological functions (Wrzaczek et al., 2010; Burdiak et al., 2015). Their extracellular regions have two conserved DUF domains each with 4 cysteine residues as potential targets for thiol redox regulation. Members of the gene family have been shown to be expressed on the transcript and protein level in response to pathogen challenge, flg22 and SA, underpinning a potential role in immunity (Wrzaczek et al., 2010; Yadeta et al., 2017). Transcripts of five of the six CRKs we identified were also significantly upregulated following ozone treatment (Wrzaczek et al., 2010). Collectively these processes as well as senescence all involve ROS, although the spatio-temporal mechanisms of ROS production, interaction and signaling in the senescence syndrome are not well understood. A probable ROS dependent role of CRK5 in cell death and senescence is documented (Burdiak et al., 2015). Our results further support an as of yet little known role of the CRKs in senescence. Considering CRKs as ROS sensors and signaling molecules leads to the question of chloroplast derived ROS production affecting the extra-cellular redox state akin to the oxidative burst upon pathogen perception.

*Arabidopsis* floral organ abscission as the culmination of petal senescence and nutrient remobilization for fruit ripening and ultimately release is a well-studied model of general abscission processes (Patharkar and Walker, 2018). Here we detected most of the components of the abscission signaling cascade increasing in abundance in flowers as they age and set seeds in siliques following fertilization. Recently it has been shown that the same mechanisms also act in *Arabidopsis* cauline leaf abscission in response to drought (Patharkar and Walker, 2016) and indeed it is common that plants shed their leaves in response to various environmental stimuli. However, leaf abscission is not known to be on a developmental clock and *Arabidopsis* does not abscise rosette leaves (Stenvik et al., 2006), making it intriguing that we also found the abundance of the proteins to increase in both rosette and cauline leaves in the course of ageing and senescence.

ABA plays a major role in fruit ripening (Jia et al., 2011) and the leaf senescence syndrome (Song et al., 2016), yet its role in abscission is debated. ABA promotes ethylene biosynthesis during the later stages of fruit ripening in tomato (Zhang et al., 2009) and ethylene is known to be a regulator of abscission in *Arabidopsis* (Patterson and Bleecker, 2004). The hormone is also a key factor in the response to drought (Huang et al., 2008). *HAE*, *HLS2* and *IDA* expression in *Arabidopsis* seedlings is induced by ABA and to a lesser extent ethylene (eFP browser (Goda et al., 2008)). It is possible, that the elevated ABA and ethylene levels in the course of senescence led to the accumulation of the abscission signaling proteins in rosette and cauline leaves. Therefore one may speculate on possible other yet unknown functions of this signaling module outside of organ abscission.

### Photosynthesis and Primary Metabolism in PTI

It is known that pathogen infection leads to inhibition of photosynthesis and it has been shown that this is an active response of the plant to the invading pathogen. Downregulation of a host of genes related to the light dependent reaction and particularly photosystem II and parameters of photosynthetic activity has recently been reported to be dependent on constitutive MPK3/MPK6 activation in ETI (Su et al., 2018). The resulting increase in chloroplast localized ROS is hypothesized to support the programmed cell death in the hypersensitive response (HR). Measurement of the photosynthetic parameters upon exposure to 100 nM flg22 for up to 24 hours or infiltration with a *Pseudomonas syringae pv tomato DC3000* (*Pst* DC3000) strain that lacks a type III secretion system to deliver effectors in this study indicated that downregulation of photosynthesis does not occur in PTI, which feature only transient MPK3/MPK6 activation. An older proteomics study however discloses downregulation of some photosynthetic proteins and non-photochemical quenching (NPQ) 2 hours after flg22 treatment (100 nM to 10 µM) and decrease of electron flux through PSII upon 7 days of exposure, suggesting inhibition of photosynthesis may occur in some PTI scenarios (Gohre et al., 2012). This is in line with our findings where plants were exposed to 1 µM flg22 for sixteen hours. Conceivably higher flg22 doses may lead to prolonged MPK3/MPK6 activation and photosynthetic inhibition also in PTI. As an afterthought, the abundance of both MPK3 and MPK6 increased somewhat in the discovery proteomics results (Supplemental table 14) upon PAMP exposure although we did not measure phosphorylation levels.

### Jasmonate and Salicylic Acid Cross Talk

The induction of PTI downstream of FLS2 binding flg22 is a commonly accepted model of bacteria induced basal immunity in plants. As the majority of bacteria adopt biotrophic lifestyles, it can also be considered a model of plant resistance to biotrophs although not exclusively, as some bacteria, such as *Erwinia carotovora* are necrotophs. Also bacteria will be recognized by more than one pattern recognition receptor (PRR) such as EF-TU RECEPTOR (EFR) and others in addition to FLS2, inducing multiple partially overlapping responses, so the flg22/FLS2 model may be overly specific and somewhat artificial.

The importance of SA and JA in resistance to biotrophs and necrotrophs respectively as well as their generally antagonistic modes of action are well documented (Pieterse et al., 2012). It has also been shown that both hormones interact and play roles in flg22 induced PTI (Denoux et al., 2008; Yi et al., 2014; Hillmer et al., 2017; Mine et al., 2017). Our results suggests flg22 induced PTI at the steady state of 16 hours after continuous flg22 exposure prioritizes SA mediated defenses over JA mediated defenses because of dampening of JA and JA-Ile accumulation in culture grown seedlings. SA has been shown to accumulate in and be essential for flg22 triggered defenses (Tsuda et al., 2008). The suppressive effect of SA on JA levels has been extensively explored and shown to be NPR1 dependent in *Pst* DC3000 infected *Arabidopsis* plants (Spoel et al., 2003) and integral to the trade-off between defense against biotrophic and necrotrophic pathogens (Spoel et al., 2007). However most of this interplay is downstream of JA synthesis at the level of inhibition of JA responsive gene transcription by transcriptional regulators primarily WRKY70 and TGAs and ORA59 (Li et al., 2004; Leon-Reyes et al., 2010; Shim et al., 2013; Van der Does et al., 2013; Zander et al., 2014). Here we propose a model wherein a MYC2 dependent negative feed-back loop in equilibrium with JA synthesis controls JA-Ile and JA levels via clearance of the phytohormones by deconjugation and hydroxylation by IAR3 and JOX2 respectively. Four major arguments for this model can be extrapolated from our measurements.

1. JA and JA-Ile levels were very low and did not increase significantly following flg22 exposure in the wild type. However, the abundance of all of the proteins in the JA synthesis pathway and of the OPR3 transcript was significantly increased after induction of PTI (16 and 3 hours respectively). It has been reported, that JA biosynthesis and signaling gene expression was upregulated in *Arabidopsis* at earlier time points (1h and 3h) following flg22 exposure (Denoux et al., 2008). The OPR3 transcript levels were again basal 16 hours post flg22 perception whereas protein abundance was still elevated, highlighting the importance of measuring protein levels directly.
2. Absolute levels of the JA intermediate OPDA were very high (µg/g FW) and increased more than 4-fold following treatment with the PAMP in both the wild type and the *myc234* background.
3. JA and JA-Ile levels were strongly elevated in the *myc234* mutant as opposed to the wild type. This indicates suppression of the phytohormone and its bioactive conjugate is dependent on MYC2 in PTI. Moreover, this result together with point 3. above leads us to doubt that arrest of the JA synthesis pathway at the point of OPDA synthesis is the reason for the basal JA and JA-Ile levels. Also OPDA REDUCTASE 3 (OPR3) which reduces OPDA was the pathway component with the greatest fold change in abundance.
4. The amidohydrolase IAR3 and the 2OG oxygenase JOX2 were among the proteins with the most significantly increased fold change in abundance following 16 hours of flg22 exposure. The same goes for the JOX2 transcript after three hours. MYC2 binding sites were identified as the top scoring motifs in *in silico* analysis of both the JOX2 and IAR3 promotor regions. These proteins deconjugate JA-Ile and hydroxylate free JA respectively, thereby inactivating the former as well as depleting both of them.

These results lead us to hypothesize that deconjugation and subsequent modification of JA-Ile play an important role in control of the JA pathway in defense against biotrophs as has been shown against the necrotrophic fungus *Botrytis cinerea* (Caarls et al., 2017). JA/JA-Ile clearance is presumably in some type of equilibrium with JA synthesis because it has been reported that the JA pathway is party to PTI induction and SA accumulation via EDS5 (Mine et al., 2017). The involvement of JA signaling in SA accumulation is underscored in our data by the diminished SA levels in the *myc234* background at the earlier time points (1h and 3h after flg22 exposure). Thus our model reconciles significant induction of the JA synthesis pathway proteins and known SA JA antagonism at the level of transcriptional regulation with the low observed JA and JA-Ile levels we measured and which were reported previously (Spoel et al., 2003).

### Auxin/IAA homeostasis

Auxins are among the most studied phytohormones and next to their preeminent role in growth and development have established functions in plant immunity. Free IAA and IAA signaling both increase plant susceptibility to biotrophic pathogens (Kunkel and Harper, 2018) whereas IAA acts synergistically with JA in resistance to necrotrophic pathogens (Qi et al., 2012). Flg22 induces transcription of a microRNA miR393 that targets the auxin receptor TRANSPORT INHIBITOR RESPONSE 1 (TIR1), thereby dampening auxin signaling (Navarro et al., 2006). Free IAA levels were not reported to change dramatically in response to flg22 (Navarro et al., 2006) nor to *PStDC3000* infection or SA (Qi NewPhytol 2012) and we also observed a slight but significant decrease after 16 hours of exposure, presumably by downregulation of the IAM pathway indicated by a decrease in the abundance of AMIDASE1 (AMI1) when steady state PTI is reached. IAA is synthesized primarily from tryptophan by way of several biosynthesis pathways and tryptophan also channels into defense compound synthesis, especially indolic glucosinolates and camalexin. Thus branching points in tryptophan/IAA biosynthesis pathways, particularly indole-3-acetaldoxime (IAOx) (Sugawara et al., 2009) may represent an important nexus in the growth defense trade off. Here we measured substantial, significant increase in protein abundance in the entire tryptophan as well as in defense compound biosynthesis pathways in response to flg22. The abundance of enzymes (AAO1, NIT1 and NIT4) in some alternative auxin synthesis pathways with known secondary roles in defense also increased.

It is known that JA induces these pathways upon *Alternaria brassicicola* infection also leading to increase in free IAA levels in the response to necrotrophic pathogens (Qi et al., 2012). In our study JA and JA-Ile levels remained basal. However, we measured a significant increase in the abundance of ethylene biosynthesis proteins ACC OXIDASE 2 (ACO2) and 1-AMINO-CYCLOPROPANE-1-CARBOXYLATE SYNTHASE 7 (ACS7) and a significant, more than 5-fold increase of ET 16 hours after PAMP exposure. Clay and co-workers reported expression of tryptophan and IG biosynthesis pathway transcripts to be mediated by ET via myb transcription factors. These pathways play important roles in defense against both biotrophic and necrotrophic pathogens and our results suggest that they are induced independently by different phytohormones, ET and JA respectively. These two hormones are generally synergistic posing new questions about the role of JA in the flg22 response.

Auxins are unique among phytohormones in that they are transported directionally from cell to cell by a polar transport system with many components (Friml, 2003). Indeed, the distribution and local IAA concentration have a profound impact on cellular processes more so than absolute IAA levels or aspects of synthesis and catabolism (Teale et al., 2006) and auxin transport mutants have been reported to be defective in mounting SAR (Truman et al., 2010b). Concurrently, we did not observe major changes in free IAA levels, however the abundance of two of the most well studied auxin efflux carriers, PIN3 and PIN7, decreased and that of a protein directing the localization of the influx carrier AUX1 increased after flg22 exposure. This suggests that IAA transport and local IAA gradients also play an important role in the defense against biotrophic in addition to necrotrophic pathogens (Qi et al., 2012).

### Methods Summary

For detailed description of methods and optimization see Supplemental Methods and Data. In brief, proteins were extracted from *Arabidopsis thaliana* suspension cultures or soil grown plants and tissues with 4% SDS, separated into 5 bands by SDS-PAGE, in gel digested and measured on an Orbitrap Velos Pro mass spectrometer with a conventional DDA scan strategy using a 50 cm C18 liquid chromatography column and an extended gradient of 9 hours. MS data was searched using the Mascot and Andromeda search engines. Search results were concatenated and peptides and proteins identified with global FDR thresholds of 0.01 in the Scaffold software. Proteins were quantified by way of PSM counting. For parallel reaction monitoring (PRM) proteins were extracted with 4% SDS, digested with an optimized FASP protocol and peptides measured on a QExactive Plus mass spectrometer. Up to three proteotypic peptide m/z per protein were targeted, peptides and proteins were identified using the Mascot search engine and data analysis and AUC quantification of 6 fragment ions was done using the Skyline software. Statistical significance of protein fold changes was inferred at a threshold of 0.05 if one of the quantified peptides tested significant. qPCR was performed with the EvaGreen Kit from Bio&Sell according to manufacturer’s instructions, phytohormone and amino acid measurements according to (Ziegler et al., 2014). Bioinformatics analysis including gene ontology analysis and pathway visualization and mapping was done with DAVID, MapMan, STRING; PTM analysis with MSFragger. Multivariate data analysis was done in case of fuzzy c-means clustering with the m-fuzz R package, hierarchical clustering and PCA with Perseus. Co-expression of proteins from neighboring genes was analyzed using an in-house program programmed in R.

## Supporting information

Bassal Supplemental Methods and Data

## Supplemental Figure Legends

**Supplemental figure 1.**
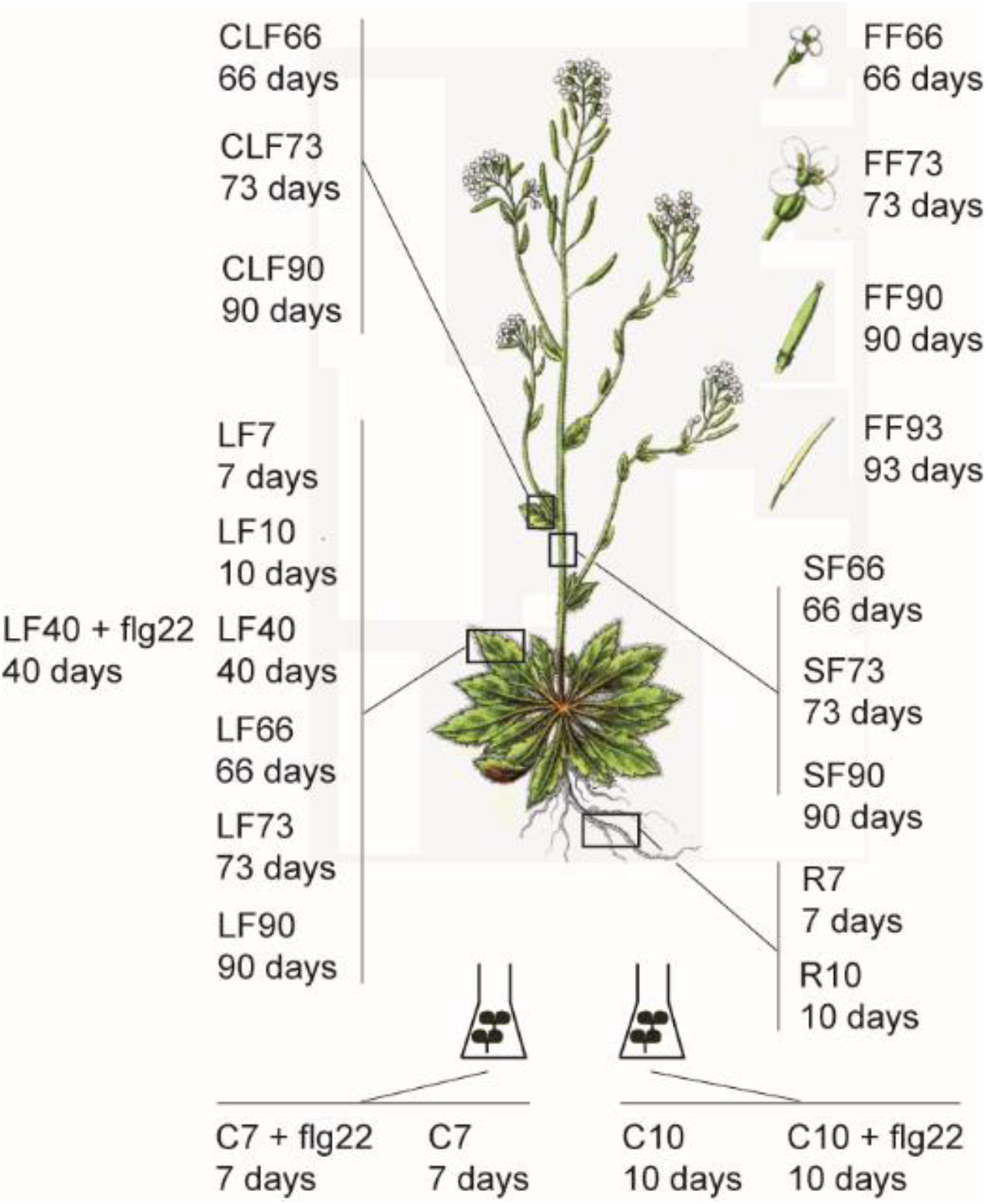
Sampling of different *Arabidopsis thaliana* tissues and seedlings at various stages in the plants life (in days). Samples correspond to Supplemental methods and data tables 1 and 2.

**Supplemental figure 2.**
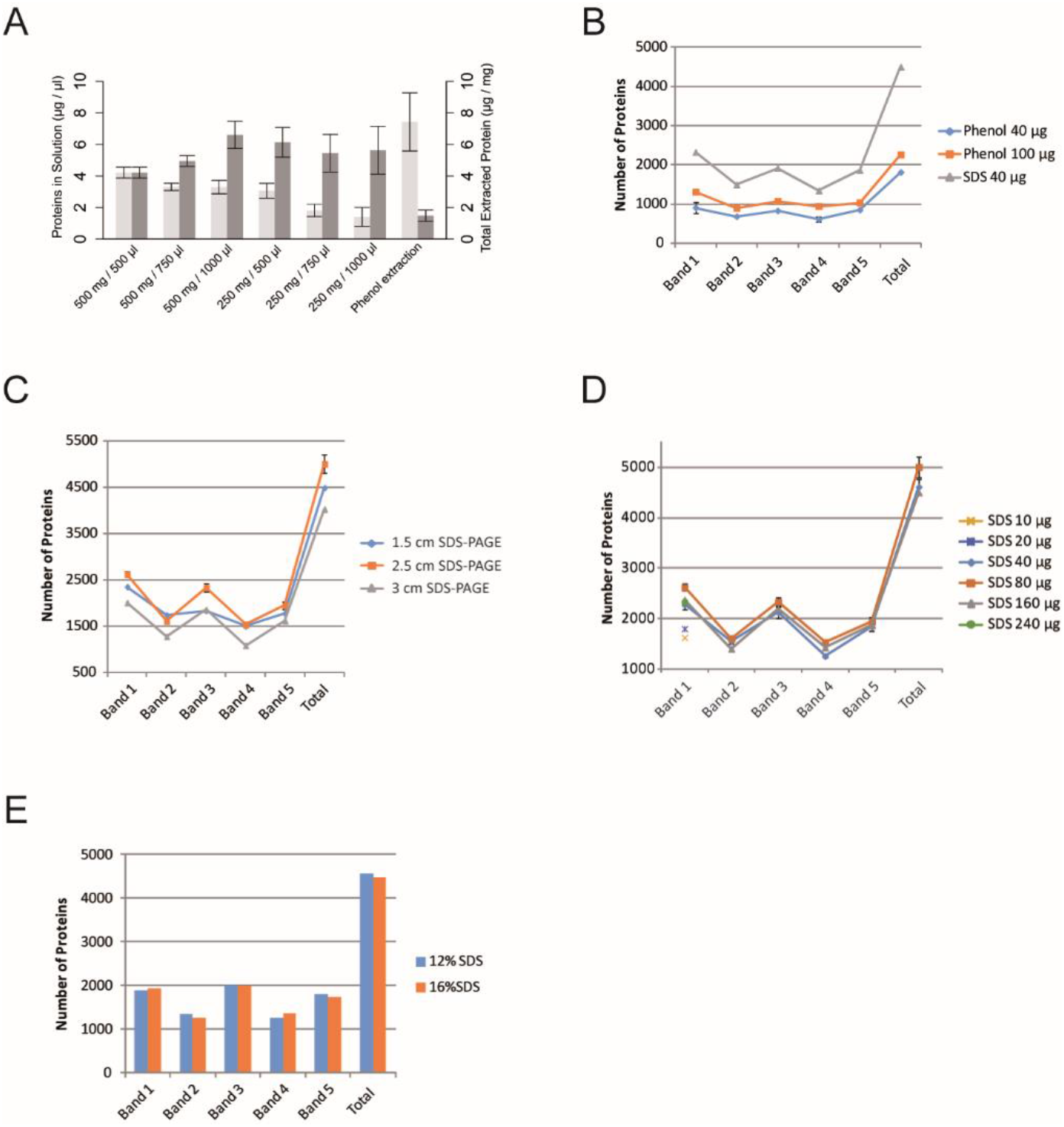
Experiments to optimize GeLC-MS approach for deep proteomics analysis of plant tissues A. Total amount of extracted proteins and concentration of proteins in solution for various tissue amounts and tissue to buffer ratios using the SDS based protein extraction described in this work. For 250 mg / 750 µl and 250 mg / 1000 µl mean values and standard deviations (error bars) of four experiments are shown, for 500 mg / 500 µl, 500 mg / 750 µl and 250 mg / 500 µl of eight and for 500 mg / 1000 µl of 18 experiments are shown. For phenol extraction (500 mg / 1000 µl) mean values and standard deviations of six experiments are shown. B. Number of proteins identified (Mascot, no significance filters) in bands 1 through 5 and in total (cumulative sum of all bands) employing a 180 minute LC gradient when loading different amounts of phenol extracted proteins onto the SDS-PAGE as compared to 40 µg of SDS extracted proteins. Mean values and standard errors of 2 experiments are shown for Phenol 40 µg C. Number of proteins identified (Mascot, no significance filters) in bands 1 through 5 and in total (cumulative sum of all bands) employing a 180 minute LC gradient when separating 80 µg of SDS extracted proteins over different distances in SDS-PAGE. Mean values and standard errors of 2 experiments are shown for 2.5 cm SDS-PAGE. D. Number of proteins identified (Mascot, no significance filters) in bands 1 through 5 and in total (cumulative sum of all bands) employing a 180 minute LC gradient when separating different amounts of SDS extracted proteins for 2.5 cm on SDS-PAGE. Mean values and standard errors of 3 experiments are shown for 40 µg and 80 µg of SDS extracted proteins. Only band 1 was measured for 10 µg, 20 µg and 240 µg of SDS extracted proteins. E. Number of proteins identified (Mascot, no significance filters) in bands 1 through 5 and in total (cumulative sum of all bands) employing a 180 minute LC gradient when separating 80 µg of SDS extracted proteins for 2.5 cm on 12% and 16% SDS-PAGE.

**Supplemental figure 3.**
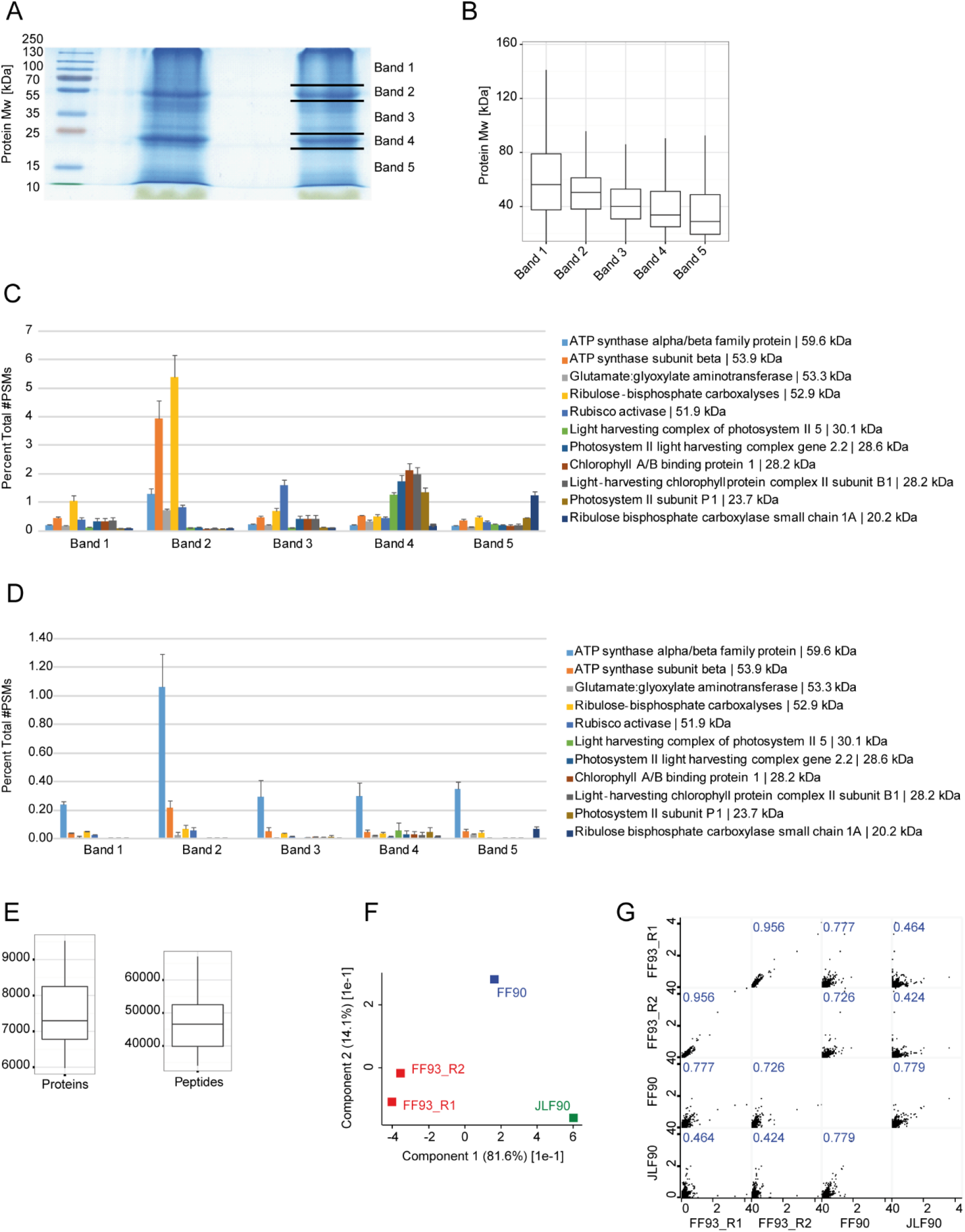
GeLC-MS approach for deep proteomics analysis of plant tissues A. SDS extracted plant protein extract is separated on a 12% SDS-PAGE for a total distance of exactly 2.5 cm. High staining intensity bands are excised and analyzed separately leading to 5 protein fractions. Contaminants such as secondary metabolites, chromophoric compounds and leaf pigments (colored green) precede the proteins and can be readily eliminated. B. Proteins are fractionated in the gel bands according to their size. Box plots show molecular weight (Mw) of all proteins identified with at least one unique peptide and a peptide FDR threshold (q-value) < 1% with the Mascot software linked to Proteome Discoverer from 8 green tissue samples (LF7, LF10, LF40, LF66, LF73, JLF66, JLF73, JLF90; see Supplementary Methods and Data for explanation) in the respective bands. Boxes represent the inter-quartile range (IQR) between the first and third quartile. Whiskers extend up to 1.5*IQR plus the third quartile and down to first quartile minus 1.5*IQR. C. Highly abundant leaf proteins involved in photosynthesis are fractionated. Mean number of peptide spectral matches (#PSMs) identified as above for the 8 green tissues samples as above are plotted for each of the indicated proteins as a percentile of the total PSMs of all proteins per band. Error bars indicate standard errors (SE). D. As C. for two root samples R7 and R10 (see Supplementary Methods and Data for explanation). E. Between 6,000 and 9,000 proteins and 35,000 and 65,000 unique peptides were identified with a peptide and protein q-value < 1% with the Mascot and MaxQuant software integrated in the Scaffold software per tissue sample. Box plots show protein and peptide numbers from all 23 samples; otherwise as above. F. Protein identification and quantification is repeatable. Principal component analysis of two repeated GeLC-MS analyses of flowering stage 4 sample (FF4) and FF3 and JLF3 samples. The two principal components capture more than 96% of the total variance indicating the two dimensional subspace almost perfectly represents the relationship of the higher dimensional samples. G. Protein identification and quantification is repeatable. Pair wise scatter plots of the four samples in F. Pearson correlation coefficients are given.

**Supplemental figure 4.**
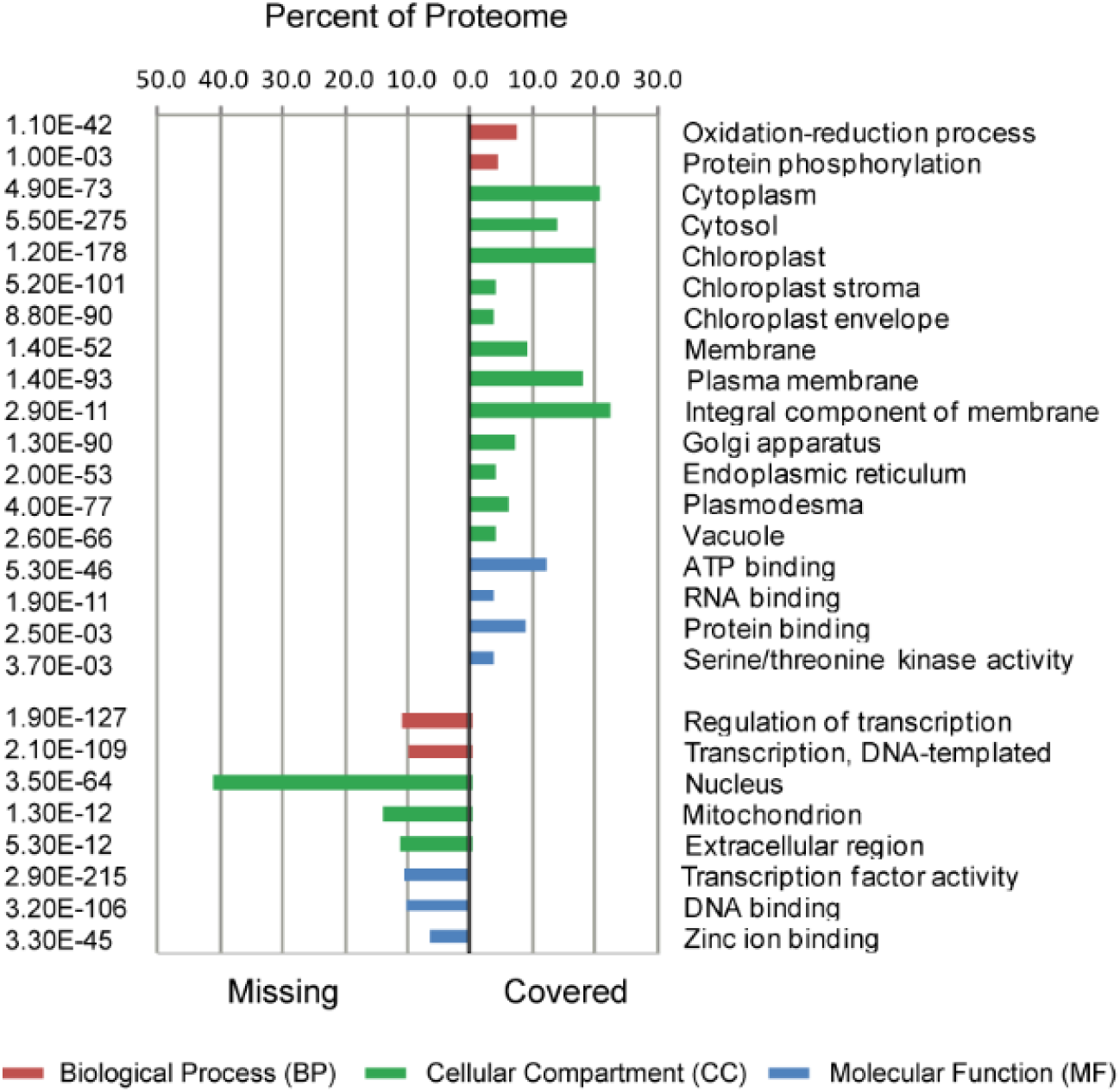
DAVID Missing and Covered GO terms. Gene ontology classification of all proteins identified in the study, the “covered” Arabidopsis proteins and of all remaining protein coding genes for which no cognate peptide was identified, the “missing” proteome. The percentage of the respective proteome is given for each GO bin as well as the Benjamini corrected p-value for enrichment.

**Supplemental figure 5.**
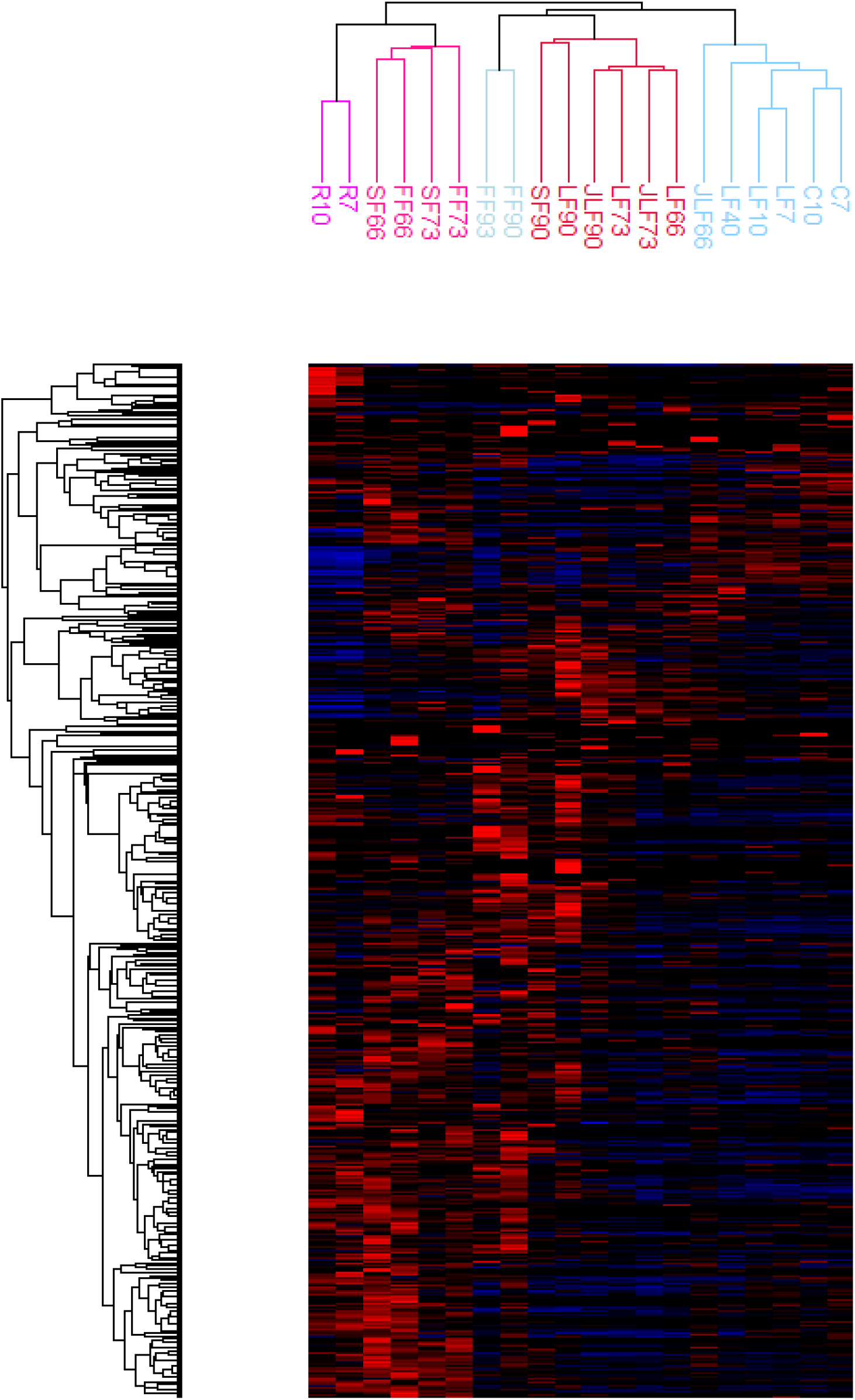
Hierarchical cluster analysis of the deep proteomics measurements of sampled tissues. Row tree was generated using Spearman correlation, column tree using Pearson correlation. Values were z-score transformed so color gradient goes from -3 (dark blue) to 0 (mean value; black) to 4 (bright red).

**Supplemental figure 6.**
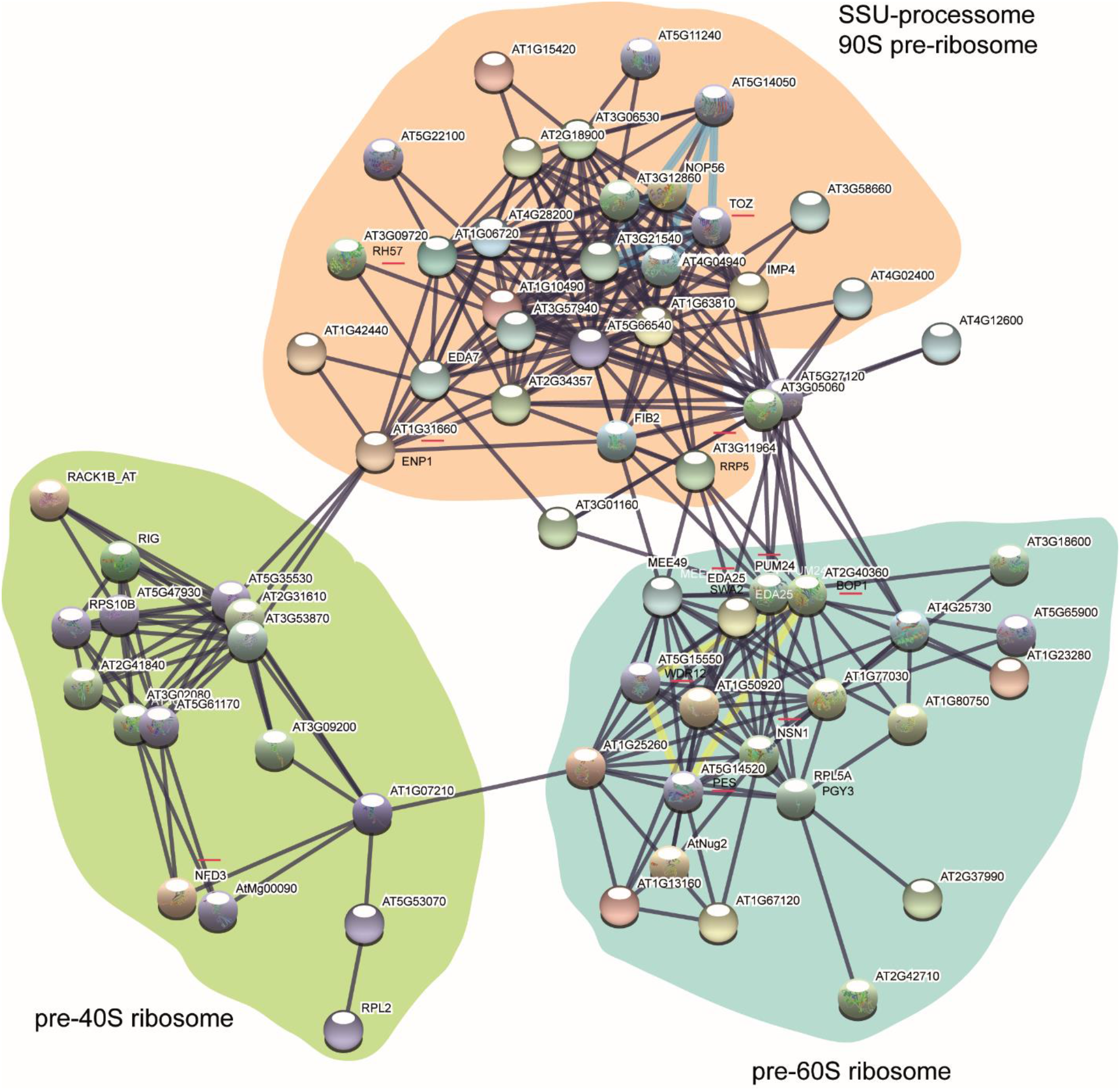
All cluster 4 proteins STRING physical interactions. Physical protein interaction networks produced with STRING database of all cluster 4 proteins. Thick blue edges highlight the UTP-B, t-UTP complexes, thick yellow edges the PeBoW complex. Red underlines indicate gene deletion has a developmental phenotype. Ribosomal core complexes are color coded.

**Supplemental figure 7.**
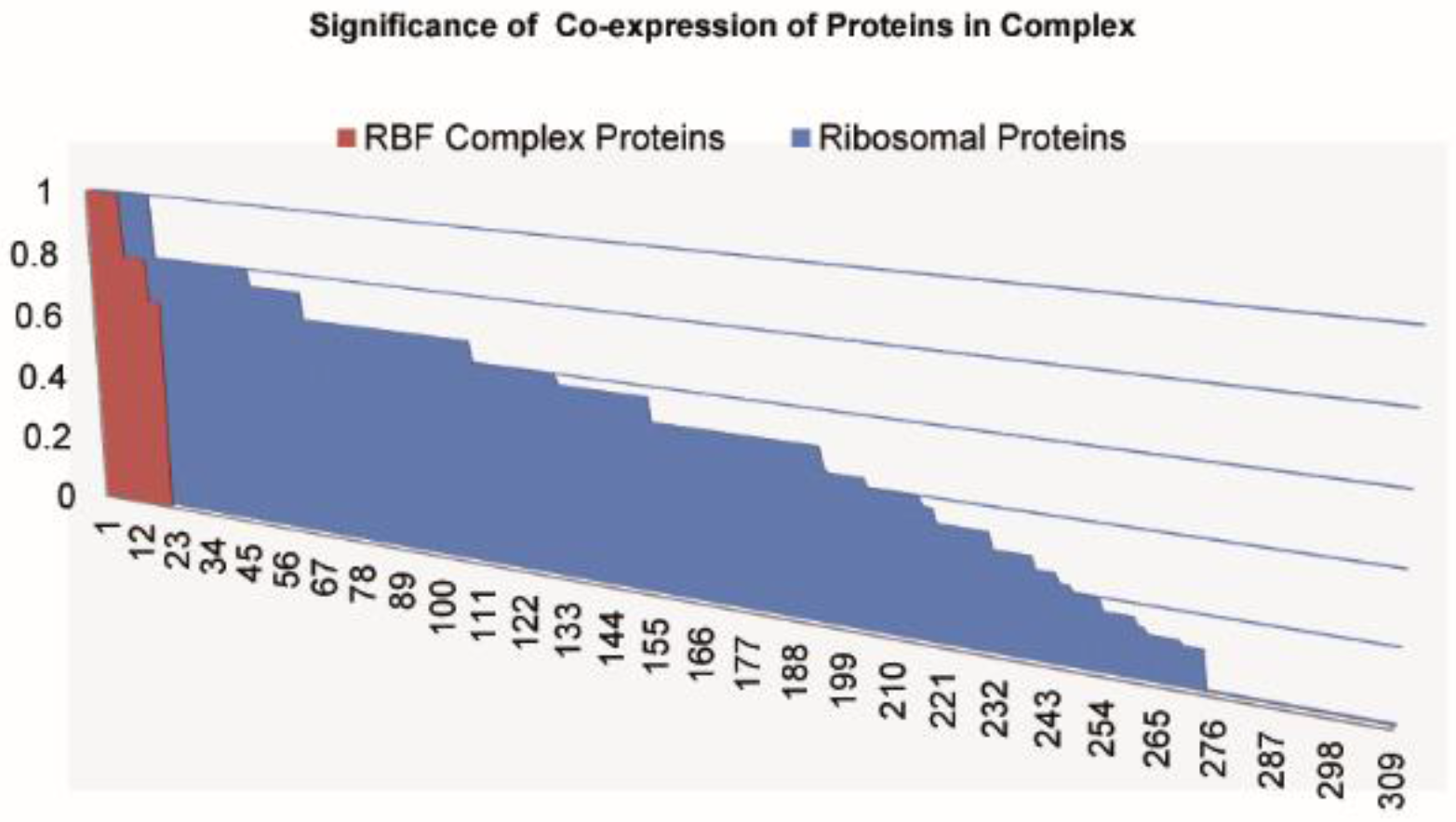
Significance of RBF complex protein co-expression. A 95% confidence interval was calculated for the mean expression value of RBF complex proteins in cluster 4 and all ribosomal proteins and ribosome constituents as background. To assess the significance of RBF protein tissue and developmental specific expression according to the cluster 4 expression pattern, the number of times the expression value for samples R7, R10, SF66 and FF66 were outside the interval were counted for each protein and divided by the number of expression values of all samples that were outside of the interval. A ratio of 1 indicates only the 4 samples mentioned above were significantly removed from the mean, indicating 95% confidence in protein expression specific to these tissues and developmental stages, as found for many of the RBFs (red bars). Ribosomal proteins in blue had much lower ratios indicating expression values in more diverse tissues and developmental stages were significantly removed from the mean and therefore alternate and non-specific expression patterns.

**Supplemental figure 8.**
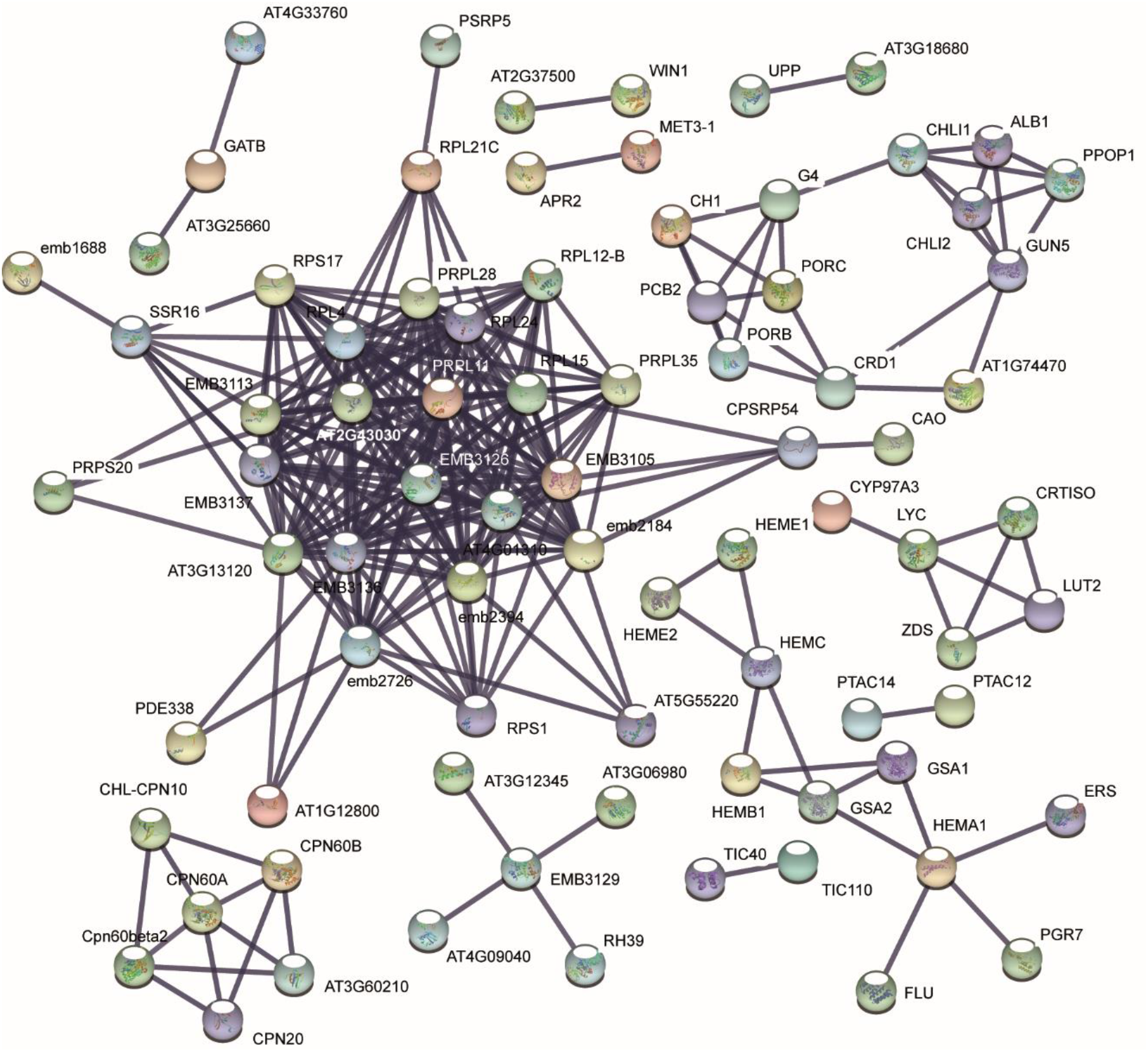
Cluster 11 proteins Chloroplast. Complete physical / functional protein interaction network generated with the STRING database using all proteins assigned to the GOTERM category chloroplast (Supplemental table 11) as input set. Proteins are labelled.

**Supplemental figure 9.**
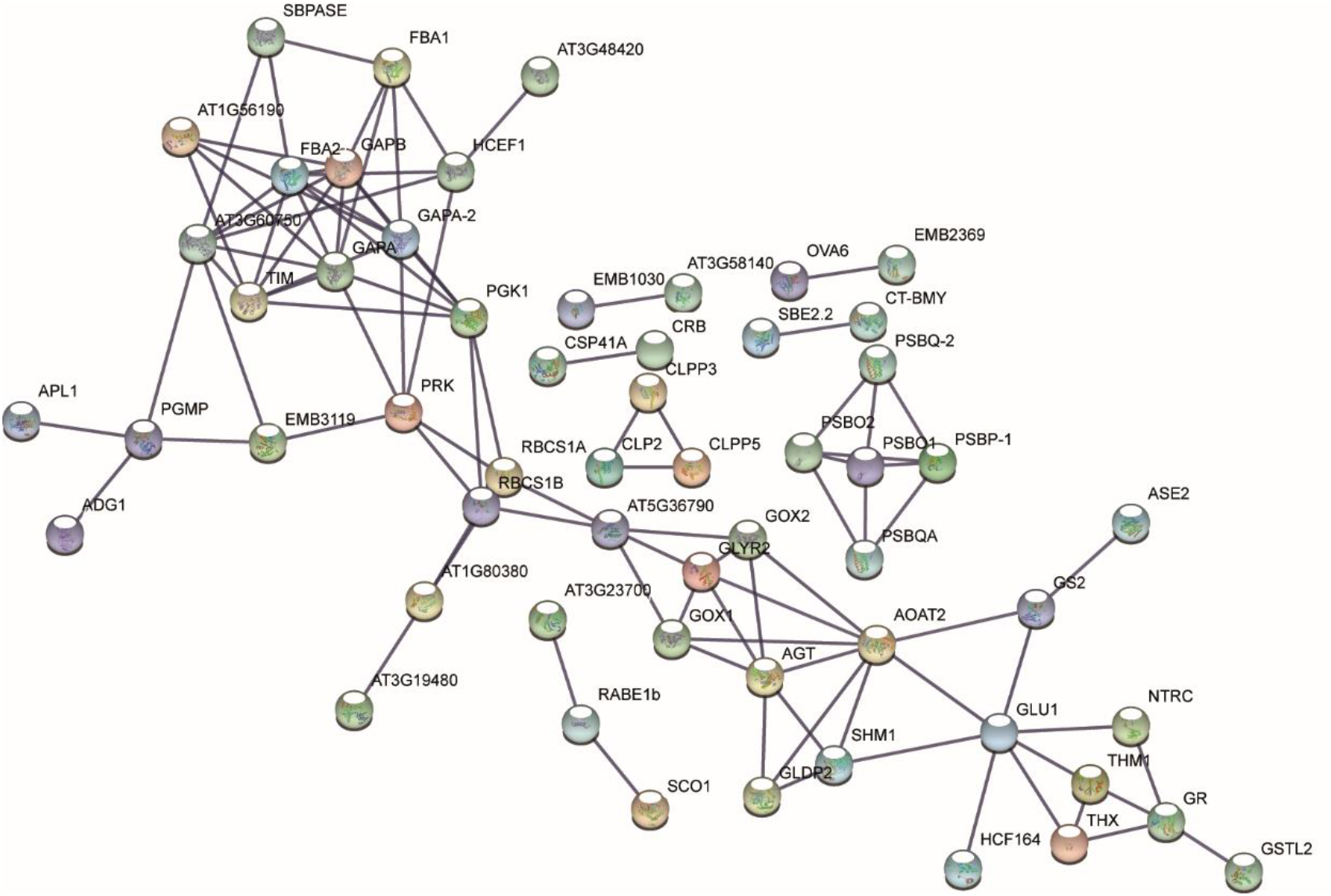
Cluster 8 proteins Stroma. Complete physical / functional protein interaction network generated with the STRING database using all proteins assigned to the GOTERM category Stroma (Supplemental table 12) as input set. Proteins are labelled.

**Supplemental figure 10.**
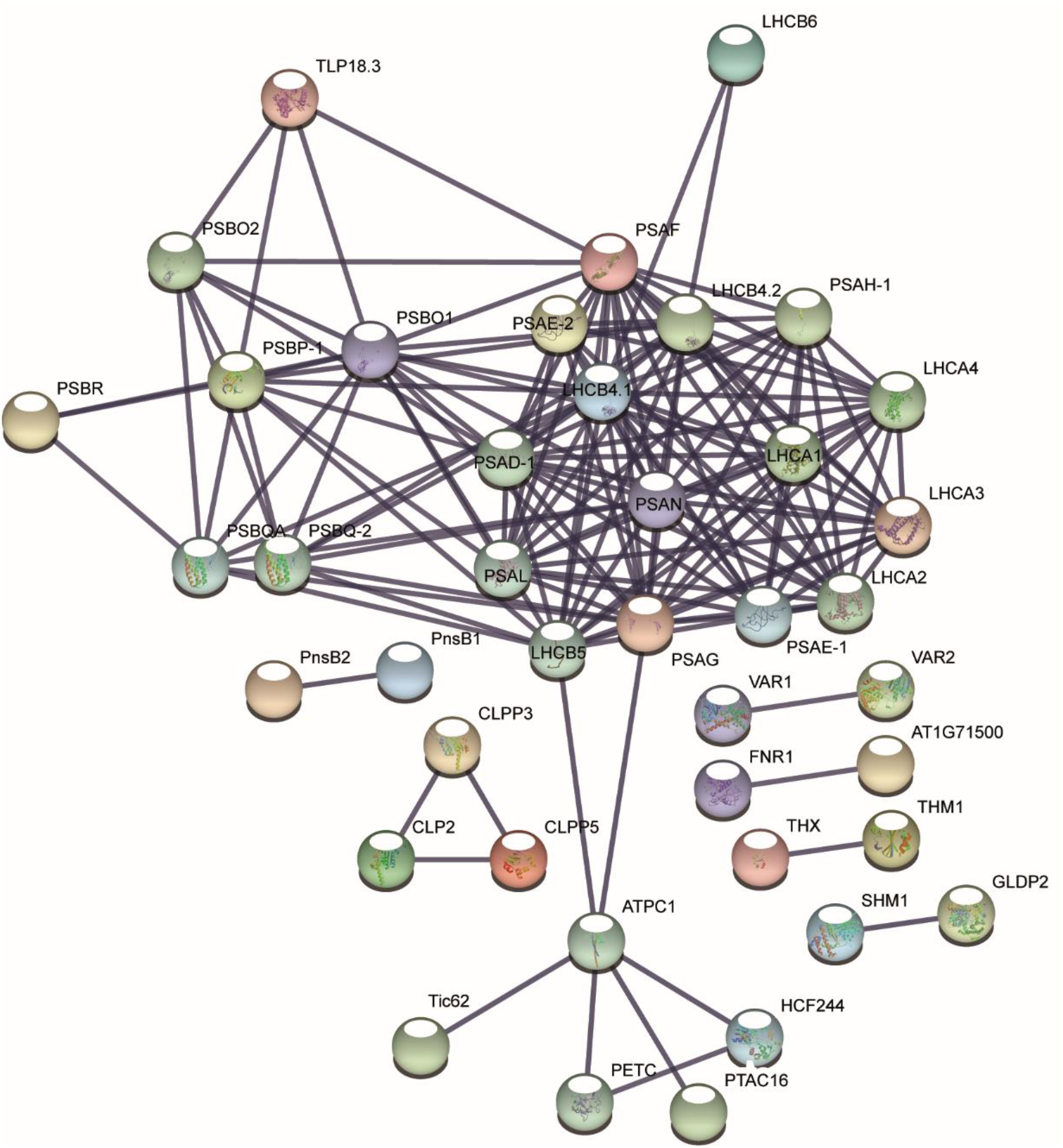
Cluster 8 proteins Thylakoid. Complete physical / functional protein interaction network generated with the STRING database using all proteins assigned to the GOTERM category Thylakoid (Supplemental table 12) as input set. Proteins are labelled.

**Supplemental figure 11.**
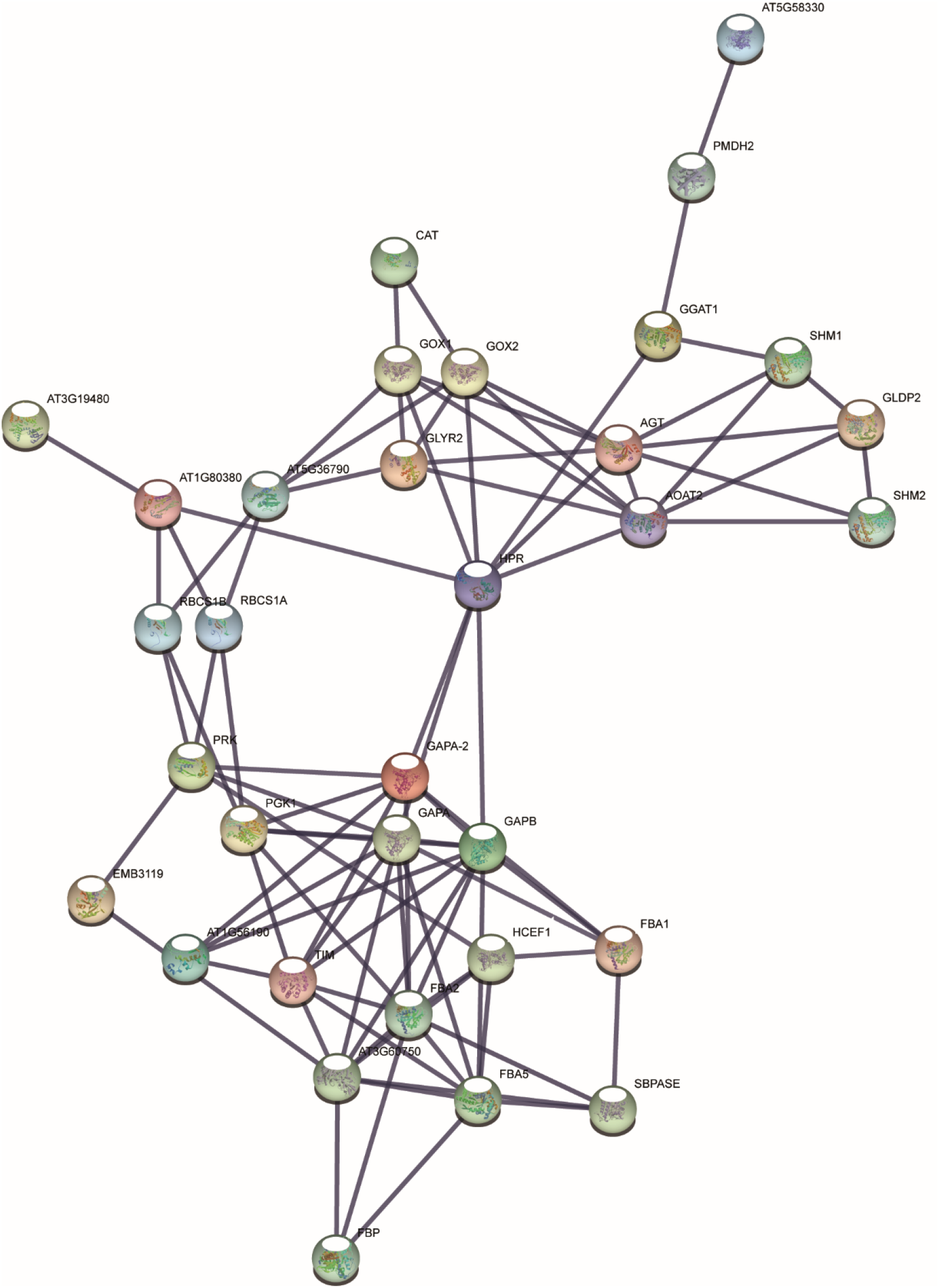
Cluster 8 proteins Carbon fixation. Complete physical / functional protein interaction network generated with the STRING database using all proteins assigned to the GOTERM category Carbon metabolism (C Metabolism) (Supplemental table 12) as input set. Proteins are labelled.

**Supplemental figure 12.**
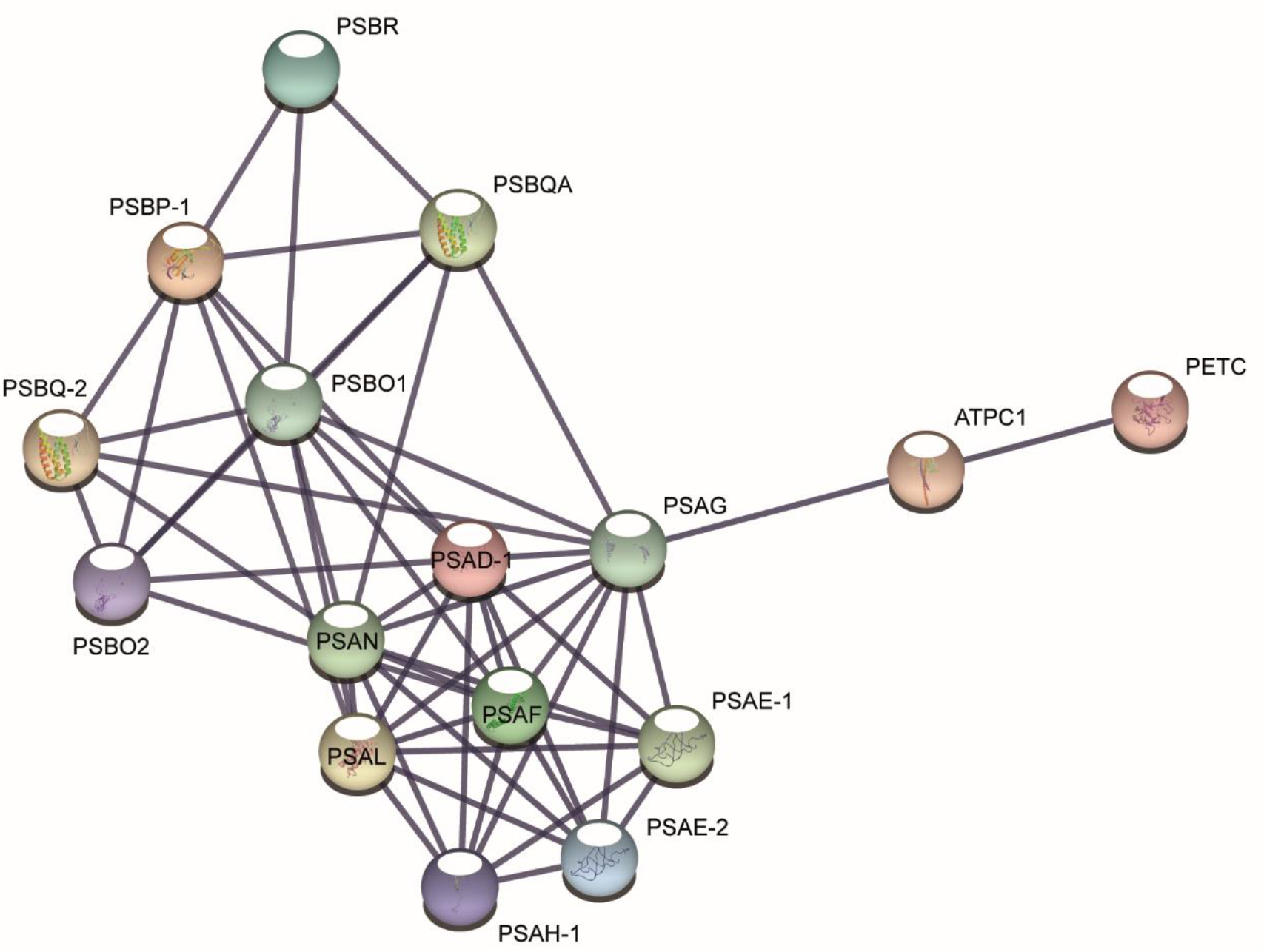
Cluster 8 proteins Stroma. Complete physical / functional protein interaction network generated with the STRING database using all proteins assigned to the GOTERM category Photosynthesis (Supplemental table 12) as input set. Proteins are labelled.

**Supplemental figure 13.**
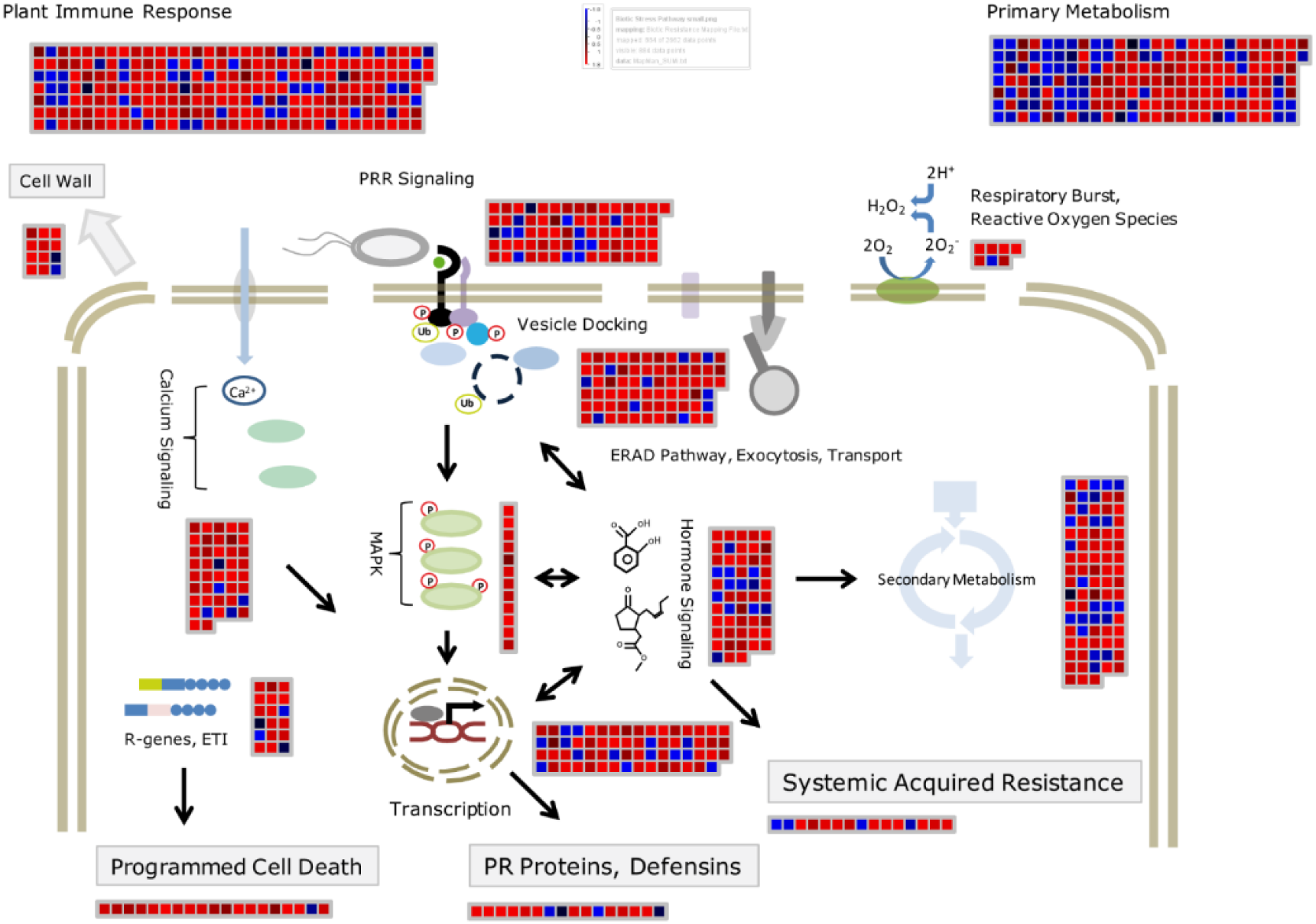
Flg22 MapMan. The MapMan software was used to characterize and map proteins with changes in their abundance following 16 hours of flg22 treatment to various avenues of plant immunity. The PTI pathway was drawn by ourselves and populated with a mapping concatenating the the MapMan Affymetrix mapping depicted on the pathway and related to PTI with corresponding GO terms from the TAIR GO slim ontology. Protein abundance is shown as the sum of z-score transformed spectral counts acquired for the respective proteins in measurements of flg22 treated 7 and 10 day old liquid culture grown seedlings. Red indicates an increase in abundance blue a decrease (black is no change). Max and minimum values in the color bar are 1.8 and – 1.8 respectively.

**Supplemental figure 14.**
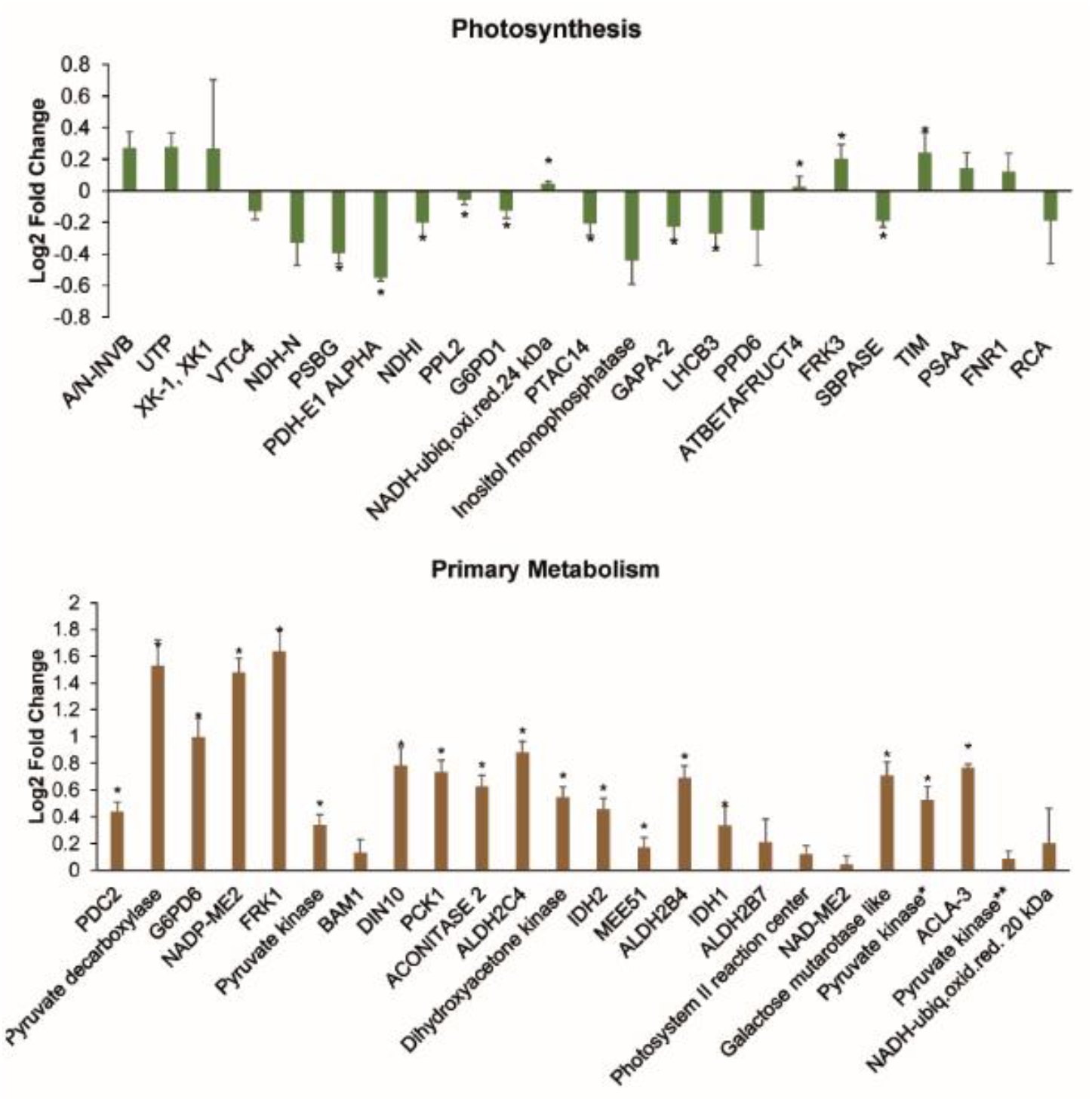
Targeted PRM based quantification of proteins involved in photosynthesis and primary metabolism. Bars represent log2 fold changes of protein abundance after 16 hours of flg22 exposure (1µM concentration in medium) estimated by area under the curve label free protein quantification index (PQI) of the 6 most intense product ions from MS2 spectra of targeted proteotypic peptides. Bars represent median PQI of all quantified proteotypic peptides for a given protein in 9 measurements (3 biological replicates each measured 3 times). Standard error is indicated. Star indicates significance α=0.05 if fold change of at least one of the quantified peptides was significant.

**Supplemental figure 15.**
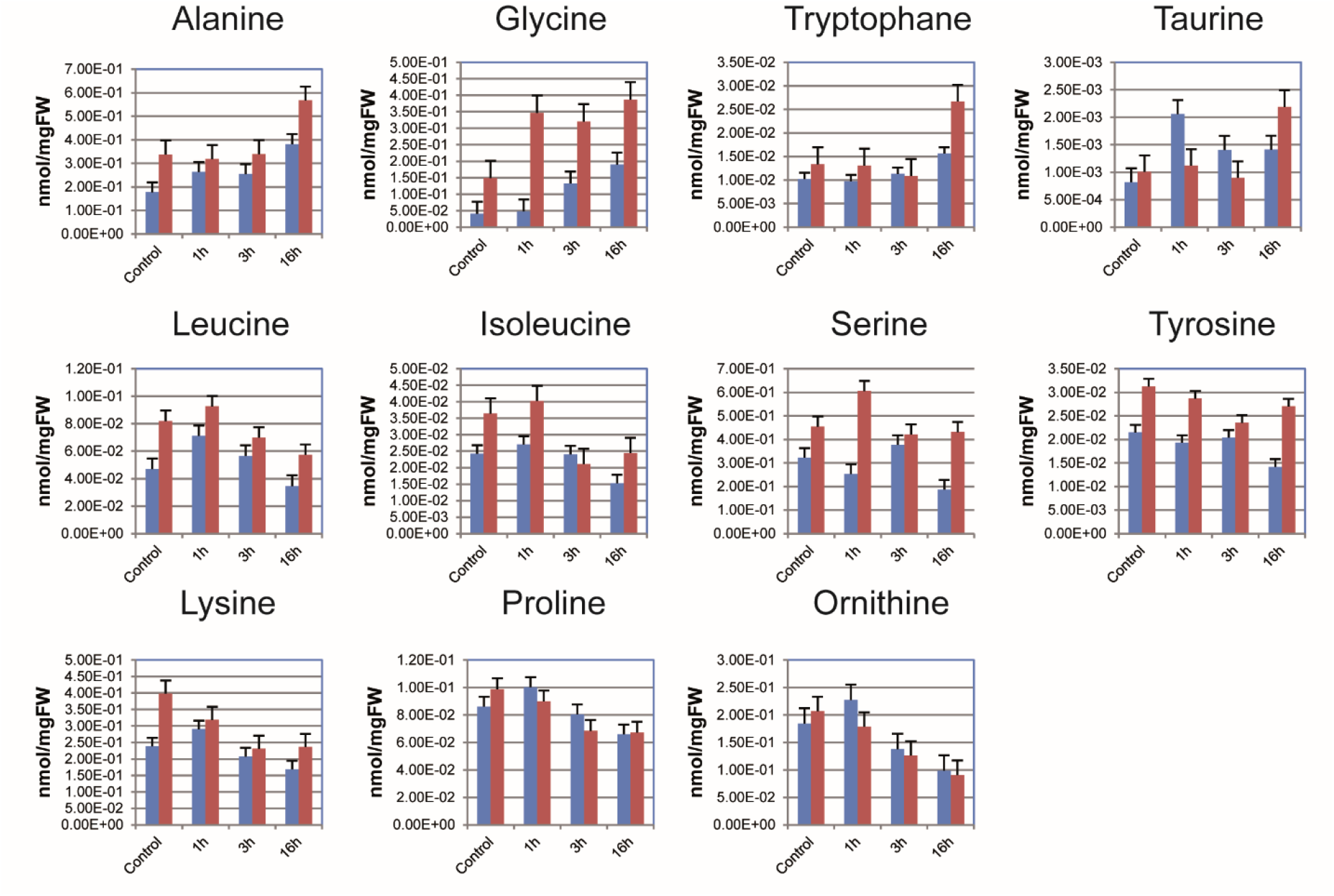
Absolute quantification of amino acids without and 1, 3 and 16 hours after flg22 exposure (1µM concentration in medium) in Col-0 Wt and *myc234* mutant backgrounds using GC-MS. Bars represent mean of three biological replicates; standard error is given.

## References

Agarwal, P., Kapoor, S., and Tyagi, A.K. (2011). Transcription factors regulating the progression of monocot and dicot seed development. Bioessays 33, 189–202.

Ahn, C.S., Cho, H.K., Lee, D.H., Sim, H.J., Kim, S.G., and Pai, H.S. (2016). Functional characterization of the ribosome biogenesis factors PES, BOP1, and WDR12 (PeBoW), and mechanisms of defective cell growth and proliferation caused by PeBoW deficiency in Arabidopsis. J Exp Bot 67, 5217–5232.

Baerenfaller, K., Grossmann, J., Grobei, M.A., Hull, R., Hirsch-Hoffmann, M., Yalovsky, S., Zimmermann, P., Grossniklaus, U., Gruissem, W., and Baginsky, S. (2008). Genome-scale proteomics reveals Arabidopsis thaliana gene models and proteome dynamics. Science 320, 938–941.

Bagwan, N., Bonzon-Kulichenko, E., Calvo, E., Lechuga-Vieco, A.V., Michalakopoulos, S., Trevisan-Herraz, M., Ezkurdia, I., Rodriguez, J.M., Magni, R., Latorre-Pellicer, A., Enriquez, J.A., and Vazquez, J. (2018). Comprehensive Quantification of the Modified Proteome Reveals Oxidative Heart Damage in Mitochondrial Heteroplasmy. Cell Rep 23, 3685–3697 e3684.

Bekker-Jensen, D.B., Kelstrup, C.D., Batth, T.S., Larsen, S.C., Haldrup, C., Bramsen, J.B., Sorensen, K.D., Hoyer, S., Orntoft, T.F., Andersen, C.L., Nielsen, M.L., and Olsen, J.V. (2017). An Optimized Shotgun Strategy for the Rapid Generation of Comprehensive Human Proteomes. Cell Syst 4, 587-+.

Bigeard, J., Colcombet, J., and Hirt, H. (2015). Signaling mechanisms in pattern-triggered immunity (PTI). Mol Plant 8, 521–539.

Bleecker, A.B., and Patterson, S.E. (1997). Last exit: senescence, abscission, and meristem arrest in Arabidopsis. Plant Cell 9, 1169–1179.

Boudsocq, M., Willmann, M.R., McCormack, M., Lee, H., Shan, L., He, P., Bush, J., Cheng, S.H., and Sheen, J. (2010). Differential innate immune signalling via Ca(2+) sensor protein kinases. Nature 464, 418–422.

Bowman, J.L., Smyth, D.R., and Meyerowitz, E.M. (1989). Genes directing flower development in Arabidopsis. Plant Cell 1, 37–52.

Breeze, E., Harrison, E., McHattie, S., Hughes, L., Hickman, R., Hill, C., Kiddle, S., Kim, Y.S., Penfold, C.A., Jenkins, D., Zhang, C., Morris, K., Jenner, C., Jackson, S., Thomas, B., Tabrett, A., Legaie, R., Moore, J.D., Wild, D.L., Ott, S., Rand, D., Beynon, J., Denby, K., Mead, A., and Buchanan-Wollaston, V. (2011). High-resolution temporal profiling of transcripts during Arabidopsis leaf senescence reveals a distinct chronology of processes and regulation. Plant Cell 23, 873–894.

Bu, Q., Jiang, H., Li, C.B., Zhai, Q., Zhang, J., Wu, X., Sun, J., Xie, Q., and Li, C. (2008). Role of the Arabidopsis thaliana NAC transcription factors ANAC019 and ANAC055 in regulating jasmonic acid-signaled defense responses. Cell Res 18, 756–767.

Burdiak, P., Rusaczonek, A., Witon, D., Glow, D., and Karpinski, S. (2015). Cysteine-rich receptor-like kinase CRK5 as a regulator of growth, development, and ultraviolet radiation responses in Arabidopsis thaliana. J Exp Bot 66, 3325–3337.

Byrne, M.E. (2009). A role for the ribosome in development. Trends Plant Sci 14, 512–519.

Caarls, L., Elberse, J., Awwanah, M., Ludwig, N.R., de Vries, M., Zeilmaker, T., Van Wees, S.C.M., Schuurink, R.C., and Van den Ackerveken, G. (2017). Arabidopsis JASMONATE-INDUCED OXYGENASES down-regulate plant immunity by hydroxylation and inactivation of the hormone jasmonic acid. Proc Natl Acad Sci U S A 114, 6388–6393.

Cao, F.Y., Yoshioka, K., and Desveaux, D. (2011). The roles of ABA in plant-pathogen interactions. J Plant Res 124, 489–499.

Chen, C.N., Chen, H.R., Yeh, S.Y., Vittore, G., and Ho, T.H. (2009). Autophagy is enhanced and floral development is impaired in AtHVA22d RNA interference Arabidopsis. Plant Physiol 149, 1679–1689.

Chi, H., Liu, C., Yang, H., Zeng, W.F., Wu, L., Zhou, W.J., Wang, R.M., Niu, X.N., Ding, Y.H., Zhang, Y., Wang, Z.W., Chen, Z.L., Sun, R.X., Liu, T., Tan, G.M., Dong, M.Q., Xu, P., Zhang, P.H., and He, S.M. (2018). Comprehensive identification of peptides in tandem mass spectra using an efficient open search engine. Nat Biotechnol.

Chick, J.M., Kolippakkam, D., Nusinow, D.P., Zhai, B., Rad, R., Huttlin, E.L., and Gygi, S.P. (2015). A mass-tolerant database search identifies a large proportion of unassigned spectra in shotgun proteomics as modified peptides. Nat Biotechnol 33, 743–749.

Cho, H.K., Ahn, C.S., Lee, H.S., Kim, J.K., and Pai, H.S. (2013). Pescadillo plays an essential role in plant cell growth and survival by modulating ribosome biogenesis. Plant J 76, 393–405.

Choi du, S., Lim, C.W., and Hwang, B.K. (2016). Proteomics and functional analyses of Arabidopsis nitrilases involved in the defense response to microbial pathogens. Planta 244, 449–465.

Christ, B., Sussenbacher, I., Moser, S., Bichsel, N., Egert, A., Muller, T., Krautler, B., and Hortensteiner, S. (2013). Cytochrome P450 CYP89A9 is involved in the formation of major chlorophyll catabolites during leaf senescence in Arabidopsis. Plant Cell 25, 1868–1880.

Clay, N.K., Adio, A.M., Denoux, C., Jander, G., and Ausubel, F.M. (2009). Glucosinolate metabolites required for an Arabidopsis innate immune response. Science 323, 95–101.

Dekker, C., Roe, S.M., McCormack, E.A., Beuron, F., Pearl, L.H., and Willison, K.R. (2011). The crystal structure of yeast CCT reveals intrinsic asymmetry of eukaryotic cytosolic chaperonins. EMBO J 30, 3078–3090.

Denoux, C., Galletti, R., Mammarella, N., Gopalan, S., Werck, D., De Lorenzo, G., Ferrari, S., Ausubel, F.M., and Dewdney, J. (2008). Activation of defense response pathways by OGs and Flg22 elicitors in Arabidopsis seedlings. Mol Plant 1, 423–445.

Duncan, O., Trosch, J., Fenske, R., Taylor, N.L., and Millar, A.H. (2017). Resource: Mapping the Triticum aestivum proteome. Plant J 89, 601–616.

Finkelstein, R., Reeves, W., Ariizumi, T., and Steber, C. (2008). Molecular aspects of seed dormancy. Annu Rev Plant Biol 59, 387–415.

Friml, J. (2003). Auxin transport - shaping the plant. Curr Opin Plant Biol 6, 7–12.

Fujita, M., Fujita, Y., Maruyama, K., Seki, M., Hiratsu, K., Ohme-Takagi, M., Tran, L.S., Yamaguchi-Shinozaki, K., and Shinozaki, K. (2004). A dehydration-induced NAC protein, RD26, is involved in a novel ABA-dependent stress-signaling pathway. Plant J 39, 863–876.

Futschik, M.E., and Carlisle, B. (2005). Noise-robust soft clustering of gene expression time-course data. J Bioinform Comput Biol 3, 965–988.

Gavin, A.C., Aloy, P., Grandi, P., Krause, R., Boesche, M., Marzioch, M., Rau, C., Jensen, L.J., Bastuck, S., Dumpelfeld, B., Edelmann, A., Heurtier, M.A., Hoffman, V., Hoefert, C., Klein, K., Hudak, M., Michon, A.M., Schelder, M., Schirle, M., Remor, M., Rudi, T., Hooper, S., Bauer, A., Bouwmeester, T., Casari, G., Drewes, G., Neubauer, G., Rick, J.M., Kuster, B., Bork, P., Russell, R.B., and Superti-Furga, G. (2006). Proteome survey reveals modularity of the yeast cell machinery. Nature 440, 631–636.

Gavin, A.C., Bosche, M., Krause, R., Grandi, P., Marzioch, M., Bauer, A., Schultz, J., Rick, J.M., Michon, A.M., Cruciat, C.M., Remor, M., Hofert, C., Schelder, M., Brajenovic, M., Ruffner, H., Merino, A., Klein, K., Hudak, M., Dickson, D., Rudi, T., Gnau, V., Bauch, A., Bastuck, S., Huhse, B., Leutwein, C., Heurtier, M.A., Copley, R.R., Edelmann, A., Querfurth, E., Rybin, V., Drewes, G., Raida, M., Bouwmeester, T., Bork, P., Seraphin, B., Kuster, B., Neubauer, G., and Superti-Furga, G. (2002). Functional organization of the yeast proteome by systematic analysis of protein complexes. Nature 415, 141–147.

Goda, H., Sasaki, E., Akiyama, K., Maruyama-Nakashita, A., Nakabayashi, K., Li, W., Ogawa, M., Yamauchi, Y., Preston, J., Aoki, K., Kiba, T., Takatsuto, S., Fujioka, S., Asami, T., Nakano, T., Kato, H., Mizuno, T., Sakakibara, H., Yamaguchi, S., Nambara, E., Kamiya, Y., Takahashi, H., Hirai, M.Y., Sakurai, T., Shinozaki, K., Saito, K., Yoshida, S., and Shimada, Y. (2008). The AtGenExpress hormone and chemical treatment data set: experimental design, data evaluation, model data analysis and data access. Plant J 55, 526–542.

Gohre, V., Jones, A.M., Sklenar, J., Robatzek, S., and Weber, A.P. (2012). Molecular crosstalk between PAMP-triggered immunity and photosynthesis. Mol Plant Microbe Interact 25, 1083–1092.

Grigorova, B., Mara, C., Hollender, C., Sijacic, P., Chen, X., and Liu, Z. (2011). LEUNIG and SEUSS co-repressors regulate miR172 expression in Arabidopsis flowers. Development 138, 2451–2456.

Guan, D., Yan, B., Thieme, C., Hua, J., Zhu, H., Boheler, K.R., Zhao, Z., Kragler, F., Xia, Y., and Zhang, S. (2017). PlaMoM: a comprehensive database compiles plant mobile macromolecules. Nucleic Acids Res 45, D1021–D1028.

Han, X., Kumar, D., Chen, H., Wu, S., and Kim, J.Y. (2014). Transcription factor-mediated cell-to-cell signalling in plants. J Exp Bot 65, 1737–1749.

Hauri, S., Comoglio, F., Seimiya, M., Gerstung, M., Glatter, T., Hansen, K., Aebersold, R., Paro, R., Gstaiger, M., and Beisel, C. (2016). A High-Density Map for Navigating the Human Polycomb Complexome. Cell Rep 17, 583–595.

Henras, A.K., Plisson-Chastang, C., O’Donohue, M.F., Chakraborty, A., and Gleizes, P.E. (2015). An overview of pre-ribosomal RNA processing in eukaryotes. Wiley Interdiscip Rev RNA 6, 225–242.

Henras, A.K., Soudet, J., Gerus, M., Lebaron, S., Caizergues-Ferrer, M., Mougin, A., and Henry, Y. (2008). The post-transcriptional steps of eukaryotic ribosome biogenesis. Cell Mol Life Sci 65, 2334–2359.

Hillmer, R.A., Tsuda, K., Rallapalli, G., Asai, S., Truman, W., Papke, M.D., Sakakibara, H., Jones, J.D.G., Myers, C.L., and Katagiri, F. (2017). The highly buffered Arabidopsis immune signaling network conceals the functions of its components. PLoS Genet 13, e1006639.

Hoehenwarter, W., Monchgesang, S., Neumann, S., Majovsky, P., Abel, S., and Muller, J. (2016). Comparative expression profiling reveals a role of the root apoplast in local phosphate response. BMC Plant Biol 16, 106.

Horstman, A., Fukuoka, H., Muino, J.M., Nitsch, L., Guo, C., Passarinho, P., Sanchez-Perez, G., Immink, R., Angenent, G., and Boutilier, K. (2015). AIL and HDG proteins act antagonistically to control cell proliferation. Development 142, 454–464.

Huang, D., Wu, W., Abrams, S.R., and Cutler, A.J. (2008). The relationship of drought-related gene expression in Arabidopsis thaliana to hormonal and environmental factors. J Exp Bot 59, 2991–3007.

Ishida, T., Maekawa, S., and Yanagisawa, S. (2016). The Pre-rRNA Processing Complex in Arabidopsis Includes Two WD40-Domain-Containing Proteins Encoded by Glucose-Inducible Genes and Plant-Specific Proteins. Mol Plant 9, 312–315.

Jang, G., Lee, S., Chang, S.H., Kim, J.K., and Choi, Y.D. (2018). Jasmonic acid modulates xylem development by controlling polar auxin transport in vascular tissues. Plant Biotechnol Rep 12, 265–271.

Jia, H.F., Chai, Y.M., Li, C.L., Lu, D., Luo, J.J., Qin, L., and Shen, Y.Y. (2011). Abscisic acid plays an important role in the regulation of strawberry fruit ripening. Plant Physiol 157, 188–199.

Kanei, M., Horiguchi, G., and Tsukaya, H. (2012). Stable establishment of cotyledon identity during embryogenesis in Arabidopsis by ANGUSTIFOLIA3 and HANABA TARANU. Development 139, 2436–2446.

Kong, A.T., Leprevost, F.V., Avtonomov, D.M., Mellacheruvu, D., and Nesvizhskii, A.I. (2017). MSFragger: ultrafast and comprehensive peptide identification in mass spectrometry-based proteomics. Nat Methods 14, 513–520.

Krogan, N.J., Cagney, G., Yu, H., Zhong, G., Guo, X., Ignatchenko, A., Li, J., Pu, S., Datta, N., Tikuisis, A.P., Punna, T., Peregrin-Alvarez, J.M., Shales, M., Zhang, X., Davey, M., Robinson, M.D., Paccanaro, A., Bray, J.E., Sheung, A., Beattie, B., Richards, D.P., Canadien, V., Lalev, A., Mena, F., Wong, P., Starostine, A., Canete, M.M., Vlasblom, J., Wu, S., Orsi, C., Collins, S.R., Chandran, S., Haw, R., Rilstone, J.J., Gandi, K., Thompson, N.J., Musso, G., St Onge, P., Ghanny, S., Lam, M.H., Butland, G., Altaf-Ul, A.M., Kanaya, S., Shilatifard, A., O’Shea, E., Weissman, J.S., Ingles, C.J., Hughes, T.R., Parkinson, J., Gerstein, M., Wodak, S.J., Emili, A., and Greenblatt, J.F. (2006). Global landscape of protein complexes in the yeast Saccharomyces cerevisiae. Nature 440, 637–643.

Kunkel, B.N., and Harper, C.P. (2018). The roles of auxin during interactions between bacterial plant pathogens and their hosts. J Exp Bot 69, 245–254.

Kustatscher, G., Grabowski, P., Schrader, T.A., Passmore, J.B., Schrader, M., and Rappsilber, J. (2019). Co-regulation map of the human proteome enables identification of protein functions. Nat Biotechnol 37, 1361–1371.

Lan, P., Li, W., and Schmidt, W. (2012). Complementary proteome and transcriptome profiling in phosphate-deficient Arabidopsis roots reveals multiple levels of gene regulation. Mol Cell Proteomics 11, 1156–1166.

Lee, J.H., Yoon, H.J., Terzaghi, W., Martinez, C., Dai, M., Li, J., Byun, M.O., and Deng, X.W. (2010). DWA1 and DWA2, two Arabidopsis DWD protein components of CUL4-based E3 ligases, act together as negative regulators in ABA signal transduction. Plant Cell 22, 1716–1732.

Lee, J.H., Terzaghi, W., Gusmaroli, G., Charron, J.B., Yoon, H.J., Chen, H., He, Y.J., Xiong, Y., and Deng, X.W. (2008). Characterization of Arabidopsis and rice DWD proteins and their roles as substrate receptors for CUL4-RING E3 ubiquitin ligases. Plant Cell 20, 152–167.

Leon-Reyes, A., Van der Does, D., De Lange, E.S., Delker, C., Wasternack, C., Van Wees, S.C., Ritsema, T., and Pieterse, C.M. (2010). Salicylate-mediated suppression of jasmonate-responsive gene expression in Arabidopsis is targeted downstream of the jasmonate biosynthesis pathway. Planta 232, 1423–1432.

Li, J., Brader, G., and Palva, E.T. (2004). The WRKY70 transcription factor: a node of convergence for jasmonate-mediated and salicylate-mediated signals in plant defense. Plant Cell 16, 319–331.

Liu, X., Li, Y., and Zhong, S. (2017). Interplay between Light and Plant Hormones in the Control of Arabidopsis Seedling Chlorophyll Biosynthesis. Front Plant Sci 8, 1433.

Lopez-Molina, L., Mongrand, S., Kinoshita, N., and Chua, N.H. (2003). AFP is a novel negative regulator of ABA signaling that promotes ABI5 protein degradation. Genes Dev 17, 410–418.

Lynch, T.J., Erickson, B.J., Miller, D.R., and Finkelstein, R.R. (2017). ABI5-binding proteins (AFPs) alter transcription of ABA-induced genes via a variety of interactions with chromatin modifiers. Plant Mol Biol 93, 403–418.

Meng, X., Zhou, J., Tang, J., Li, B., de Oliveira, M.V.V., Chai, J., He, P., and Shan, L. (2016). Ligand-Induced Receptor-like Kinase Complex Regulates Floral Organ Abscission in Arabidopsis. Cell Rep 14, 1330–1338.

Merchante, C., Stepanova, A.N., and Alonso, J.M. (2017). Translation regulation in plants: an interesting past, an exciting present and a promising future. Plant J 90, 628–653.

Meteignier, L.V., El Oirdi, M., Cohen, M., Barff, T., Matteau, D., Lucier, J.F., Rodrigue, S., Jacques, P.E., Yoshioka, K., and Moffett, P. (2017). Translatome analysis of an NB-LRR immune response identifies important contributors to plant immunity in Arabidopsis. J Exp Bot 68, 2333–2344.

Mine, A., Nobori, T., Salazar-Rondon, M.C., Winkelmuller, T.M., Anver, S., Becker, D., and Tsuda, K. (2017). An incoherent feed-forward loop mediates robustness and tunability in a plant immune network. EMBO Rep 18, 464–476.

Missbach, S., Weis, B.L., Martin, R., Simm, S., Bohnsack, M.T., and Schleiff, E. (2013). 40S ribosome biogenesis co-factors are essential for gametophyte and embryo development. PLoS One 8, e54084.

Mordret, E., Dahan, O., Asraf, O., Rak, R., Yehonadav, A., Barnabas, G.D., Cox, J., Geiger, T., Lindner, A.B., and Pilpel, Y. (2019). Systematic Detection of Amino Acid Substitutions in Proteomes Reveals Mechanistic Basis of Ribosome Errors and Selection for Translation Fidelity. Mol Cell 75, 427–441 e425.

Muller, T.M., Bottcher, C., and Glawischnig, E. (2019). Dissection of the network of indolic defence compounds in Arabidopsis thaliana by multiple mutant analysis. Phytochemistry 161, 11–20.

Nakashima, K., Takasaki, H., Mizoi, J., Shinozaki, K., and Yamaguchi-Shinozaki, K. (2012). NAC transcription factors in plant abiotic stress responses. Biochim Biophys Acta 1819, 97–103.

Navarro, L., Dunoyer, P., Jay, F., Arnold, B., Dharmasiri, N., Estelle, M., Voinnet, O., and Jones, J.D. (2006). A plant miRNA contributes to antibacterial resistance by repressing auxin signaling. Science 312, 436–439.

Nee, G., Kramer, K., Nakabayashi, K., Yuan, B., Xiang, Y., Miatton, E., Finkemeier, I., and Soppe, W.J.J. (2017). DELAY OF GERMINATION1 requires PP2C phosphatases of the ABA signalling pathway to control seed dormancy. Nat Commun 8, 72.

Nickstadt, A., Thomma, B.P.H.J., Feussner, I., Kangasjarvi, J., Zeier, J., Loeffler, C., Scheel, D., and Berger, S. (2004). The jasmonate-insensitive mutant jin1 shows increased resistance to biotrophic as well as necrotrophic pathogens. Mol Plant Pathol 5, 425–434.

Obayashi, T., Hayashi, S., Saeki, M., Ohta, H., and Kinoshita, K. (2009). ATTED-II provides coexpressed gene networks for Arabidopsis. Nucleic Acids Res 37, D987–991.

Paez Valencia, J., Goodman, K., and Otegui, M.S. (2016). Endocytosis and Endosomal Trafficking in Plants. Annu Rev Plant Biol 67, 309–335.

Patharkar, O.R., and Walker, J.C. (2016). Core Mechanisms Regulating Developmentally Timed and Environmentally Triggered Abscission. Plant Physiol 172, 510–520.

Patharkar, O.R., and Walker, J.C. (2018). Advances in abscission signaling. J Exp Bot 69, 733–740.

Patterson, S.E., and Bleecker, A.B. (2004). Ethylene-dependent and -independent processes associated with floral organ abscission in Arabidopsis. Plant Physiol 134, 194–203.

Pieterse, C.M., Van der Does, D., Zamioudis, C., Leon-Reyes, A., and Van Wees, S.C. (2012). Hormonal modulation of plant immunity. Annu Rev Cell Dev Biol 28, 489–521.

Ponnala, L., Wang, Y., Sun, Q., and van Wijk, K.J. (2014). Correlation of mRNA and protein abundance in the developing maize leaf. Plant J 78, 424–440.

Qi, L., Yan, J., Li, Y., Jiang, H., Sun, J., Chen, Q., Li, H., Chu, J., Yan, C., Sun, X., Yu, Y., Li, C., and Li, C. (2012). Arabidopsis thaliana plants differentially modulate auxin biosynthesis and transport during defense responses to the necrotrophic pathogen Alternaria brassicicola. New Phytol 195, 872–882.

Robatzek, S., Chinchilla, D., and Boller, T. (2006). Ligand-induced endocytosis of the pattern recognition receptor FLS2 in Arabidopsis. Genes Dev 20, 537–542.

Rogers, H., and Munne-Bosch, S. (2016). Production and Scavenging of Reactive Oxygen Species and Redox Signaling during Leaf and Flower Senescence: Similar But Different. Plant Physiol 171, 1560–1568.

Rounds, C.M., and Bezanilla, M. (2013). Growth mechanisms in tip-growing plant cells. Annu Rev Plant Biol 64, 243–265.

Schmid, M., Davison, T.S., Henz, S.R., Pape, U.J., Demar, M., Vingron, M., Scholkopf, B., Weigel, D., and Lohmann, J.U. (2005). A gene expression map of Arabidopsis thaliana development. Nat Genet 37, 501–506.

Sherp, A.M., Westfall, C.S., Alvarez, S., and Jez, J.M. (2018). Arabidopsis thaliana GH3.15 acyl acid amido synthetase has a highly specific substrate preference for the auxin precursor indole-3-butyric acid. J Biol Chem 293, 4277–4288.

Shim, J.S., Jung, C., Lee, S., Min, K., Lee, Y.W., Choi, Y., Lee, J.S., Song, J.T., Kim, J.K., and Choi, Y.D. (2013). AtMYB44 regulates WRKY70 expression and modulates antagonistic interaction between salicylic acid and jasmonic acid signaling. Plant J 73, 483–495.

Skinner, O.S., and Kelleher, N.L. (2015). Illuminating the dark matter of shotgun proteomics. Nat Biotechnol 33, 717–718.

Smyth, D.R., Bowman, J.L., and Meyerowitz, E.M. (1990). Early flower development in Arabidopsis. Plant Cell 2, 755–767.

Soltanieh, S., Lapensee, M., and Dragon, F. (2014). Nucleolar proteins Bfr2 and Enp2 interact with DEAD-box RNA helicase Dbp4 in two different complexes. Nucleic Acids Res 42, 3194–3206.

Song, G., Hsu, P.Y., and Walley, J.W. (2018). Assessment and Refinement of Sample Preparation Methods for Deep and Quantitative Plant Proteome Profiling. Proteomics 18, e1800220.

Song, S.K., Hofhuis, H., Lee, M.M., and Clark, S.E. (2008). Key divisions in the early Arabidopsis embryo require POL and PLL1 phosphatases to establish the root stem cell organizer and vascular axis. Dev Cell 15, 98–109.

Song, Y., Xiang, F., Zhang, G., Miao, Y., Miao, C., and Song, C.P. (2016). Abscisic Acid as an Internal Integrator of Multiple Physiological Processes Modulates Leaf Senescence Onset in Arabidopsis thaliana. Front Plant Sci 7, 181.

Spoel, S.H., Johnson, J.S., and Dong, X. (2007). Regulation of tradeoffs between plant defenses against pathogens with different lifestyles. Proc Natl Acad Sci U S A 104, 18842–18847.

Spoel, S.H., Koornneef, A., Claessens, S.M., Korzelius, J.P., Van Pelt, J.A., Mueller, M.J., Buchala, A.J., Metraux, J.P., Brown, R., Kazan, K., Van Loon, L.C., Dong, X., and Pieterse, C.M. (2003). NPR1 modulates cross-talk between salicylate- and jasmonate-dependent defense pathways through a novel function in the cytosol. Plant Cell 15, 760–770.

Stenvik, G.E., Butenko, M.A., Urbanowicz, B.R., Rose, J.K., and Aalen, R.B. (2006). Overexpression of INFLORESCENCE DEFICIENT IN ABSCISSION activates cell separation in vestigial abscission zones in Arabidopsis. Plant Cell 18, 1467–1476.

Su, J., Yang, L., Zhu, Q., Wu, H., He, Y., Liu, Y., Xu, J., Jiang, D., and Zhang, S. (2018). Active photosynthetic inhibition mediated by MPK3/MPK6 is critical to effector-triggered immunity. PLoS Biol 16, e2004122.

Sugawara, S., Hishiyama, S., Jikumaru, Y., Hanada, A., Nishimura, T., Koshiba, T., Zhao, Y., Kamiya, Y., and Kasahara, H. (2009). Biochemical analyses of indole-3-acetaldoxime-dependent auxin biosynthesis in Arabidopsis. Proc Natl Acad Sci U S A 106, 5430–5435.

Szymanski, J., Levin, Y., Savidor, A., Breitel, D., Chappell-Maor, L., Heinig, U., Topfer, N., and Aharoni, A. (2017). Label-free deep shotgun proteomics reveals protein dynamics during tomato fruit tissues development. Plant J 90, 396–417.

Tabassum, N., Eschen-Lippold, L., Athmer, B., Baruah, M., Brode, M., Maldonado-Bonilla, L.D., Hoehenwarter, W., Hause, G., Scheel, D., and Lee, J. (2019). Phosphorylation-dependent control of an RNA granule-localized protein that fine-tunes defence gene expression at a post-transcriptional level. Plant J.

Takasaki, H., Maruyama, K., Takahashi, F., Fujita, M., Yoshida, T., Nakashima, K., Myouga, F., Toyooka, K., Yamaguchi-Shinozaki, K., and Shinozaki, K. (2015). SNAC-As, stress-responsive NAC transcription factors, mediate ABA-inducible leaf senescence. Plant J 84, 1114–1123.

Teale, W.D., Paponov, I.A., and Palme, K. (2006). Auxin in action: signalling, transport and the control of plant growth and development. Nat Rev Mol Cell Biol 7, 847–859.

Thieme, C.J., Rojas-Triana, M., Stecyk, E., Schudoma, C., Zhang, W., Yang, L., Minambres, M., Walther, D., Schulze, W.X., Paz-Ares, J., Scheible, W.R., and Kragler, F. (2015). Endogenous Arabidopsis messenger RNAs transported to distant tissues. Nat Plants 1, 15025.

Tran, L.S., Nakashima, K., Sakuma, Y., Simpson, S.D., Fujita, Y., Maruyama, K., Fujita, M., Seki, M., Shinozaki, K., and Yamaguchi-Shinozaki, K. (2004). Isolation and functional analysis of Arabidopsis stress-inducible NAC transcription factors that bind to a drought-responsive cis-element in the early responsive to dehydration stress 1 promoter. Plant Cell 16, 2481–2498.

Truman, W.M., Bennett, M.H., Turnbull, C.G.N., and Grant, M.R. (2010a). Arabidopsis Auxin Mutants Are Compromised in Systemic Acquired Resistance and Exhibit Aberrant Accumulation of Various Indolic Compounds. Plant Physiology 152, 1562–1573.

Truman, W.M., Bennett, M.H., Turnbull, C.G., and Grant, M.R. (2010b). Arabidopsis auxin mutants are compromised in systemic acquired resistance and exhibit aberrant accumulation of various indolic compounds. Plant Physiol 152, 1562–1573.

Tsuda, K., Sato, M., Glazebrook, J., Cohen, J.D., and Katagiri, F. (2008). Interplay between MAMP-triggered and SA-mediated defense responses. Plant J 53, 763–775.

Van der Does, D., Leon-Reyes, A., Koornneef, A., Van Verk, M.C., Rodenburg, N., Pauwels, L., Goossens, A., Korbes, A.P., Memelink, J., Ritsema, T., Van Wees, S.C., and Pieterse, C.M. (2013). Salicylic acid suppresses jasmonic acid signaling downstream of SCFCOI1-JAZ by targeting GCC promoter motifs via transcription factor ORA59. Plant Cell 25, 744–761.

Walley, J.W., Sartor, R.C., Shen, Z., Schmitz, R.J., Wu, K.J., Urich, M.A., Nery, J.R., Smith, L.G., Schnable, J.C., Ecker, J.R., and Briggs, S.P. (2016). Integration of omic networks in a developmental atlas of maize. Science 353, 814–818.

Wan, C., Borgeson, B., Phanse, S., Tu, F., Drew, K., Clark, G., Xiong, X., Kagan, O., Kwan, J., Bezginov, A., Chessman, K., Pal, S., Cromar, G., Papoulas, O., Ni, Z., Boutz, D.R., Stoilova, S., Havugimana, P.C., Guo, X., Malty, R.H., Sarov, M., Greenblatt, J., Babu, M., Derry, W.B., Tillier, E.R., Wallingford, J.B., Parkinson, J., Marcotte, E.M., and Emili, A. (2015). Panorama of ancient metazoan macromolecular complexes. Nature 525, 339–344.

Wang, X., Gao, J., Zhu, Z., Dong, X., Wang, X., Ren, G., Zhou, X., and Kuai, B. (2015). TCP transcription factors are critical for the coordinated regulation of isochorismate synthase 1 expression in Arabidopsis thaliana. Plant J 82, 151–162.

Wang, Y., Li, L., Ye, T., Lu, Y., Chen, X., and Wu, Y. (2013). The inhibitory effect of ABA on floral transition is mediated by ABI5 in Arabidopsis. J Exp Bot 64, 675–684.

Waters, M.T., and Langdale, J.A. (2009). The making of a chloroplast. EMBO J 28, 2861–2873.

Weis, B.L., Kovacevic, J., Missbach, S., and Schleiff, E. (2015). Plant-Specific Features of Ribosome Biogenesis. Trends Plant Sci 20, 729–740.

Wisniewski, J.R., Hein, M.Y., Cox, J., and Mann, M. (2014). A “proteomic ruler” for protein copy number and concentration estimation without spike-in standards. Mol Cell Proteomics 13, 3497–3506.

Wojciechowska, N., Marzec-Schmidt, K., Kalemba, E.M., Zarzynska-Nowak, A., Jagodzinski, A.M., and Bagniewska-Zadworna, A. (2018). Autophagy counteracts instantaneous cell death during seasonal senescence of the fine roots and leaves in Populus trichocarpa. BMC Plant Biol 18, 260.

Wrzaczek, M., Brosche, M., Salojarvi, J., Kangasjarvi, S., Idanheimo, N., Mersmann, S., Robatzek, S., Karpinski, S., Karpinska, B., and Kangasjarvi, J. (2010). Transcriptional regulation of the CRK/DUF26 group of receptor-like protein kinases by ozone and plant hormones in Arabidopsis. BMC Plant Biol 10, 95.

Xu, G., Greene, G.H., Yoo, H., Liu, L., Marques, J., Motley, J., and Dong, X. (2017). Global translational reprogramming is a fundamental layer of immune regulation in plants. Nature 545, 487–490.

Yadeta, K.A., Elmore, J.M., Creer, A.Y., Feng, B., Franco, J.Y., Rufian, J.S., He, P., Phinney, B., and Coaker, G. (2017). A Cysteine-Rich Protein Kinase Associates with a Membrane Immune Complex and the Cysteine Residues Are Required for Cell Death. Plant Physiol 173, 771–787.

Ye, H., Liu, S., Tang, B., Chen, J., Xie, Z., Nolan, T.M., Jiang, H., Guo, H., Lin, H.Y., Li, L., Wang, Y., Tong, H., Zhang, M., Chu, C., Li, Z., Aluru, M., Aluru, S., Schnable, P.S., and Yin, Y. (2017). RD26 mediates crosstalk between drought and brassinosteroid signalling pathways. Nat Commun 8, 14573.

Yi, S.Y., Shirasu, K., Moon, J.S., Lee, S.G., and Kwon, S.Y. (2014). The activated SA and JA signaling pathways have an influence on flg22-triggered oxidative burst and callose deposition. PLoS One 9, e88951.

Zander, M., Thurow, C., and Gatz, C. (2014). TGA Transcription Factors Activate the Salicylic Acid-Suppressible Branch of the Ethylene-Induced Defense Program by Regulating ORA59 Expression. Plant Physiol 165, 1671–1683.

Zhang, M., Yuan, B., and Leng, P. (2009). The role of ABA in triggering ethylene biosynthesis and ripening of tomato fruit. J Exp Bot 60, 1579–1588.

Zheng, X.Y., Zhou, M., Yoo, H., Pruneda-Paz, J.L., Spivey, N.W., Kay, S.A., and Dong, X. (2015). Spatial and temporal regulation of biosynthesis of the plant immune signal salicylic acid. Proc Natl Acad Sci U S A 112, 9166–9173.

Zheng, X.Y., Spivey, N.W., Zeng, W., Liu, P.P., Fu, Z.Q., Klessig, D.F., He, S.Y., and Dong, X. (2012). Coronatine promotes Pseudomonas syringae virulence in plants by activating a signaling cascade that inhibits salicylic acid accumulation. Cell Host Microbe 11, 587–596.

Ziegler, J., Qwegwer, J., Schubert, M., Erickson, J.L., Schattat, M., Burstenbinder, K., Grubb, C.D., and Abel, S. (2014). Simultaneous analysis of apolar phytohormones and 1-aminocyclopropan-1-carboxylic acid by high performance liquid chromatography/electrospray negative ion tandem mass spectrometry via 9-fluorenylmethoxycarbonyl chloride derivatization. J Chromatogr A 1362, 102–109.

