## Supplementary material for "Reshaping of the *Arabidopsis thaliana* proteome landscape and co-regulation of proteins in development and immunity": Bassal Supplemental Methods and Data

### Bassal et al., Supplemental Methods and Data

#### Plant Material and Cultivation for Proteomics Experiments

*Arabidopsis thaliana* Col-0 accession was used for all experiments. Seeds were cold-stratified and sterilized with chlorine gas. Seeds were grown in liquid culture in ¼ Murashige & Skoog medium supplemented with Micronutrient Solution (Sigma-Aldrich, M0529) and Vitamin Solution (Sigma Aldrich, M3900) both to a concentration of 0.25x under long day conditions (16 h light, 8 h dark) at 22°C. For seedlings grown on plates, ¼ Murashige& Skoog medium was mixed with 1% plant agar. The mixture was autoclaved and 50 ml were poured onto a plate while hot. The agar was left to dry under a laminar flow hood. The sterilized seeds were mixed with 1 mL of ¼ Murashige& Skoog medium with 0.1% plant agar and then pipetted onto the agar plate in a line for uniform seedling growth. The plants were grown under long day conditions (16 h light, 8 h dark) at 22°C. Alternatively seeds were sown in steam sterilized soil. Germinated seedlings were transplanted into individual pots at the two cotyledon stage. Plants were grown in a growth chamber in one flat consisting of 60 pots under short day conditions (8 h light, 16 h dark) at 22°C with weekly watering. Plants were transferred to the green house upon flowering where they were grown until senescent in spring (April to May, approximately 15 h light). Plants were grown to various arbitrarily determined flowering stages and dissected into various tissues as well as treated with various chemicals, as described in Tables 1 and 2 and Supplemental Figure 1. Following harvest plants or dissected tissues were snap frozen in liquid nitrogen, ground to a fine powder and stored at -80°C.

#### Protein Extraction

Proteins were extracted from frozen, ground tissue with extraction buffer (EB)consisting of 50 mM Tris Base, 10 mM EDTA, 10 mM dithiothreitol (DTT), 4% SDS, 1.5% protease inhibitor cocktail for plants (Sigma-Aldrich, P9599) in dd$H_2O$ prepared fresh. The tissue was suspended in EB and the suspension agitated for 30 min at room temperature (RT). Following centrifugation at 16,000 g for 10 min at 10°C, the supernatant was ultra-centrifuged at 100,000 g for 1 h at 10 °C. The cleared supernatant was retained and protein concentration determined with the 2-D Quant kit (GE Healthcare) according to the manufacturer's instructions.

Different weight amounts of tissue with different volumetric amounts of EB were tested (Supplemental Figure 2). A 1:2 ratio of tissue to buffer was optimal, yielding both the highest concentrated protein extract in µg/µl of EB as well as the highest amount of protein extracted from tissue in µg/mg. There was a slight increase of both protein concentration and extracted protein amount when 500 mg of tissue were used as opposed to 250 mg but this was not significant. Proteins were also phenol extracted from 500 mg of tissue according to Majovsky et al. (Majovsky et al., 2014) to compare the SDS based method described here to a more conventional procedure finding wide spread application in plant proteomics studies. The total extracted protein was more than 3 times greater with the SDS based procedure described here. Subsequently 500 mg of tissue and 1000 ml EB were used for all experiments.

SDS-PAGE

SDS-PAGE according to Laemmli was used to fractionate the *Arabidopsis thaliana* proteome on the protein level. The gel lanes were cut into 5 bands resulting in 5 molecular weight fractions of the total protein extract (Supplemental Figure 3). Each band was further processed separately and the resulting peptide fractions also measured separately. The SDS based extraction led to substantially more protein identifications by LC-MS using a three hour gradient (details not given as this LC-MS method was only used for the purpose of optimization) in each band and in total as compared to the phenol extraction procedure even when 100 µg of phenol extracted proteins was loaded onto the gel lane as opposed to 40 µg of SDS extracted proteins, underscoring the quantitative results described above (Supplemental Figure 2).

A number of parameters were optimized. Three different migration lengths for protein in the gel matrix were tested: 1.5 cm, 2.5 cm and 3 cm. Allowing proteins to migrate precisely 2.5 cm into the gel led to the highest number of protein identifications in all bands except band 2 and in total (Supplemental Figure 2). Different amounts of protein extract, from 10 µg to 240 µg were loaded onto the gel lanes. Eighty µg proved to be the optimal amount leading to the highest number of protein identifications in each band as well as in total (Supplemental Figure 2). The optimal acrylamide concentration of the gel matrix was also assayed. There was essentially no difference in the amount of proteins identified in each band and in total between acrylamide concentrations of 12.5% and 16% (Supplemental Figure 2).

For all experiments, subsequently 80 µg of SDS extracted proteins were denatured in 1x SDS loading mix (66 mM Tris HCl, pH 6.8, 10% Glycerol, 0.01% (w/v) Bromphenol blue, 0.1 M DTT) for 5 min at 95°C. Proteins were loaded onto a gel consisting of a 5% stacking, 12.5% separating gel. Proteins were left to migrate at a potential of 80 V until the bromphenol blue front reached the stacking / separating gel interface. The potential was increased to 100 V and the proteins were left to migrate until the front reached a demarcation of exactly 2.5 cm inside the stacking gel. Proteins were fixed in the gel matrix, the gel was stained and destained with the Colloidal Blue Staining Kit (Thermo Fisher Scientific, LC6025) according to the manufacturer's instructions. Gel staining was allowed to proceed overnight at room temperature. The 2.5 cm gel lane was excised and divided into five bands (Supplemental Figure 3).

Protein In-gel Digestion with Trypsin

Each gel band was chopped into small 1mm$^3$ pieces. An appropriate volume of $ddH_2O$ was added to cover the gel pieces. The pieces were agitated for 10 min at 22°C at 850 rpm, subsequently the supernatant was discarded and the same procedure was repeated. An appropriate volume of 30% acetonitrile (ACN) in 100 mM $NH_4HCO_3$ was added for the duration and temperature as above. This treatment was repeated until the gel pieces were completely destained. The gel pieces were then washed with $ddH_2O$ and incubated with ACN for 15 min as above. The gel pieces were dried in a vacuum concentrator for 15 min.

For the reduction of cysteine bonds, an appropriate volume of 10 mM DTT in 100 mM $NH_4HCO_3$ was added to cover the dried gel pieces and agitated for 5 min at 850 rpm and 22°C. Then the temperature was increased to 50°C and agitation was prolonged for another 30 min. The supernatant was discarded and ACN was added followed by agitation for 15 min at 850 rpm at 22°C. The supernatant was discarded.

An appropriate volume of 54 mM iodoacetamide (IAA) in 100 mM $NH_4HCO_3$ was added to the gel pieces to alkylate free sulfhydryl groups and agitated for 15 min at 850 rpm and 22°C in darkness. The supernatant was discarded and the gel pieces were washed twice for 10 min each with 30% ACN in 100 mM $NH_4HCO_3$ The supernatant was discarded and the gel pieces were dried in a vacuum concentrator for 15 min.

An appropriate volume of 50 mM $NH_4HCO_3$, pH 8.5 in 5% ACN in $ddH_2O$ was added to the dried gel pieces. An appropriate volume of trypsin in digestion buffer (3 ng/ µl) was added to cover the gel pieces which were gently agitated at 37°C over night. An equal volume of 35% ACN, 0.4% trifluoroacetic acid (TFA) in $ddH_2O$ was added to the digest and agitated for 40 min at 22°C at 850 rpm. The supernatant was retained and the same volume of 35% ACN, 0.4% TFA in $ddH_2O$ was added and agitated for 15 min at 22°C at 850 rpm. The supernatants were combined and the peptides in solution were dried completely in a vacuum concentrator.

Desalting of in-gel digested proteins

Dried peptides were dissolved in 100 µl of 0.1% formic acid (FA). Stop and go extraction (STAGE)-tips were prepared by inserting six layers of Empore™ C18 solid phase immobilized on an inert PTFE matrix (3M) into 100 µl pipette tips using an in-house manufactured instrument. STAGE-tips were inserted into 2 ml reaction tubes using in-house-built adaptors and conditioned by applying 100 µL of 80% ACN, 0.1% (FA). STAGE-tips were centrifuged at 1,500 g for 2 min at room temperature. The flow-through was discarded and stage tips were equilibrated by applying 100 µl of 0.1% FA. The STAGE-tips were centrifuged at 1,500 g for 2 min. The flow-through was discarded and the procedure was repeated. One hundred µL of protein digest was added to the conditioned STAGE-tips and centrifuged at 1,500 g for 2 min. The flow-through was discarded. STAGE-tips were washed with 100 µl of 0.1% FA. After centrifugation, the flow-through was discarded and the same procedure was repeated. Following the final wash the stage tips were inserted into new 2 ml reaction tubes. For peptide elution, 50 µl of 80% ACN, 0.1% FA was added to the STAGE-tips. STAGE-tips were centrifuged at 1,500 g for 1 min, the eluate was retained and the procedure was repeated. The combined eluates were transferred to 1.5 mL reaction tubes and the peptides were dried completely in a vacuum concentrator and stored at -80°C.

Liquid chromatography and mass spectrometry (LC-MS) of in-gel digested proteins.

Desalted, dried peptides were dissolved in 10 µL of 5%ACN, 0.1% TFA and put in an ultrasonic bath for 6 minutes. The entirety of dried peptides extracted from a gel band was injected into an EASY-nLC1000 liquid chromatography machine connected to an Orbitrap Velos Pro mass spectrometer via an EASY Spray ion source (all from Thermo Fisher Scientific). Peptides were separated by way of reverse phase chemistry employing a C18 stationary phase in an Acclaim PepMap 100 pre-column (length 2cm, inner diameter 75 µm, particle diameter 3 µm) in-line with an EASY-Spray ES803 column (length 50 cm, inner diameter 75 µm, particle diameter 2 µm) (both from Thermo Fisher Scientific) and electrosprayed on-line. Mobile phase gradient elution was optimized using different gradients and flow rates (Supplementary table XX). A 9h gradient inclining from 5% to 40% ACN, 0.1% FA preceded by 55 minutes of isocratic flow of 5% ACN, 0.1%FA with a flow rate of 250 nl/min proved optimal (highest number of identified peptides and proteins, 0.1% peptide FDR, protein identification with at least 1 unique peptide).

Peptides were ionized in positive polarity using a capillary temperature of 275°C and a source voltage of 1.9 kV. The delta multipole offset was -7 V. Full MS survey scans in the Orbitrap mass analyzer were carried out with a resolution of 60,000 a mass range of 350-2000 m/z (this parameters was optimized; 350-1850 m/z was also tested) a maximum ion injection time (maxIT) of 500 ms and the automatic gain control (AGC) set to 1e06 acquiring a single microscan. Full MS spectra were internally calibrated on the fly using the lock mass option with the m/z 445.120024.Data dependent acquisition (DDA) of up to 20 MS/MS scans of selected ions were carried out in the linear ion trap (LTQ) mass analyzer. A minimum MS signal intensity threshold of 500 (1000 was also tested) was required to trigger an MS/MS scan. Selected ions were isolated with a window of 2 Da and fragmented with a normalized collision energy of 35%, a Q-value of 0.25 and an activation time of 10 ms. M/Z of fragmented ions were placed on an inclusion list with a repeat duration of 30 s, an exclusion duration of 240 s (180 s were also tested) and a parent mass window of 10 ppm. A single microscan was acquired and the AGC was set to 10e4 and the maxIT to 200 ms.

Identification and quantification of peptides and proteins.

Peptides were identified by matching MS/MS spectral data to *in silico* generated peptide fragment ion masses. The TAIR10 protein database (https://www.arabidopsis.org/download_files/Proteins/TAIR10_protein_lists/TAIR10_pep_20101214) with common contaminants amended (35394 sequences, 14486974 residues) was searched using both the Mascot search engine v2.5.1 (Matrix Science) linked to Proteome Discoverer v1.4 (Thermo Fisher) running on an in-house server and the Andromeda search engine in the MaxQuant distribution v.1.5.2.8 from the TU Munich (http://www.biochem.mpg.de/5111795/maxquant). For searches with Mascot the enzyme specificity was set to trypsin and two missed cleavages were tolerated. Carbamidomethylation of cysteine was set as a fixed modification and oxidation of methionine as a variable modification. The precursor ion m/z error tolerance was set to 7 ppm and the product ion m/z error tolerance to 0.8 Da. A decoy database search was performed to determine the peptide level false discovery rate (FDR).

Searches with Andromeda were performed with the enzyme specificity set to trypsin excluding cleavage before proline (P/trypsin) and two tolerated missed cleavages. Carbamidomethylation of cysteine was set as a fixed modification and oxidation of methionine and acetylation of protein N-termini as variable modifications. The precursor ion m/z error tolerance was set to 4.5 ppm and the product ion m/z error tolerance was set to 0.8 Da. A decoy database search was performed to determine the peptide and protein FDRs.

Search results from Mascot and Andromeda were combined and concatenated using the Scaffold software (Proteome Software). Peptide identifications were accepted if they could be established to achieve an FDR less than 1,0% by the Scaffold Local FDR algorithm. Protein identifications were accepted if they could be established to achieve an FDR less than 1,0% and contained at least 1 identified peptide. Protein probabilities were assigned by the Protein Prophet algorithm (Nesvizhskii et al., 2003). Proteins that contained similar peptides and could not be differentiated based on MS/MS analysis alone were grouped to satisfy the principles of parsimony. Proteins sharing significant peptide evidence were grouped into clusters (protein groups, henceforth referred to as proteins). The abundance of each identified protein in each sample was inferred by the protein quantification index (PQI) “Total Spectrum Count”, meaning the number of MS/MS spectra annotated with peptide sequences assigned to it.

Global analysis of protein post-translational modification was performed with the MSFragger database search engine featuring a GUI (Kong et al., 2017). MS .raw files were converted to mzML with the ProteoWizard file converter v 3.0.18142 and a concatenated target decoy database featuring reversed target sequences was created with a perl script available from Mascot. Default open search parameters were used with the exception of fragment mass tolerance set to 0.4 absolute mass units (ABS). Peptide and protein probabilities were assigned using the peptide and protein prophet algorithms linked to MSFragger via Philosopher v. 20180530.

Bioinformatics Analysis of the Data

*Arabidopsis thaliana* TAIR10 gene models and gene descriptions were downloaded from http://www.arabidopsis.org/download/index-auto.jsp?dir=%2Fdownload_files%2FGenes%2FTAIR10_genome_release. Microarray based gene expression profiling of *Arabidopsis thaliana* development dataset was downloaded from http://jsp.weigelworld.org/AtGenExpress/resources/. The total abundance of a protein in *Arabidopsis thaliana* was determined by summing the individual PQI values in the samples where it was quantified. PQI values for multiple gene models of the same locus, identified and quantified as distinct proteins, were summed to give a single PQI value for protein expression at the respective locus, where applicable. The same procedure was applied to cognate transcripts quantified in the microarray dataset. Log10 transcript and protein abundance were then mapped to their respective genetic loci (Figure 1). Gene ontology analysis of both measured and not measured (missing) nuclear genome encoded proteins was performed using the DAVID Bioinformatics Resources 6.8 (Huang da et al., 2009) using TAIR10 as background, a minimum category membership number (bin count) of 500 and considering a Benjamini corrected significance threshold α of 0.01. The ratio of bin counts to the entire measured and missing proteome for significant Biological Process, Cellular Compartment and Molecular Function categories is shown (Supplemental Figure 4).

Essentially two deep proteomics experiments were performed, I.) Tissue Development, II.) PAMP triggered immunity (PTI response). Two quantitative PQI matrices containing the protein identifiers in the rows, sample identifiers in the columns and the respective PQI values in the cells were produced one as an input for bioinformatics analysis for each experiment. Intersections and VENN plots of tissue specific proteins were calculated and produced using a tool featuring a web interface on line under the URL: http://bioinformatics.psb.ugent.be/webtools/Venn/ as well as with InteractiVenn (Heberle et al., 2015).

I. Tissue Development

The protein quantification matrix for this experiment contained PQI values quantifying protein expression at 15,601 nuclear, mitochondrial and chloroplastidic genetic loci (AGI codes) in 20 samples of different tissue at various stages of development (Table 1, Sample names in red). PQI values for multiple gene models of the same locus, identified and quantified as distinct proteins, were summed to give a single PQI value for protein expression at the respective locus, where applicable. The matrix was imported into the Perseus software v.1.5.3.2 from the TU Munich (http://www.biochem.mpg.de/5111810/perseus).

Principal component analysis (PCA)

For a first general overview of the data, i.e. the samples and their relationship to one another, the matrix column vectors were converted to unit vector norm. This projected the samples onto a unit hypersphere to correct for possible inequalities of a primarily technical nature such as unequal sample loading or LC-MS response. It was followed by PCA, linearly mapping the samples into an optimal two dimensional sub-space of the original high dimensional space. 20 quantiles were determined for the loadings of principal components 1 and 2. The proteins corresponding to the 1$^{st}$ and 20$^{th}$ quantile of both components were extracted and imported into MapMan v3.0.0 hosted by the Forschungszentrum Jülich (https://mapman.gabipd.org/home) , using the respective loadings themselves as numeric values. To categorize their functions, these proteins were mapped to the "Overview" pathways available in the software with the Arabidopsis Affymetrx TAIR10 mapping (Ath_AFFY_ATH1_TAIR10_Aug2012) downloaded from the MapMan store (https://mapman.gabipd.org/mapmanstore). AGI codes of proteins in functional categories (bins) to which mapping was considered statistically significant by way of a Benjamini-Hochberg corrected Wilcoxon Rank Sum test at a significance level $\alpha$ of 5% were retained.

Hierarchical cluster analysis (HCL)

Row vectors (proteins) with less than 2 PQI entries were deleted from the "Tissue Matrix" reflecting the minimum number of 2 samples per tissue in roots, to decrease stochastic noise, reducing the matrix to 12,810 rows. Subsequently the column vectors were converted to unit vector norm and the row vectors were z-score transformed resulting in a mean PQI value of 0 and unit variance for every protein in the 20 tissue samples. Because we sampled several tissues at several points in the plant's lifetime we can consider changes in a protein's abundance to be relative to its overall mean protein abundance in *Arabidopsis* using this transformation. HCL was done using Spearman correlation as distance to cluster proteins (rows) and Pearson correlation to cluster samples (columns). Pre-processing with 300 rounds of k-means clustering was performed.

Fuzzy c-means cluster analysis

The columns in the z-score transformed matrix used for HCL were ordered according to the HCL column dendrogram as well as according to first tissues and then plant age to produce two matrices which were both exported. C-means clustering which is a type of soft partitioning where variables can be assigned to more than 1 cluster with a membership value between 0 and 1 quantifying the quality of fit to the respective cluster centers was performed using the Mfuzz R package downloaded from Bioconductor (https://bioconductor.org/packages/release/bioc/html/Mfuzz.html) in R Studio 1.0.136 with R v3.4. The optimal number of clusters and fuzzifier m were determined using the cselection and partcoef functions in the package (number of clusters = 16, m =1.3) to lead to uniform clustering of the randomized input matrix, i.e. no clustering of random data. This was done 1000 times to calculate a permutation based FDR for the clustering of the input data. Gene ontology of cluster members was done with the DAVID

Bioinformatics Resources 6.8 using the 12,810 input loci as background, a minimum category membership number (bin count) of 20 and considering a Benjamini corrected significance threshold α of 0.01. To further evaluate the possible biological role of the proteins in significant functional categories, their AGI codes were used to query the STRING database for physical interaction setting the stringency to highest confidence interactions, which we have shown to be true positive previously (Hoehenwarter et al., 2013), using experiments, databases, co-occurrence and co-expression as interaction sources and showing only interactions between proteins in the input set. For analysis of cluster 4 interactions only experimental interaction sources were allowed to ensure the resulting networks displayed only *bona fide* physical protein interactions meaning proteins in complex. Ninety five percent (95%) confidence intervals were determined of the raw spectral counts in all of the 20 different tissue samples for the clustered proteins to assess significance of individual values in the entire distribution. The ratio of significant (individual PQI value outside of confidence interval) PQI values in tissue samples where clustered proteins showed clear increases or decreases in abundance to significant PQI values in all samples was calculated and plotted for RBF protein complexes and all ribosomal proteins in the Tissue Development matrix.

II. PTI response

The protein quantification matrix for this experiment contained PQI values for 8,344 protein groups quantified in measurements of 7 and 10 day old seedlings grown in liquid culture mock treated with $H_2O$ or treated with 1µM flg22 for 16 hours (Table 1, Sample names in green). The sample PQI values (column vectors) were scaled to unit vector norm. Proteins that were quantified in less than half of all or in half of the samples not specific to treatment and control groups were removed leaving 7092 quantified proteins. Protein PQI values were z-score transformed. Hierarchical clustering of the proteins was done with maximum distance, average linkage and preprocessing with k-means clustering. Samples were clustered in the same manner employing euclidean distance. Proteins were partitioned into six clusters. The cluster termed 7080 contained 1774 proteins whose abundance increased upon seedling exposure to flg22, the cluster termed 7082 contained 915 proteins whose abundance decreased accordingly (Supplemental Table 15).

The MapMan software was used to assign the flg22 responsive proteins to functional categories in and map them to a model of PAMP triggered immunity. The *Arabidopsis* Affymetrix TAIR10 mapping was downloaded from the MapMan store. The Arabidopsis GO slim ontology (ATH_GO_GOSLIM.txt) and the TAIR GO slim categories (TAIR_GO_slim_categories.txt) were downloaded from https://www.arabidopsis.org/download/index-auto.jsp?dir=%2Fdownload_files%2FGO_and_PO_Annotations%2FGene_Ontology_Annotations. A plant PAMP triggered immunity pathway was custom drawn and uploaded into the MapMan software. The respective mapping was custom constructed by supplementing bins from the MapMAn Affymetrix mapping depicted on the pathway and related to PTI with corresponding GO terms from the TAIR GO slim ontology, thus producing a mapping file that draws from both sources (Supplemental Table 16). The flg22 responsive proteins from both clusters (increasing and decreasing in abundance) were mapped to the pathway. The sum of their z-scores in both flg22 treated samples was used as a proxy for changes in protein abundance (positive if abundance increased, negative if abundance decreased following flg22 exposure). Transcription factor / co-activator binding sites were determined with AGRIS.

Protein Extraction and FASP for Targeted Proteomics

Seedling were ground in liquid nitrogen. 1 mL EB was added to 500 mg ground tissue and vortexed for 30 secs and then kept in a thermomixer at 600 rpm at 95°C for 10 min. Samples were vortexed again for 30 secs and then returned to a thermomixer at 600 rpm at 23°C for 20 min. The extracts were centrifuged 16,000 g for 10 min at 10°C. The supernatants were transferred to new tubes and again centrifuged at 20,000 g for 30 min at 10°C. The supernatants were retained and protein concentration was determined by 2D-Quant. Three pools of seedlings were grown for each biological condition ("untreated" (C) and "flg 22 supplemented" (F)) and analyzed further (three biological replicates).

Amicon Ultra 30K filters were kept in 5% Tween shaking overnight at 60 rpm. Filters were washed with $ddH_2O$ by shaking at 60 rpm for 30 minutes, this was repeated twice. Filter units were assembled and 100 µg of proteins in solution were volume adjusted to 200 µL with UA (50 mM Tris Base, 8 M Urea, pH 8.0) and then added each to filter units. Then it was centrifuged at 16,100 g for 10 min at room temperature (RT), the flow through was discarded. The filter units were washed with 200 µL UA three times by centrifugation at 16,100 g for 10 min at RT, the flow through was discarded. 100 µL of UAD (50 mM Tris Base, 8 M Urea, 100 mM DTT, pH 8.0) was added to each sample and incubated in a thermomixer at 600 rpm for 1 hr at 22°C, then samples were centrifuged at 16,100 g for 10 min at RT, the flow through was discarded. 100 µL of UAI (50mM Tris Base, 8M Urea, 50 mM IAA, pH 8.0) was added, samples were mixed in a thermomixer at 600 rpm for 1 hr at 22°C and centrifuged at 16,100 g for 10 min at RT, the flow through was discarded. After that, filter units were washed three times with 200 µL UA by centrifugation at 16,100 g for 10 min at RT, the flow through was discarded. 100µL of ABC (50 mM $NH_4HCO_3$, pH 8.0) was added to each sample and then centrifuged at 16,100 g for 10 min at RT, this was repeated twice. Then 50 µL of ABC with 10 µL Lys-C (0.05 µg/µL) were added to filter units and samples were incubated in a thermomixer at 600 rpm for 4 hrs at 37°C. Afterwards, 10 µL (0.2 µg/µL) Trypsin was added to each sample and then kept in thermomixer overnight at 600 rpm at 37°CFilters were transferred to new collection tubes and then centrifuged at 16,100 g for 10 min at RT, the flow through was kept. 40 µL ABC was added to each filter and then centrifuged again at 16,100 g for 10 min at RT, the flow through was kept, this step was repeated once. Then samples were dried in a vacuum concentrator. Digested peptides were desalted as above.

PRM analysis

Proteotyipc peptides for target proteins were selected by first measuring the sample using a conventional DDA scan strategy. The up to three identified peptides with the highest #PSMs unique to target proteins (mapping exclusively to target protein master protein groups in Proteome Discoverer 2.1 with unique m/z) were used to populate a peptide target list for Parallel Reaction Monitoring (PRM) analysis of all 99 target proteins. Target proteins lacking three proteotypic peptides at this stage were subjected to d::pPop analysis for *in silico* determination of proteotypic peptides with favorable ESI response and corresponding m/z. Samples were then analyzed with a TDA scan strategy targeting *in silico* determined m/z followed by non-retention time (RT) scheduled PRM of any remaining m/z not leading to proteotypic peptide identification in the earlier two MS measurements (DDA and TDA). The results from the three iterative MS scan strategies allowed construction of a PRM target list that contained at least on measured proteotypic peptide m/z for every target protein plus corresponding RTs for RT scheduling. A RT window of +/- 7 min was placed around measured RTs for retention time scheduling of the 243 peptides in quantitative PRM analysis.

Dried peptides were dissolved in 5% acetonitrile, 0.1% trifluoric acid, and injected into an EASY-nLC 1000 liquid chromatography system (Thermo Fisher Scientific). Peptides were separated using liquid chromatography C18 reverse phase chemistry employing a 180 min gradient increasing from 5% to 40% acetonitrile in 0.1% FA, and a flow rate of 250 nL/min. Eluted peptides were electrosprayed on-line into a QExactive Plus mass spectrometer (Thermo Fisher Scientific). The spray voltage was 1.9 kV, the capillary temperature 275°C and the Z-Lens voltage 240 V. A full MS survey scan was carried out with chromatographic peak width set to 15 s, resolution 35,000, automatic gain control (AGC) 1E+06 and a max injection time (IT) of 100 ms. The full scan was followed by RT scheduled PRM scanning without multiplexing with HCD fragmentation. MS/MS scans were acquired with resolution 17,500, AGC 2E+05, IT 100 ms, loop count 10, isolation width 1.6 m/z, isolation offset 0.5 and a normalized collision energy 27. Each of the three samples for each condition was injected three times (three technical replicates per biological replicate).

Peptides and proteins were identified using the Mascot software v2.5.0 (Matrix Science) linked to Proteome Discoverer v2.1 (Thermo Fisher Scientific).The enzyme was set to trypsin. A precursor ion mass error of 5 ppm and a fragment ion mass error of 0.02 Da were tolerated in searches of the TAIR10 database amended with common contaminants (35934 sequences, 14486974 residues). Carbamidomethylation of cysteine was set as a fixed modification and oxidation of methionine (M) tolerated as a variable modifications. A PSM, peptide and protein level false discovery rate (FDR) was calculated for all identified spectra and peptides and proteins based on the target-decoy database model. The significance threshold α was set at 0.01 to accept PSM, peptide and protein identifications.

#### Data Analysis

Quantitative analysis of PRM data was done with the Skyline software v.4.2.0 (Pino et al., 2017). A spectral library was created using the DDA/TDA measurements described above. All target protein primary structures were concatenated in a FASTA file and imported. Raw PRM data was imported with the following "Transition Settings": Filter tab: "Precursor charges" were set to 2,3, "Ion charges" were set to 1,2 and "Ion types" were set to y,b,p, "Product ion selection" was set from m/z > precursor to 6 ions. Library tab: "Pick" was set to 6 product ions. Instrument tab: "Method match tolerance m/z" kept at a default of 0.055. Full-Scan tab: "Isotope peaks included" set to Count, "precursor mass analyzer" set to Orbitrap, "Peaks" set to 3, "Resolving power" set to 30,000 at 400 m/z. Under MS/MS filtering, the "Acquisition method" was set to Targeted, "product mass analyzer" set to Orbitrap, "resolving power" set to 17,500 at 400 m/z. In retention time filtering, "Include all matching scans" was selected. The sum of six picked product ion signal peak areas was extracted as PQI for each target peptide.

The matrix of target peptide PQI values in all measured samples was imported into the Perseus software v.1.6.6.0 (Tyanova et al., 2016). PQI vales were log2(x) transformed, and grouped by categories (C and F), each category consisting of three measurements of each of the three seedling pools so 9 total measurements. A permutation based multiples testing corrected two sample t-test was used to assess the significance of changes in peptide abundance between the conditions using a significance threshold $\alpha$ of 5% (q-value < 0.05). For each peptide, its median untransformed PQI in the three replicate sample measurements was used as an estimate of its abundance and used to calculate fold changes of abundance for each of the three samples per condition (i.e. F1/C1, F2/C2, F3/C3). Outliers in peptide fold changes were removed manually. Median peptide fold changes for each protein were used to infer fold changes in protein abundance in the three samples per condition, respectively. Protein fold changes were log2(x) transformed and the median and the standard error were calculated. Protein fold changes were considered significant if at least one of the peptides used for inference showed a statistically significant change in PQI between the conditions as determined above.

Measurement of Hormones and Metabolites

Phytohormone concentrations were determined from 50 mg of tissue fresh weight. Sample processing, data acquisition, instrumental setup, and quantifications (using 2 ng of ABA-D6, 5 ng IAA-D5, 5 ng JA-D2, 0,75 ng JA-Ile-D2, 30 ng OPDA-D5, 45 ng ACC-D4, and 1.5 ng SA-D4 as internal standards per sample) were performed as described (Ziegler et al., 2014).

Table 1

| Sample name | Description |
|---|---|
| **FF66** | Flowers from the 1st developmental flowering stage, grown for 66 days under short day conditions |
| **FF73** | Flowers from the 2nd developmental flowering stage, grown for 73 days under short day conditions |
| **FF90** | Green siliques from the 3rd developmental flowering stage, grown for 90 days under short day conditions |
| **FF93** | Brown siliques from the 4th developmental flowering stage, grown for 93 days under short day conditions |
| **SF66** | Inflorescence from the 1st developmental flowering stage, grown for 66 days under short day conditions |
| **SF73** | Inflorescence from the 2nd developmental flowering stage, grown for 73 days under short day condition |
| **SF90** | Inflorescence from the 3rd developmental flowering stage, grown for 90 days under short day conditions |
| **R7** | Roots grown in agar medium for seven days under long day conditions |
| **R10** | Roots grown in agar medium for ten days under long day conditions |
| **CLF66** | Juvenile Leaves from the 1st developmental flowering stage, grown for 66 days under short day conditions |
| **CLF73** | Juvenile Leaves from the 2nd developmental flowering stage, grown for 73 days under short day conditions |
| **CLF90** | Juvenile Leaves from the 3rd developmental flowering stage, grown for 90 days under short day conditions |
| **LF66** | Rosette Leaves from the first developmental flowering stage, grown for 66 under short day conditions |
| **LF73** | Rosette Leaves from the second developmental flowering stage, grown for 73 days under short day conditions |
| **LF90** | Rosette Leaves from the third developmental flowering stage, grown for 90 days under short day conditions |
| **LF40** | Plant leaves grown in soil under short day condition for 40 days without flg22 treatment |
| **LF40 + flg22** | Plant leaves grown in soil under short day condition for 40 days with flg22 treatment (plants sprayed with 1µM in the morning) under short day conditions |
| **C7** | Plant grown in liquid culture for seven days under short day condition |
| **C7 + flg22** | Plant grown in liquid culture for seven days with flg22 treatment (1 µM growth medium concentration for 16 hours) under long day conditions |
| **C10** | Plant grown in liquid culture for ten days under long day conditions |
| **C10 + flg22** | Plant grown in liquid culture for ten days with flg22 treatment (1 µM growth medium for 16 hours) under long day conditions |
| **LF7** | Leaves grown in agar medium for seven days under long day conditions |
| **LF10** | Leaves grown in agar medium for ten days under long day conditions |

Table 2

| Plant Age in Days for Sampling | Description |
|---|---|
| **7** | Seedling |
| **10** | Seedling |
| **40** | Mature rosette leaves before bolting |
| **66** | Mature plant, opening of first flower buds (Supplemental Figure 1) |
| **73** | Mature flowering plant, flower buds fully opened (Supplemental Figure 1) |
| **90** | Mature flowering plant, appearance of first green siliques, rosette leaves beginning purple color |
| **93** | Senescent plant, appearance of brown siliques |

Table 3

| Sample name | Protein numbers | Total unique peptide number | Total unique spectral number | Total MS2 |
|---|---|---|---|---|
| **Leaves (LF)** | | | | |
| **LF7** | 6371 | 36552 | 42849 | 442592 |
| **LF10** | 6298 | 34054 | 40081 | 458049 |
| **LF40** | 6056 | 36654 | 44076 | 554763 |
| **LF66** | 7096 | 46589 | 59867 | 524479 |
| **LF73** | 6862 | 46172 | 57580 | 472643 |
| **LF90** | 7277 | 47813 | 59387 | 544959 |
| **Cauline Leaves (CLF)** | | | | |
| **CLF66** | 7606 | 47813 | 66047 | 493577 |
| **CLF73** | 6252 | 39271 | 49935 | 496702 |
| **CLF90** | 7505 | 50676 | 65384 | 480931 |
| **Roots (R)** | | | | |
| **R7** | 7533 | 48551 | 63728 | 515761 |
| **R10** | 8525 | 54293 | 72986 | 548599 |
| **Stem (SF)** | | | | |
| **SF66** | 9104 | 67083 | 86517 | 516562 |
| **SF73** | 8531 | 57965 | 75329 | 505263 |
| **SF90** | 7978 | 50417 | 63898 | 494507 |
| **Flowers (FF)** | | | | |
| **FF66** | 9295 | 64241 | 83136 | 518934 |
| **FF73** | 8913 | 58341 | 74704 | 525789 |
| **Siliques (FF)** | | | | |
| **FF90** | 9524 | 63345 | 80116 | 501454 |
| **FF93** | 7481 | 44409 | 54374 | 414514 |
| **Other** | | | | |
| **C7** | 6697 | 36961 | 43652 | 459995 |
| **C7 + flg22** | 6949 | 41687 | 49480 | 485253 |
| **C10** | 7044 | 40487 | 50442 | 589299 |
| **C10 + flg22** | 7297 | 42532 | 52455 | 561851 |
| **LF40 + flg22** | 5977 | 35421 | 42771 | 498141 |
| **Total** | 15790 | | | 11604617 |

References

**Heberle, H., Meirelles, G.V., da Silva, F.R., Telles, G.P., and Minghim, R.** (2015). InteractiVenn: a web-based tool for the analysis of sets through Venn diagrams. BMC Bioinformatics **16,** 169.

**Hoehenwarter, W., Thomas, M., Nukarinen, E., Egelhofer, V., Rohrig, H., Weckwerth, W., Conrath, U., and Beckers, G.J.** (2013). Identification of novel in vivo MAP kinase substrates in Arabidopsis thaliana through use of tandem metal oxide affinity chromatography. Mol Cell Proteomics **12,** 369-380.

**Huang da, W., Sherman, B.T., and Lempicki, R.A.** (2009). Bioinformatics enrichment tools: paths toward the comprehensive functional analysis of large gene lists. Nucleic Acids Res **37,** 1-13.

**Kong, A.T., Leprevost, F.V., Avtonomov, D.M., Mellacheruvu, D., and Nesvizhskii, A.I.** (2017). MSFragger: ultrafast and comprehensive peptide identification in mass spectrometry-based proteomics. Nat Methods **14,** 513-520.

**Majovsky, P., Naumann, C., Lee, C.W., Lassowskat, I., Trujillo, M., Dissmeyer, N., and Hoehenwarter, W.** (2014). Targeted proteomics analysis of protein degradation in plant signaling on an LTQ-Orbitrap mass spectrometer. J Proteome Res **13,** 4246-4258.

**Nesvizhskii, A.I., Keller, A., Kolker, E., and Aebersold, R.** (2003). A statistical model for identifying proteins by tandem mass spectrometry. Anal Chem **75,** 4646-4658.

**Pino, L.K., Searle, B.C., Bollinger, J.G., Nunn, B., MacLean, B., and MacCoss, M.J.** (2017). The Skyline ecosystem: Informatics for quantitative mass spectrometry proteomics. Mass Spectrom Rev.

**Tyanova, S., Temu, T., Sinitcyn, P., Carlson, A., Hein, M.Y., Geiger, T., Mann, M., and Cox, J.** (2016). The Perseus computational platform for comprehensive analysis of (prote)omics data. Nat Methods **13,** 731-740.

**Ziegler, J., Qwegwer, J., Schubert, M., Erickson, J.L., Schattat, M., Burstenbinder, K., Grubb, C.D., and Abel, S.** (2014). Simultaneous analysis of apolar phytohormones and 1-aminocyclopropan-1-carboxylic acid by high performance liquid chromatography/electrospray negative ion tandem mass spectrometry via 9-fluorenylmethoxycarbonyl chloride derivatization. J Chromatogr A **1362,** 102-109.
